# Cardiopharyngeal deconstruction and ancestral tunicate sessility

**DOI:** 10.1101/2021.02.10.430586

**Authors:** A. Ferrández-Roldán, M. Fabregà-Torrus, G. Sánchez-Serna, E. Durán-Bello, M. Joaquín-Lluís, J. Garcia-Fernàndez, R. Albalat, C. Cañestro

## Abstract

A key problem in understanding chordate evolution has been the origin of sessility of ascidians, and whether the appendicularian free-living style represents a primitive or derived condition of tunicates. To address this problem, we performed comprehensive developmental and genomic comparative analyses of the cardiopharyngeal gene regulatory network (GRN) between appendicularians and ascidians. Our results reveal that the cardiopharyngeal GRN has suffered a process of evolutionary deconstruction with massive ancestral losses of genes (*Mesp*, *Ets1/2*, *Gata4/5/6*, *Mek1/2*, *Tbx1/10*, and RA- and FGF-signaling related genes) and subfunctions (e.g. *FoxF*, *Islet*, *Ebf*, *Mrf*, *Dach* and Bmp signaling). These losses have led to the deconstruction of two modules of the cardiopharyngeal GRN that in ascidians are related to early and late multipotent state cells involved in lineage fate determination towards first and secondary heart fields, and siphon muscle. Our results allow us to propose an evolutionary scenario, in which the evolutionary deconstruction of the cardiopharyngeal GRN has had an adaptive impact on the acceleration of the developmental cardiac program, the redesign of the cardiac architecture into an open-wide laminar structure, and the loss of pharyngeal muscle. Our findings, therefore, provide evidence supporting that the ancestral tunicate had a sessile ascidian-like lifestyle, and points to the deconstruction of the cardiopharyngeal GRN in appendicularians as a key event that facilitated the evolution of their pelagic free-living style connected to the innovation of the house.

## 1 Introduction

A key problem in the field of Evolutionary Developmental Biology (EvoDevo) is understanding the origin and radiation of our own phylum, the chordates^1,2^. The discovery that tunicates (a.k.a. urochordates) are the sister group of vertebrates, and therefore that the branching of cephalochordates is basal within chordates^3,4^, has provided a novel view of the last common ancestor of chordates as a free-living organism, in contrast to the traditional view proposed by Garstang (1928) in which it had a sessile ascidian-like adult lifestyle^5^. This novel view has brought renewed interest in appendicularians, whose complete free-living style could parsimoniously represent the ancestral condition of tunicates^6–8^ considering their most accepted position as the sister group of the remaining tunicates^8–11^. In such evolutionary scenario, sessility in tunicates would be a derived ascidian trait that could have evolved during the acquisition of their dramatic larval to adult metamorphosis, an event that is absent in appendicularians^8,12^. Certain developmental features of appendicularians that resemble Aplousobranchia ascidians, however, have been argued to represent traces of sessility, which therefore would favor the hypothesis that appendicularians have evolved from a tunicate ancestor with a sessile adult lifestyle and a larval dispersal stage similar to ascidians^13–15^. The fossil record reveals that the body plans of appendicularians and ascidians were already differentiated in the Early Cambrian, suggesting that the divergence between the two groups is ancient^16,17^. Thus, the problem of the lifestyle of the last common ancestor of tunicates remains unsolved.

The evolution of the cardiopharyngeal gene regulatory network appears to be a pivotal aspect to understand the evolution of the lifestyles of chordates^18–20^. The hypothesis of “a new heart for a new head” about the origin of vertebrates^18^, in addition to the evolution of placode and neural crest derivatives as proposed by Gans and Northcutt^21–23^, points to the development of a chambered heart and elaborated branchiomeric muscles as key evolutionary innovations that facilitated the transition from a peaceful filter-feeder lifestyle of ancestral olfactores to a blistering predatory lifestyle of vertebrates. The cardiopharyngeal field is the developmental domain that gives rise to the heart and branchiomeric muscles from a common pool of early cardiopharyngeal progenitors, that after a binary-stepwise process of fate choices gives rise to the first heart field (FHF), the second heart field (SHF) and to branchiomeric muscles (BM), the later including arch muscles involved in mandibular, facial, and branchial functions in the head and neck of vertebrates–reviewed in^18^–. In ascidians, the pharyngeal muscles (i.e. siphon and longitudinal muscles) are considered homologous to vertebrates BM, and their cardiopharyngeal GRN is highly conserved in comparison to vertebrates using a homologous model of binary-stepwise fate choices, thus becoming an attractive system to study heart development at unprecedented spatiotemporal single-cell resolution^20,24–27^. Consistent with this model, the pre-cardiac master regulator *Mesp* is expressed in multipotent progenitor cells both in vertebrates and in ascidians^28–31^. During ascidian gastrulation, *Mesp*+ precardiac cells divide asymmetrically, to give rise to the anterior tail muscles (ATM) under the influence of RA-signaling^32,33^ and to multipotent cardiopharyngeal progenitors (trunk ventral cells, TVCs) under the influence of FGF-signaling^34^. TVCs divide and migrate to the ventral part of the trunk where they will be specified to become first heart progenitors (FHP), second heart progenitors (SHP) and pharyngeal muscle under the influence of FGF and BMP signaling, and the upregulation of cardiac and muscular factors including *Nk4*, *Hand1/2*, *Gata4/5/6*, *Dach* and *Tbx1/10* as some of the crucial members of the cardiopharyngeal GRN also conserved in vertebrates^24,31,34–42^.

In contrast to ascidians, as far as we know, the cardiopharyngeal GRN in appendicularians has never been studied, and therefore whether evolutionary differences between the cardiopharyngeal GRNs of appendicularians and ascidians reflect evolutionary adaptations to the active free-living and adult sessile lifestyles of these two groups of tunicates, respectively, remains unknown. To address this problem, we have performed a genome survey of the cardiopharyngeal GRN in seven appendicularian species, and studied *Oikopleura dioica* as a model to investigate heart development in this group of tunicates. Our work reveals that appendicularians have suffered an evolutionary “deconstruction” of their cardiopharyngeal GRN, including numerous losses of essential genes and cardiopharyngeal subfunctions of GRN components that are crucial in ascidians and vertebrates. The term “deconstruction”, originally coined in Philosophy and later applied in disciplines such as Literature, Architecture, Fashion and Cookery, or even in Developmental Biology^43^ is not synonymous with destruction, but it generally refers to the process of dismantling or breaking apart elements that traditionally are combined, and whose analysis facilitate the recognition of structural modules. In the field of EvoDevo, our work shows how the evolutionary deconstruction by co-elimination of genes and subfunctions during the evolution of appendicularians highlights the modular organization of the cardiopharyngeal GRN. The concept of deconstruction, in contrast to simply erosion, implies that the dismantling does not affect uniformly the GRN, but to particular modules. Our work, for instance, unveils “evolvable modules” that can be related to the evolution of multipotent cellular states that favored the diversification of cardiopharyngeal structures in ascidians and vertebrates, but the loss of pharyngeal muscle and the simplification of the heart in appendicularians. Our results suggest an evolutionary scenario in which the deconstruction of cardiopharyngeal GRN in appendicularians facilitated the transition from an ancestral adult ascidian-like sessile style to a pelagic free-living style based on the evolutionary innovation of the house as the filter-feeding apparatus. Overall, our work provides an example of the “less is more” hypothesis^44^ illustrating how the study of particular gene losses helps to better understand the evolution of adaptations of certain groups of animals^45^, and supports the notion that *O. dioica* is a successful gene loser among chordates that can be used as an attractive evolutionary knockout model to understand the impact of gene loss in the evolution and modular structure of the mechanisms of development in our own phylum (Albalat and Cañestro, 2016; Ferrández-Roldán et al., 2019).

## 2 RESULTS

### 2.1 Developmental atlas of the heart in *O. dioica*

The anatomy of the heart has been thoroughly described in adults of *O. dioica* and some other appendicularian species^47–51^. Due to its chamber-less structure made of two layers (the myocardium and the pericardium) ventrally located between the left stomach and the intestine, it can be probably considered one of the simplest hearts of chordates (**Fig. 1A-A’**). The development of the heart in appendicularian embryos, to the best of our knowledge however, had never been investigated and therefore remained unknown. Thus, we first performed a morphological study by DIC microscopy in live embryos of *O. dioica* to provide the first developmental atlas of the appendicularian heart (**Fig. 1B-E’**). Up to the early hatchling stage (4.5 hpf) (**Fig. 1B**), no morphological evidence could be distinguished to recognize the position of the cardiac precursor cells. By mid-hatchling stage (6 hpf), when the borders delineating organ primordia started to appear throughout the trunk, a striated morphology distinguished from a left-side view allowed us to recognize the heart primordium for the first time located in a ventral position close to the ventral midline, anterior to the notochord, and between the already apparent stomach lobes and intestine (**Fig. 1C**). By 7.5 hpf, ventral view revealed the beginning of the expansion of an internal cavity separating the prospective myocardium and pericardium (**Fig. 1D’**). By 8.5 hpf, the heart started beating suggesting that its contractile properties were already functional (**Sup. video 1**). By late-hatchling stage (9.5 hpf), the internal cavity of the heart was fully expanded, and the position of the heart had asymmetrically relocated towards the left side of the trunk (**Fig. 1E-E’**).

**Fig 1.**
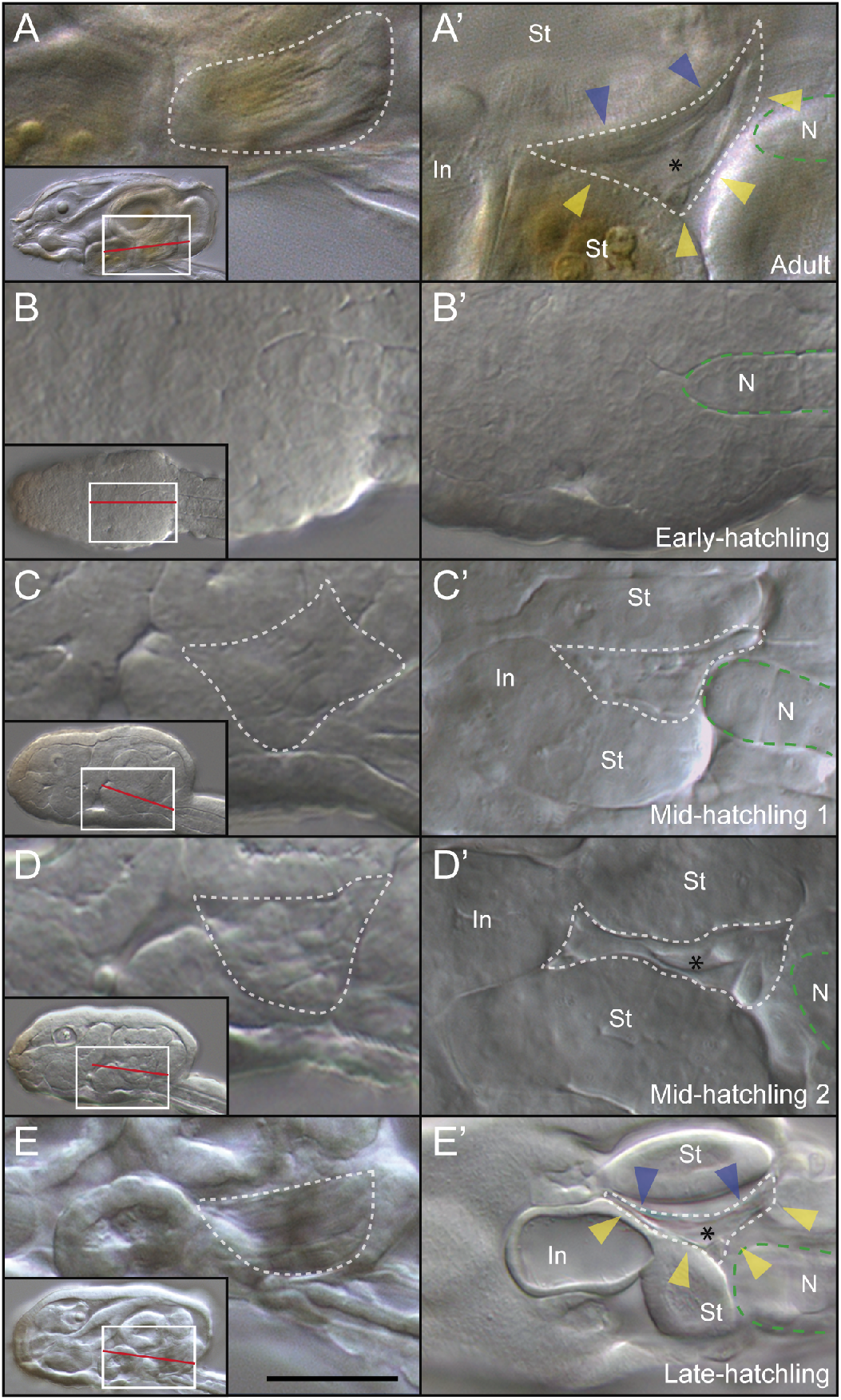
Developmental atlas of heart development in *O. dioica*. (**A**) Anatomy of the heart of an *O. dioica* day 1 adult. The myocardium (blue arrows) beats against the stomach wall. The pericardium (yellow arrows) protects the myocardium. (**B-B’**) In early-hatchling embryos (4.5 hpf) the heart progenitors are still unrecognizable. (**C-C’**) In mid hatchling 1 embryos (6 hpf) heart progenitors are recognizable for the first time. (**D-D’**) In mid-hatchling 2 embryos (7.5 hpf) the internal cavity of the heart is already recognizable. (**E-E’**) In late-hatchling embryos (9.5 hpf) the cavities of the trunk are fully expanded and the heart has already started beating. Capital letters represent left optical sections at the level indicated by white squares. Prime letters represent ventral optical sections at the level indicated by red lines. Discontinuous white lines delimit the shape of the heart during organogenesis. Discontinuous green lines delimit the notochord. Asterisks indicate the internal cavity of the heart. Intestine (In), Notochord (N), Stomach (St). Scale bar represent 20μm in big images and 60μm in small images.

To study the development of the heart at stages at which its precursor cells could not be recognized by their morphology (i.e. early- and pre-hatchling stages), we first intended to use the expression of *Mesp* as the preferred marker used in ascidians to trace the development of the cardiac cell lineage^25^. We surprisingly found, however, that *O. dioica* had lost its *Mesp* homolog (see section 1.2 below), and therefore we used *muscular Actin 1* (*ActnM1*) as an alternative cardiac marker since we had previously found that it was expressed in the heart during *O. dioica* development^52^. *ActnM1* helped us to trace back and describe the cardiac cell lineage in reverse temporal order from hatchlings to 16-cell stage embryos, and therefore to identify the early blastomeres that gave rise to the heart (**Fig 2A-G**). In the ventral part of the trunk, adjacent to the tip of the notochord at early-hatchling stage, we observed one single domain of *ActnM1* expression in which we counted 8 prospective cardiac cells (**Fig. 2A**). While one single *ActnM1* expression domain was still observed at late-tailbud stage (**Fig. 2B**), two distant bilateral domains were distinguished at mid- and early-tailbud stages, in which we counted 3 and 2 prospective cardiac cells in each domain, respectively (**Fig. 2C-D**). At incipient tailbud stage, the bilateral *ActnM1* expression domains were made of one single cell adjacently located to the first anterior tail muscle (ATM) cell (**Fig. 2E**). Integration of the expression data of *ActnM1* with the 4D-nuclear tracing (**Sup. Fig. 1**)^15^ from tailbud up to the 16-cell stage revealed that cardiac cells shared lineage with the first three ATM cells (B8.9, B8.10, and B8.11; **Fig. 2G**), and allowed us to identify B8.12 blastomere as the first cardiac progenitor cell (FCP) in *O. dioica* (**Fig. 2E, G**). We noticed that the onset and temporal progression of the *ActnM1* expression occurred earlier in the left than in the right side, suggesting an asynchronous bilateral asymmetry on the development of the FCP cells. At 64-cell stage, we observed *ActnM1* expression in cells of the vegetal hemisphere compatible with B7.5 and B7.6, which are the precursor cells of the ATM and the FCP (**Fig. 2F-G**). At 32-cell stage, according to the 4D cell fate map reconstruction (**Sup. Fig. 1**) we inferred that B6.3 should be considered the first precardiac precursor that will give rise to both the heart and ATM lineages (**Fig. 2G**), but no *ActnM1* signal was distinguished over background levels at this stage. B6.3 is the sister cell of the germline precursor B6.4, both of which descended from B5.2 located in the vegetal hemisphere at the 16-cell stage (**Fig. 2G**). Thus, our findings allowed us to infer that the cardiac precursors of appendicularians and ascidians shared cell lineage fate maps and origin from the same early blastomeres. Moreover, our observations in *O. dioica* suggested that the FCP was also originated from an asymmetric division in the tail-trunk interface that also gives rise to the anterior tail muscles in the same way as it occurs in ascidians, in both cases ending in a final ventral distant position of the trunk from the anterior muscle cells of the tail. These findings, therefore, provide solid ontogenetic evidence that the heart of ascidians and appendicularians were homologous.

**Fig 2.**
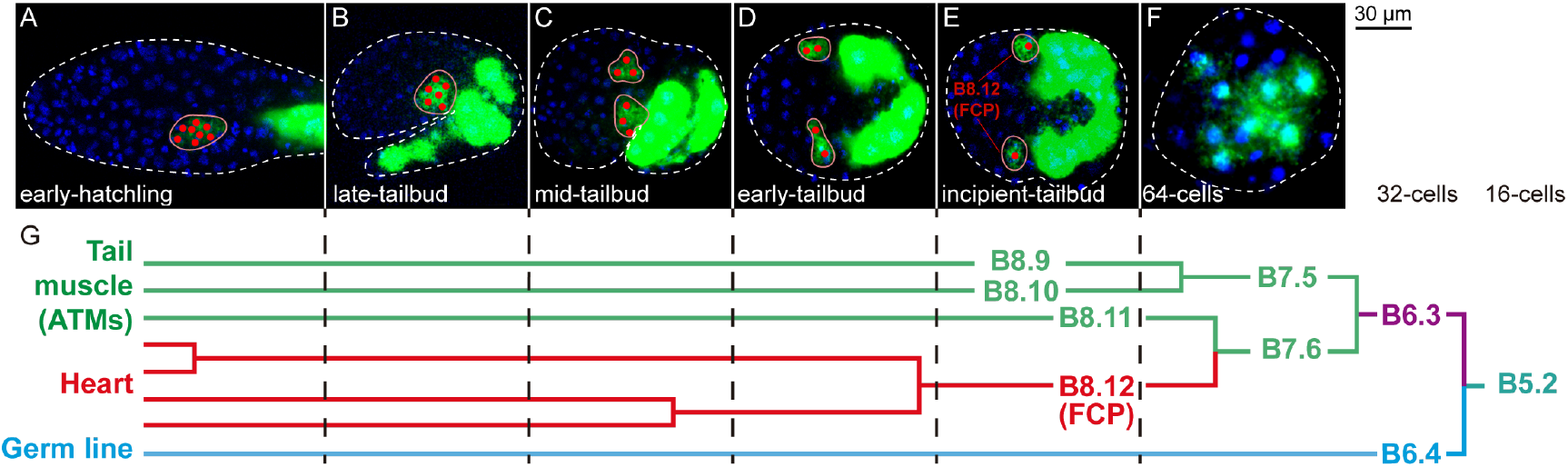
Integration of *ActnM1* expression with the fate map of the tail muscle, the heart, and the germline in *O. dioica*. (**A-F**) Dorsal sections of different embryo stages stained with *ActnM1* to detect the cardiac progenitors throughout development. Nuclei are stained with Hoechst (blue). (**G**) Cell lineage diagram showing a B5.2 blastomere from one side that originates three anterior muscle cells of the tail, the heart, and the germline (Modified from ^15^). To facilitate comparisons, the nomenclature of the blastomeres is according to that of Conklin for ascidians (1905). Fates are color-coded. Close shapes encircle cardiac progenitors. White dashed lines enclose the embryo. Red dots indicate the nuclei of the cardiac precursors. FCP, first cardiac progenitor; ATM, anterior tail muscle cell.

### 2.2 Loss of an early multipotent state module during the deconstruction of the cardiopharyngeal GRN

To investigate the evolution of the cardiogenetic toolkit of *O. dioica*, we performed a comprehensive in silico survey of conserved components of the GRN responsible for the development of the cardiac progenitor cells in ascidians and vertebrates^18,19^. In order to map gene losses in the context of tunicate evolution, we included in the survey the genomes of six other appendicularian species^53^ as well as eleven ascidian species^54^.

First, we started by searching the *O. dioica* homolog of *Mesp*, which is the earliest known marker of pre-cardiac progenitors in ascidians and vertebrates^31,55,56^. Using ascidian *Mesp* and vertebrate *Mesp1* and *Mesp2* proteins as tBLASTn queries, however, none of the resulting hits returned *Mesp* in best reciprocal blast hits (BRBH), but members of other bHLH gene families such as *Math*, *Achaete scute*, *Neurogenin*, and *Hand* (**Sup. Table 3**). Analyses by tBLASTn of the other six appendicularian species also revealed the absence of *Mesp* homologs in all their genomes, which contrasted with its presence in all eleven ascidian species (**Fig. 3**). The apparent absence of *Mesp* among BLAST hits was confirmed by phylogenetic analyses, suggesting that appendicularians had lost the homolog of *Mesp* (**Fig. 3** and **Sup. Fig. 2A**). To test for the possibility of ‘function shuffling’ among bHLH genes upon the loss of *Mesp*, we checked for the expression of the gene that gave the most significant blast hits (i.e. *Math*), but we did not observe any expression domain compatible with the position of the cardiac progenitors (**Sup. Fig. 2B-G**). These findings, therefore, suggested that the loss of *Mesp* occurred at the base of the appendicularian clade after its split from the ascidian lineage, and therefore, the activation of the cardiac pathway in appendicularians became *Mesp*-independent.

**Fig. 3.**
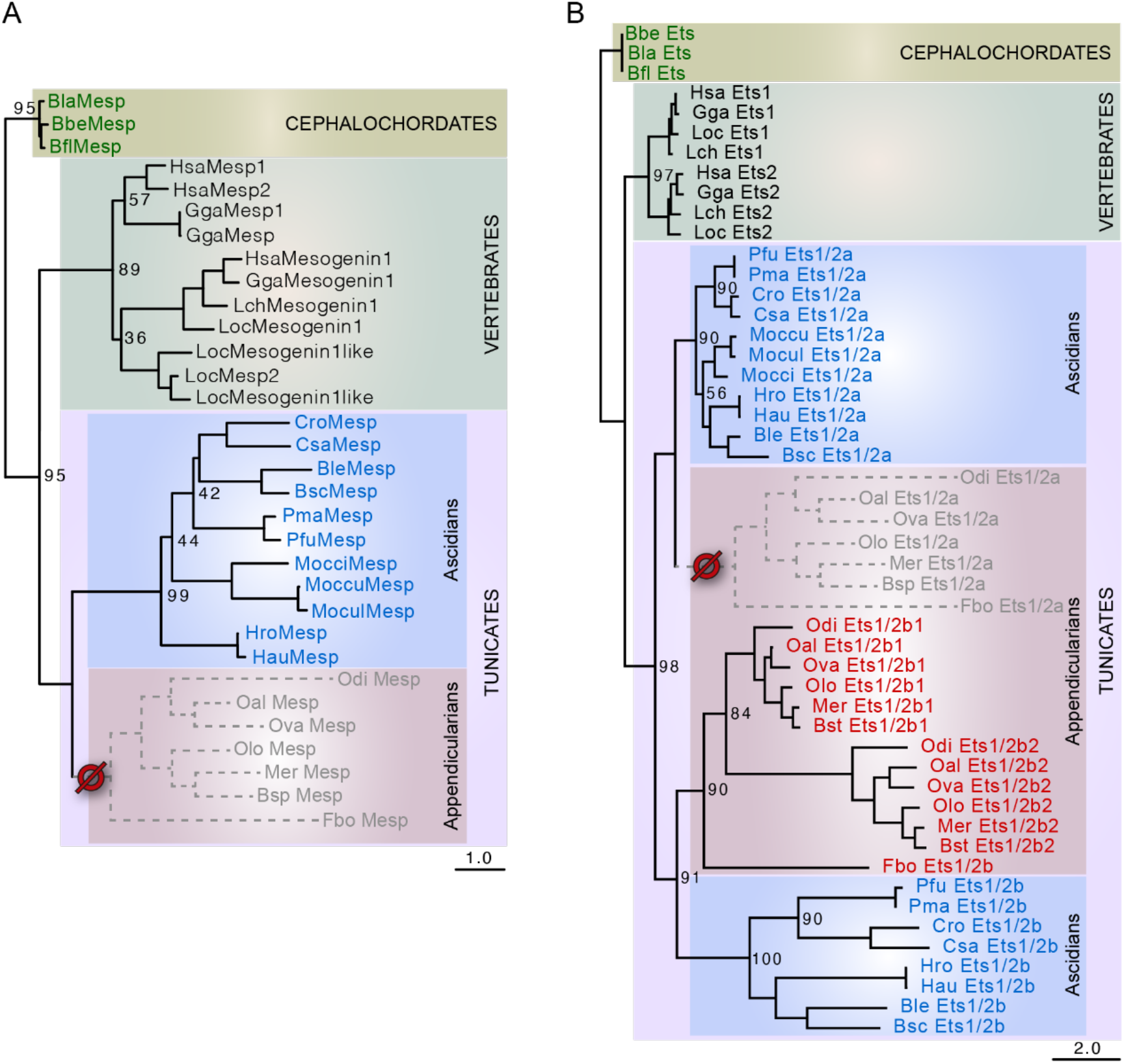
ML phylogenetic trees of *Mesp* (A) and *Ets1/2* (B) families in chordates revealing the loss of *Mesp* and *Ets1/2a* in appendicularians. The best blast hits of *Mesp* in *O. dioica* belong to close related gene families, such as *Math* and *Neurogenin* (Sup. Fig. 2A). Phylogenetic analyses of Ets1/2 including related gene families such as ERG, revealed the presence of two Ets1/2 genes in appendicularians (Sup. Fig. 3A). The phylogenetic tree of chordate Ets1/2 genes suggested that the ancestor of tunicates duplicated *Ets1/2* (*Ets1/2a* and *Ets1/2b*), and the ancestor of appendicularians lost *Ets1/2a* but duplicated *Ets1/2b* (**B**). Scale bar indicates amino acid substitutions. Bootstrap values are shown in the nodes. Vertebrates: *Gallus gallus* (Gga), *Homo sapiens* (Hsa), *Latimeria chalumnae* (Lch), *Lepisosteus oculatus* (Loc); Tunicates: *Bathochordaeus sp*. (Bsp). *Botrylloides leachii* (Ble), *Botrylloides schlosseri* (Bsc), *Ciona robusta* (Cro), *Ciona savignyi* (Csa), *Fritillaria borealis* (Fbo), *Halocynthia aurantium* (Hau), *Halocynthia roretzi* (Hro), *Mesochordaeus erythrocephalus* (Mer), *Molgula occidentalis* (Mocci), *Molgula occulta* (Moccu), *Molgula oculata* (Mocul), *Oikopleura albicans* (Oal), *Oikopleura dioica* (Odi), *Oikopleura longicauda* (Olo), *Oikopleura vanhoeffeni* (Ova), *Phallusia fumigata* (Pfu), *Phallusia mammillata* (Pma); Cephalochordates: *Branchiostoma belcheri* (Bbe), *Branchiostoma floridae* (Bfl), *Branchiostoma lanceolatum* (Bla).

We continued our survey with the search of homologs of *Ets1/2*, the direct downstream target of *Mesp* in the TVCs of ascidians, whose phosphorylation by the FGF/MAPK pathway results in the upregulation of the primary cardiogenic transcription factors^34^. BLAST searches revealed the presence of two potential *Ets1/2* homologs in *O. dioica* as well as in all other appendicularians, which was corroborated by phylogenetic analyses including closely related gene families such as *Erg* (**Sup. Fig. 3A**). Interestingly, our gene survey also revealed the presence of a previously unnoticed second *Ets1/2* gene in all ascidian species (**Sup. Fig. 3A**). Phylogenetic tree of the chordate *Ets1/2* gene family, rooted with amphioxus as an outgroup, suggested that *Ets1/2* was duplicated at the base of the tunicates, giving rise to two paralogs that we have named as *Ets1/2a* and *Ets1/2b*, being the former the direct target of *Mesp* involved in the specification of the cardiopharyngeal lineage in ascidians. This phylogenetic tree, however, suggested that the two *Ets1/2* genes of appendicularians were co-orthologs to the ascidian *Ets1/2b* and that they had been generated by a gene duplication occurred within the appendicularian lineage (*Ets1/2b1* and *Ets1/2b2*), while no *Ets1/2a* homolog was present in appendicularians (**Fig. 3B**). Experiments by whole-mount in situ hybridization with *Ets1/2b1* and *Ets1/2b2* in *O. dioica* revealed no expression domains compatible with cardiac precursors, but expression in the notochord, the tail muscle cells, the migratory endodermal strand, the ventral organ, and the oikoplastic epithelium (**Sup. Fig. 3B-M**).

Our findings, therefore, suggested that *Ets1/2a* was lost in appendicularians, and none of the *Ets1/2b* paralogs played any similar role to *Ets1/2a* in the cardiogenic pathway in ascidians. Moreover, we also analyzed the components of the FGF/MAPK pathway, since in ascidians this is responsible for *Ets1/2a* phosphorylation in the TVCs^34^. Our results revealed the absence of many of the key components of the FGF/MAPK pathway, including the absence of the homolog of the *MEK1/2*, which in ascidians, through a kinase cascade mediated by ERK, is responsible for Ets1/2a phosphorylation in the cardiogenetic pathway (**Sup. Fig. 4A)**. Blast analysis revealed a single clear homolog of *ERK* in *O. dioica* (e-value 2e^-105^; **Sup. Table 3**), whose analysis by whole-mount in situ hybridization revealed no expression in the cardiac region, but in the oikoplastic epithelium (**Sup. Fig. 4B-H)**.

Next, we searched for *O. dioica* homologs of *Gata4/5/6*, *FoxF*, *Nk4*, and *Hand1/2* families since in ascidians phosphorylated Ets1/2a activates *FoxF* and *Gata4/5/6*, two members of the primary cardiac transcription factors (CTF) in charge of cardiopharyngeal migration and specification, respectively^37^, and other downstream members of the cardiopharyngeal kernel as *Nk4*, *Hand1/2*, and *Hand-r*^25,26,57^. Blast searches and phylogenetic analyses revealed the absence of *Gata4/5/6* homologs in *O. dioica*, but the presence of four paralogs of the closely related *Gata1/2/3* (**Sup. Fig. 5)**. The absence of *Gata4/5/6* homologs in all six appendicularians, but their presence in all ascidians, suggested that the loss of *Gata4/5/6* occurred at the base of the appendicularian lineage, which was accompanied by an expansion of the *Gata1/2/3* subfamily (**Sup. Fig. 5**). In the case of *FoxF* and *Nk4*, we did find a single copy of each one in *O. dioica* and all appendicularian species (**Sup. Fig. 6 and 7**). Finally, blast searches revealed the presence of a single *Hand1/2* homolog in *O. dioica*, as well as in all other six appendicularian species, and the presence of a second paralog *Hand-r* in all ascidian species. The origin of ascidian *Hand-r* is unclear^58^, and our phylogenetic analyses did not allowed us discard if *Hand-r* was originated by an ascidian-specific duplication, and therefore its absence from appendicularians was not due to a gene loss, or on the contrary it was already present in the last common ancestor of tunicates, and subsequently lost in appendicularians (**Sup. Fig. 8**).

In an initial expression analysis by colorimetric WMISH of members of the *Gata1/2/3*, *FoxF*, *Nk4*, and *Hand1/2* gene families, we observed expression domains that were in the nearby of the presumptive cardiac region. Therefore, we performed double fluorescent in situ hybridization with *ActnM1* to test if any of these genes were expressed in the cardiac progenitors (**Fig. 4**). At the incipient tailbud stage, we observed that *Nk4* was the first CTF to be expressed in the FCP, as soon as B8.12 was originated from the division of B7.6 but, we did not observe any signal in the daughter cell B8.11 that remained positioned posteriorly as the first ATM (**Fig. 4A**). Expression of *Nk4* was also visible earlier on the left than on the right side, consistently with the asynchronous bilateral asymmetry on the development of the FCP also revealed by *ActnM1* expression (**Fig. 4A, A’**). By mid/late-tailbud stage, we observed that *Nk4* signal faded and was more difficult to detect (**Fig. 4A’’**). At that stage, we observed *Hand1/2* signal for the first time in some embryos in the cardiac progenitors localized in the ventral midline of the trunk (**Fig. 4B’’**). However, we did not observe *FoxF* signal in the cardiac precursors (**Fig 4 C-C’’’**), but in adjacent epidermal cells that continued positive for *FoxF* to finally become restricted to the Fol domain of the oikoplastic epithelium (**Fig. 4C’’’)**. Despite the absence of *Gata4/5/6* homolog in *O. dioica*, we tested for the possibility of function shuffling among paralogs by checking the expression of two of the four *Gata1/2/3* paralogs that seemed to have expression in the nearby of the cardiac area. Double fluorescent experiments, however, revealed that the *Gata1/2/3b* and *Gata1/2/3d* were not co-expressed with *ActnM1* in the FCP, but in an adjacent epidermal cell (**Fig. 4D-D’ and E-E’**). At late-tailbud and hatchling stages, the expression of *Gata1/2/3b* and *Gata1/2/3d* had expanded in lateral epidermal domains and anterior prospective placodal populations, respectively, but were never detected in the heart primordium (**Fig. 4D’’-D’’’ and E’’-E’’’**). These results, therefore, suggest that *FoxF* and *Gata* factors did not participate in the specification or migration of the cardiac progenitors in *O. dioica*, in contrast to their roles in ascidians and vertebrates. Overall, our findings of the gene co-elimination of *Mesp*, *Ets1/2a*, *MEK1/2*, *Gata4/5/6* and RA signaling^59^, and the loss of the cardiac subfunctions of *FoxF*, highlight a process of evolutionary deconstruction of the cardiopharyngeal gene regulatory network in *O. dioica* with the loss of what can be considered an “early multipotent” (EM) module that in ascidians is related to the early maintenance of the multipotent state of the TVCs after their split from the ATM lineage. In *O. dioica*, the cells resulting from the split of the ATM lineage rapidly activated the onset of the cardiac kernel (i.e. *Nk4* and *Hand1/2*), the reason why we have termed them “first cardiac precursors” (FCP), rather than TVCs as in ascidians.

**Fig. 4.**
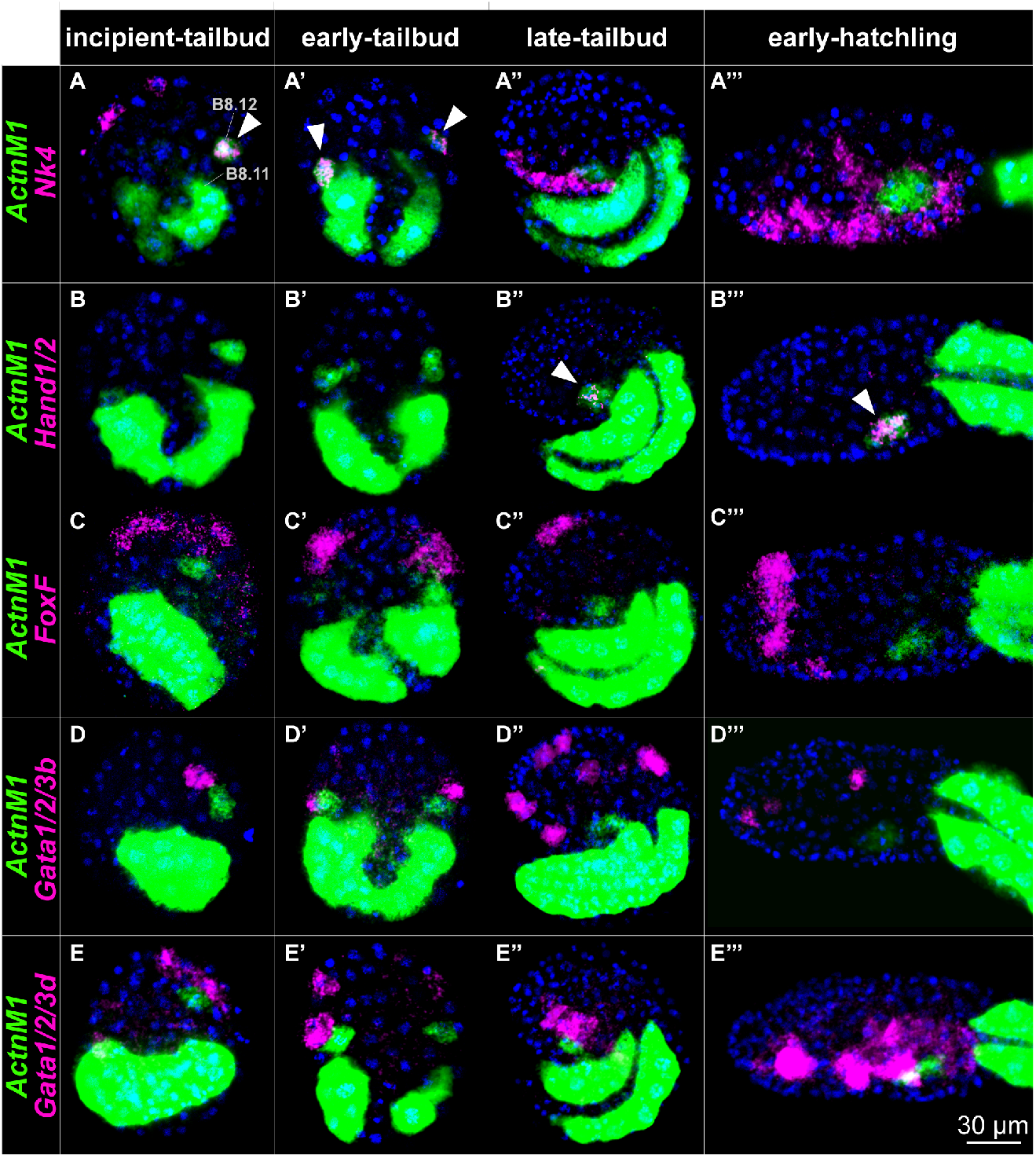
Developmental expression patterns of *O. dioica ActnM1* and prospective cardiac transcription factors. Double fluorescent in situ hybridization of *ActnM1* with *Nk4*, *Hand1/2*, *FoxF*, *Gata1/2/3b*, and *Gata1/2/3d*. *Nk4* expression signal was detected in ventral epidermis and the FCP (B8.12) from the inipient-tailbud stage (**A**) until the early-tailbud (**A’**). In later stages, we only detected expression in the epidermis, but not in the cardiac precursors (**A’’-A’’’**). *Hand1/2* was specifically expressed in the cardiac progenitors from late-tailbud to hatchling stages (**B-B’’’**). We did not detect expression of *FoxF*, *Gata1/2/3b*, nor *Gata1/2/3d* in cardiac precursors, but they were expressed in different epidermal domains (**C-E’’’**). Incipient- and early-tailbud stages correspond to ventral views oriented anterior towards the top. Late-tailbud and early-hatchling stages correspond to lateral views oriented anterior towards the left and dorsal towards the top. White arrows indicate co-expression of *ActnM1* with the corresponding gene in cardiac progenitors.

### 2.3 Loss of a late multipotent state module during the deconstruction of the cardiopharyngeal GRN

In ascidians, the TVCs undertake a series of asymmetric cell divisions and regulatory transient states through a process of binary fate decisions that do not only give rise to first and second heart precursors, but also to the atrial siphon muscle founder cells (ASMF)^20^. In our work, we then investigated *O. dioica* homologs of *Hand-r*, *Tbx1/10*, *Islet1* and *Ebf* (*Coe*), which in ascidians become activated in a FGF/MAPK-dependent manner to determine the trajectory towards the ASMF including the activation of *MyoD* (*Mrf*)^20,26,41^. Our genome survey and phylogenetic analysis revealed that in addition to the absence of *Hand-r* (see above), none of the *Tbx* genes of *O. dioica* belonged to the *Tbx1/10* family (**Sup. Fig. 9A**). The absence of *Tbx1/10* in all examined appendicularian species suggested that this gene was lost at the base of the appendicularian clade after its split from the ascidian lineage (**Sup. Fig. 9A**). In the case of *Islet1* and *Ebf*, despite the existence of one homolog of each in *O. dioica*, our results by WMISH revealed that the two of them were expressed in the nervous system, but no expression was found that could suggest the presence of presumptive muscle cells in the gill slits homologous to the atrial muscle cells of ascidians (**Sup. Fig. 9B-L**). Finally, we have identified the *MyoD* homolog in *O. dioica* which did not appear to be expressed in muscle cells, but its expression domains were restricted to some fields of the oikoplastic epithelium during late hatchling stages (**Sup. Fig. 9M-Q**). Our results, therefore, revealed that the loss of pharyngeal muscle during the evolution of appendicularians was accompanied by the loss of *Tbx1/10* and the loss of the muscle subfunctions of *Islet1*, *Ebf* and *MyoD*.

To test for the presence of presumptive second heart field in *O. dioica*, we investigated the expression of the homolog of *Dach*, which in ascidians is activated by *Tbx1/10* in the absence of FGF/MAPK-signaling, and it is sufficient to determine the second heart precursors identity^26^. Our genome survey revealed the presence of a single homolog of *Dach* in *O. dioica* (**Sup. Table 3**). Experiments of WMISH revealed, however, that *Dach* was expressed in the nervous system, the endostyle, and the trunk epidermis, but not in the heart of *O. dioica* (**Sup. Fig. 10**). The absence of *Dach* expression in the heart could suggest that *O. dioica* might lack a homolog to the second heart field of ascidians. In addition to *Dach*, single-cell transcriptomic analysis in ascidians has revealed a total of 18 cell-specific gene markers for the first heart precursors and 7 for the second^26^. Our gene survey by BRBH revealed that 12 out of those 25 cell-specific gene markers were absent in the genome of *O. dioica* (**Sup. Table 3**). Using the human genome as an outgroup, the BRBH results were compatible with 4 gene losses in *O. dioica* (3 and 1 for FHP and SHP specific gene markers, respectively), and 8 ascidian specific gene duplications (6 and 2 for FHP and SHP markers, respectively). Altogether, our results highlights that during the deconstruction of the cardiopharyngeal gene regulatory network the loss of *Tbx1/10* and the loss of subfunctions of *Dach, Islet1, Ebf* and *MyoD* might represent the loss of a late module of multipotent states that in ascidians is responsible of the differentiation of the SHF and atrial muscle, structures that appeared to have been lost during the evolution of appendicularians.

Finally, to investigate how the daughter cells of the cardiac progenitors differentiated into myocardium and pericardium in hatchling stages, we analyzed the expression patterns of some of the CTFs we had found in *O. dioica* such as *Hand1/2*, *Rapostlin (FNBP1)*, and *Nk4* as well as some structural genes that code for motor proteins such as *ActnM1*, *Myosin*, *FilaminC* (*FlnC*), and *Troponins T* (*TnnT*) (**Fig. 5A-I’’**). WMISH experiments revealed that the expression of *Hand1/2*, that started after the downregulation of *Nk4*, was specifically maintained in the heart throughout all hatchling stages (**Fig. 5A-A’’**). Despite the first heart beatings did not occur until late-hatchling (8,5 hpf), in the early-hatchling stage (5 hpf), in addition to *ActnM1*, we also observed the first expression signals of some motor genes such as *FilaminC* and *TnnT7* (**Fig. 5F, I**). By mid-hatchling stage (6,5 hpf), in addition to *Hand1/2*, we observed expression signals of the CTF *Raspostlin*, as well as additional motor genes such as *Myosin*, *TnnT1*, and *TnnT4* (**Fig. 5B’, E’, G’, H’**). Finally, by late-hatchling stage (8 hpf), we observed a second wave of *Nk4* expression, which had been downregulated since mid-tailbud stage (**Fig. 4A’’**). The ventral view of late hatchlings, in which the expansion of internal cavities allowed us to clearly differentiate the myocardium and pericardium, revealed that some of the analyzed genes were expressed both in the pericardium and myocardium (*Hand1/2*, *Rapostlin*), while others only in the myocardium (*Nk4*, *mAct1*, *TnnT1*, *TnnT4*, *Tnnt7*, and *FilaminC*). These results suggest that despite the loss of the early and late modules, many downstream regulators and structural genes that characterize the cardiac program in ascidians and vertebrates were conserved in *O. dioica*, and our results provide the first insights into the spatio-temporal dynamics of the expression patterns of CTF and motor-genes that will be useful for further functional investigations of heart development in appendicularians.

**Fig. 5.**
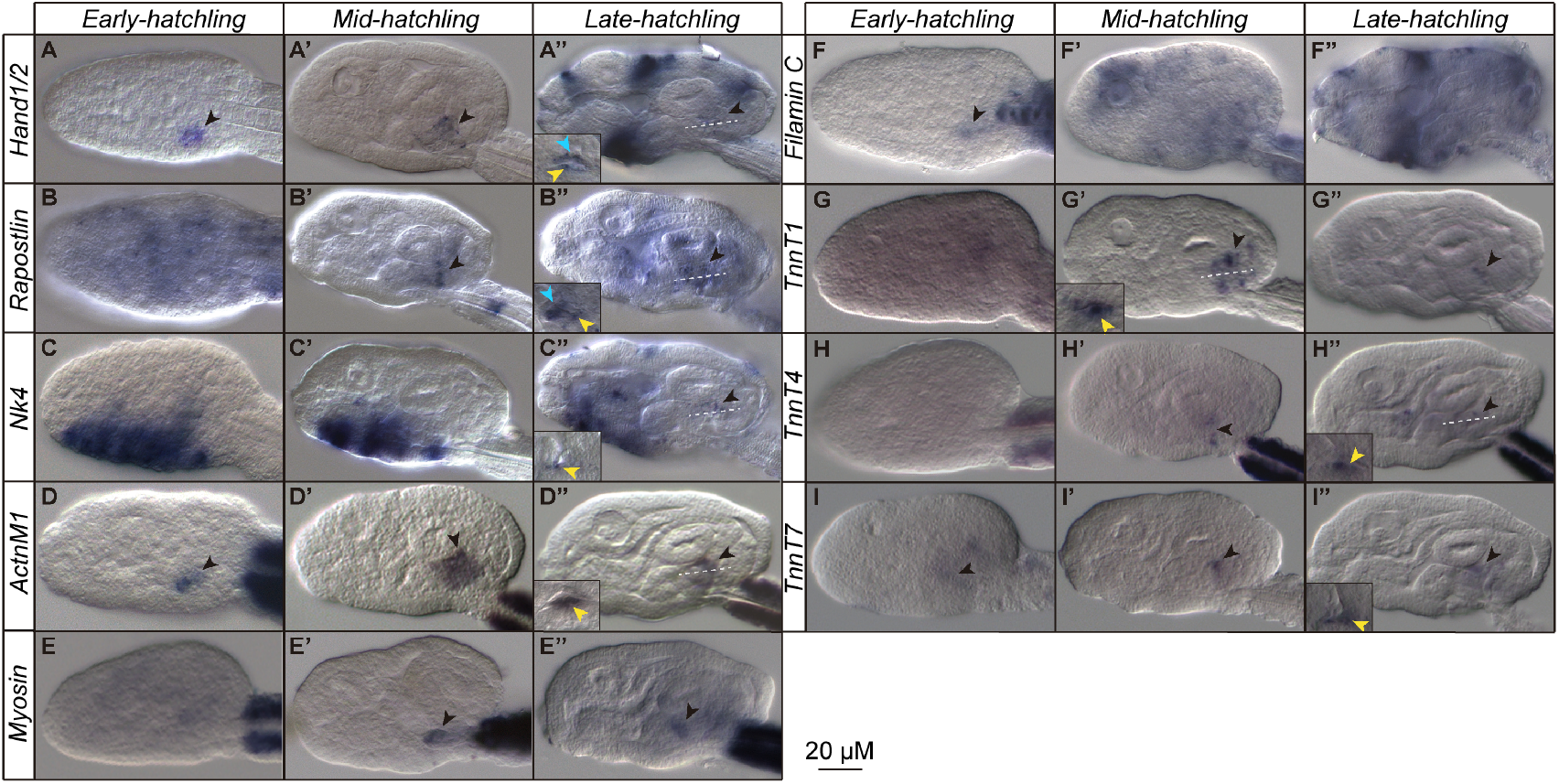
Expression atlas of cardiac genes during hatchling stages of *Oikopleura dioica* development. Developmental expression analysis by whole mount in situ hybridization of *O. dioica* cardiac transcription factors (**AC’’**) and motor proteins (**D-I’’**). Images correspond to lateral views oriented anterior to towards the left and dorsal towards the top. Detailed small images represent ventral optical section at the level indicated by white lines. Black arrows indicate expression in the heart. Blue arrows indicate expression in the pericardium. Yellow arrows indicate expression in the myocardium.

### 2.4 FGF and BMP signaling are not involved in the specification of cardiac progenitor cells in *O. dioica*

In ascidians, the FGF/MAPK signaling via phosphorylation of Ets1/2a is essential for the onset of the pan-cardiac program in the cardiopharyngeal precursors^34^. This induction event takes place only in the anterior pairs of B7.5 daughters turning them into TVCs, while the posterior B7.5 descendants become into ATMs under the influence of RA^34,60^. The loss of *Ets1/2a* in appendicularians, the absence of *Ets1/2b* expression domains compatible with a cardiac function, and the loss of *MEK1/2* in the FGF/MAPK pathway suggested the hypothesis that the determination of the FCP could have become FGF independent in *O. dioica*. To test this hypothesis, we performed FGF inhibitory treatments with SU5402, an inhibitor of FGF receptor (FGFR). Treatments with SU5402 induced developmental alterations, whose severity depended on the concentration and duration of the treatment (**Sup. Table 4**). In treatments starting at 2-cell stage, SU5402 concentrations at 50 μM induced obvious malformations at gastrula stage, affecting the proper formation of tailbud morphologies and arresting development before hatching (**Sup. Table 4**). In treatments starting after gastrulation at 32-cell stage, however, SU5402 concentrations of 50 μM did not produce any obvious altered phenotype, and required concentrations of at least 100 μM to induce aberrant morphologies in hatchling stages (**Sup. Table 4**). WMISH experiments in embryos treated with SU5402 at 50 μM from the 2-cell stage revealed that a majority of the embryos showed aberrant or no *ActnM1* signal, suggesting that the development of mesodermal derivatives such as the muscle lineage was severely affected (**Fig. 6A-B’**). Interestingly, those few embryos that were able to achieve incipient tailbud morphologies with two recognizable rows of tail muscle cells labeled by *ActnM1*, also showed the presence of presumptive cardiac progenitors, comparably to DMSO treated control embryos **(Fig. 6B’’)**. In embryos treated with SU5402 at 100 μM from the 32-cell stage, in which gastrulation was not arrested, WMISH with *Nk4* revealed the presence of cardiac precursors in most embryos, similarly to control DMSO embryos at incipient tailbud stage **(Fig. 6D-D’)**. These results suggested that despite FGF appeared to be important for early development during gastrula stage, FGF was not essential for the specification of the cardiac progenitors at incipient tailbud stage. These results together with the loss of *Mesp*, *Ets1/2a* and *MEK1/2* support the hypothesis that the specification and induction of cardiac progenitors have become FGF independent in appendicularians, in contrast to ascidians.

**Fig. 6.**
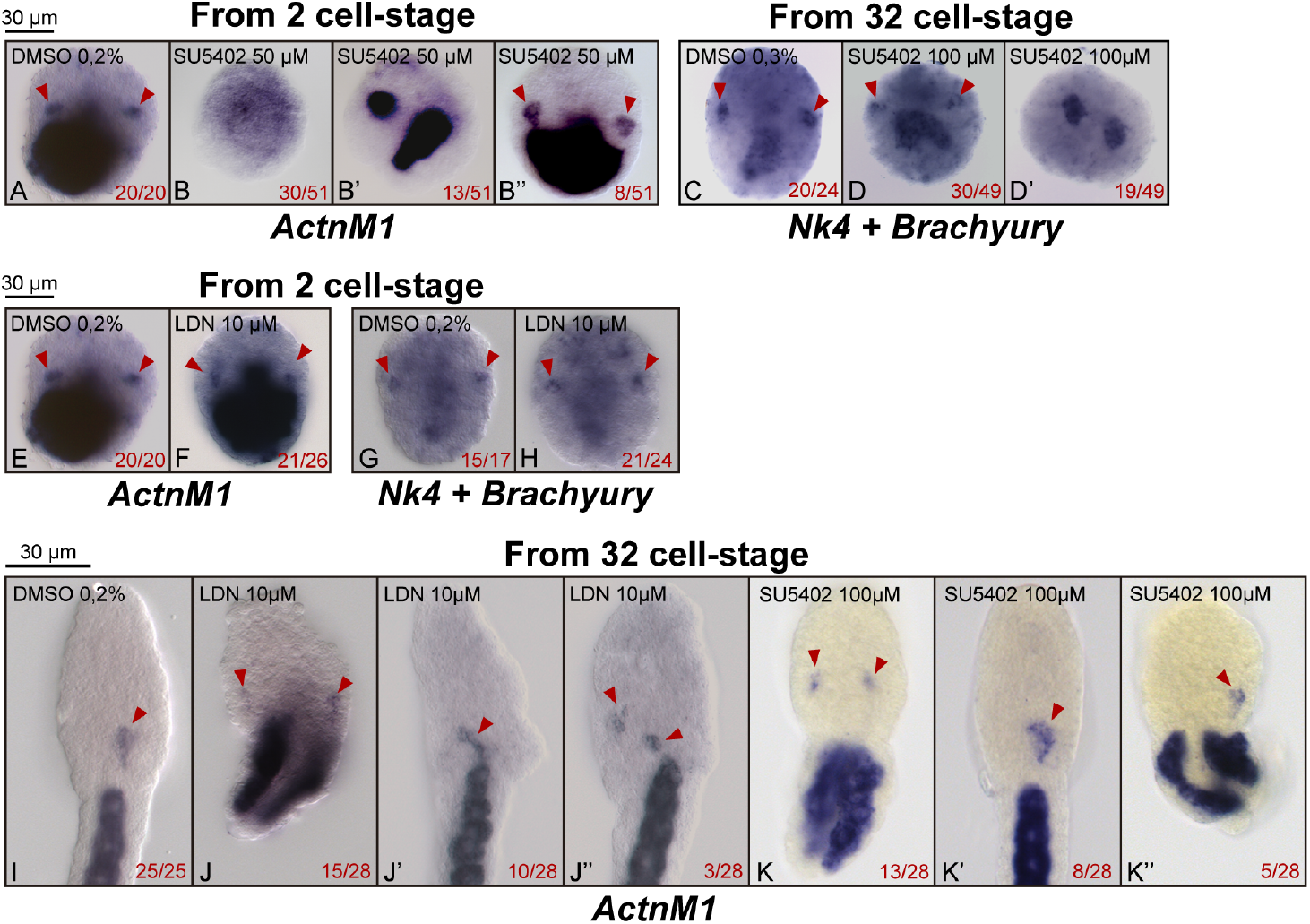
Effects FGF and BMP inhibitions in the development of the heart of *O. dioica*. Whole mount in situ hybridization of *ActnM1* in DMSO-control (**A**) and treated embryos with SU5402 50μM starting at 2-cell stage (**B-B’’**), in which more than a half of the embryos showed no specific signal of *ActnM1* (**B**). Whole mount in situ hybridization of *Nk4* and *Brachyury* in DMSO-control (**C**) and SU5402 100μM starting at 32-cell stage (**D-D’**), in which more than a half of the embryos showed expression of *Nk4* in the cardiac progenitors but alterations in the notochord development (**D**). Embryos that did not show *Nk4* signal corresponded with those that had been arrested (**D’**). Whole mount in situ hybridization of *ActnM1* in DMSO-control (**E**) and LDN 10μM starting at 2-cell stage in which most of the embryos displayed an *ActnM1* expression comparable to the DMSO control (**F**). Whole mount in situ hybridization of *Nk4* and *Brachyury* in DMSO-control (**G**) and LDN 10μM from 2-cell stage in which most of the embryos displayed *Nk4* expression in the cardiac progenitors (**H**). Whole mount in situ hybridization of *ActnM1* in DMSO-control (**I**) and treated embryos with LDN 10μM (**J-J”**) and SU5402 100μM (**K-K’’**) starting at 32-cell stage, in which many embryos display separated cardiac progenitors (**J, J’’, K, K’’**), rather than one single domain in the midline such in DMSO-control (**I**). Tailbud embryos were stained using the cross-hybridizing ActnM1 probe^56^. Hatchling embryos were stained with the specific ActnM1 probe^56^. Tailbud embryos images correspond to dorsal views with anterior to the top. Hatchling images represent ventral views with anterior to the top. Red arrowheads indicate cardiac precursors.

In ascidians, in addition to FGF, BMP signaling is also relevant for cardiac development contributing to the specification and migration of the TVCs. The migration, activated by *FoxF*, follows a gradient of BMP originated in the ventral epidermis^24^. The increasing levels of BMP initiate the upregulation of cardiac kernel genes as *Gata4/5/6* or *Nk4*, which sustains and arrests migration, respectively^24^. Our results showing the loss of *Gata4/5/6* and the apparent absence of expression of the migratory factor *FoxF* in the FCP of *O. dioica* suggested the hypothesis that the role of BMP in the cardiac gene regulatory network could have changed in appendicularians. To test this hypothesis, we performed BMP inhibitory treatments with LDN–a highly specific inhibitor of BMP receptors– during different time windows (**Sup. Table 4)**. WMISH experiments using *ActnM1* and *Nk4* probes in tailbud embryos that had been previously treated with 10 μM LDN from the 2-cell stage revealed no differences in the formation of the FCPs nor in the onset of the cardiac kernel marker *Nk4* in comparison to DMSO-control embryos (**Fig. 6F, H**). These results suggested that BMP was not necessary for the specification of the FCP nor the activation of the expression of the cardiac kernel as soon as the FCP split from the ATMs.

We also checked for the convergence of cardiac progenitors in the ventral part of the trunk after treating embryos with FGF and BMP inhibitors (**Fig. 6I-K**). In early hatchling DMSO control embryos, *ActnM1* revealed that cardiac progenitors were localized in a single domain near the trunk midline, while in SU5402 and LDN treated embryos cardiac precursors were bilaterally distributed far from the midline (**Fig. 6J, J’’, K**). Interestingly, both, BMP and FGF inhibitory treatments, also affected the elongation of the tail and its rotation relative to the trunk. Thus, while a ventral view of control embryos showed a row of muscle cells (**Fig. 6I**), most of the treated embryos showed two rows revealing that the rotation had not taken place (**Fig. 6J, J’’, K**). These results suggested that the inhibition of BMP- and FGF-signaling could affect the positioning of heart precursors towards the midline, as well as other developmental processes such as the elongation and rotation of the tail.

## 3 DISCUSSION

### 3.1 The hearts of appendicularians and ascidians are homologous

The vast morphological variability among hearts and pumping organs across metazoans has made cardiac development a hot topic in the discussion of homologies and analogies in the field of EvoDevo^35,61^. In the case of chordates, numerous studies have provided strong evidence supporting that the hearts of vertebrates and ascidians are homologous despite the great morphological differences between the multichambered complex heart of vertebrates and the tubular simple heart of ascidians^25^. At the beginning of our study, however, the striking morphological and physiological differences between the heart of ascidians and *O. dioica* casted doubt on whether these two organs were truly homologous, or on the contrary, could be analogous pumping organs. At the morphological level, the cylindrical V-shape heart of ascidians contrasts with the chamberless flat heart of *O. dioica*. At the physiological level, while in ascidians the heart does not start beating until several days after the metamorphosis in juveniles, in *O. dioica* the heart starts beating against the stomach wall as soon as 8.5 hpf at late-hatchling stage. Despite these marked differences, our results provide strong evidence that both hearts are homologous by revealing that the cardiac precursor cells of *O. dioica* and *C. robusta* share the same cell fate map and the same ontogeny. In both organisms, the muscle/heart lineage descends from the B5.2 blastomere located in the posterior-vegetal hemisphere at 16-cell stage, which also gives rise to the germline lineage (**Fig. 7**). In *O. dioica*, the division of B5.2 at the 32-cell stage splits the germline and muscle/heart lineages, being B6.3 the precursor of the later. In ascidians, however, the split of the germline and muscle/heart lineages occurs one cleavage later at the 64-cell stage, being B7.5 therefore, the muscle/heart precursor equivalent to B6.3 in *O. dioica* (**Fig. 7**).

**Fig. 7.**
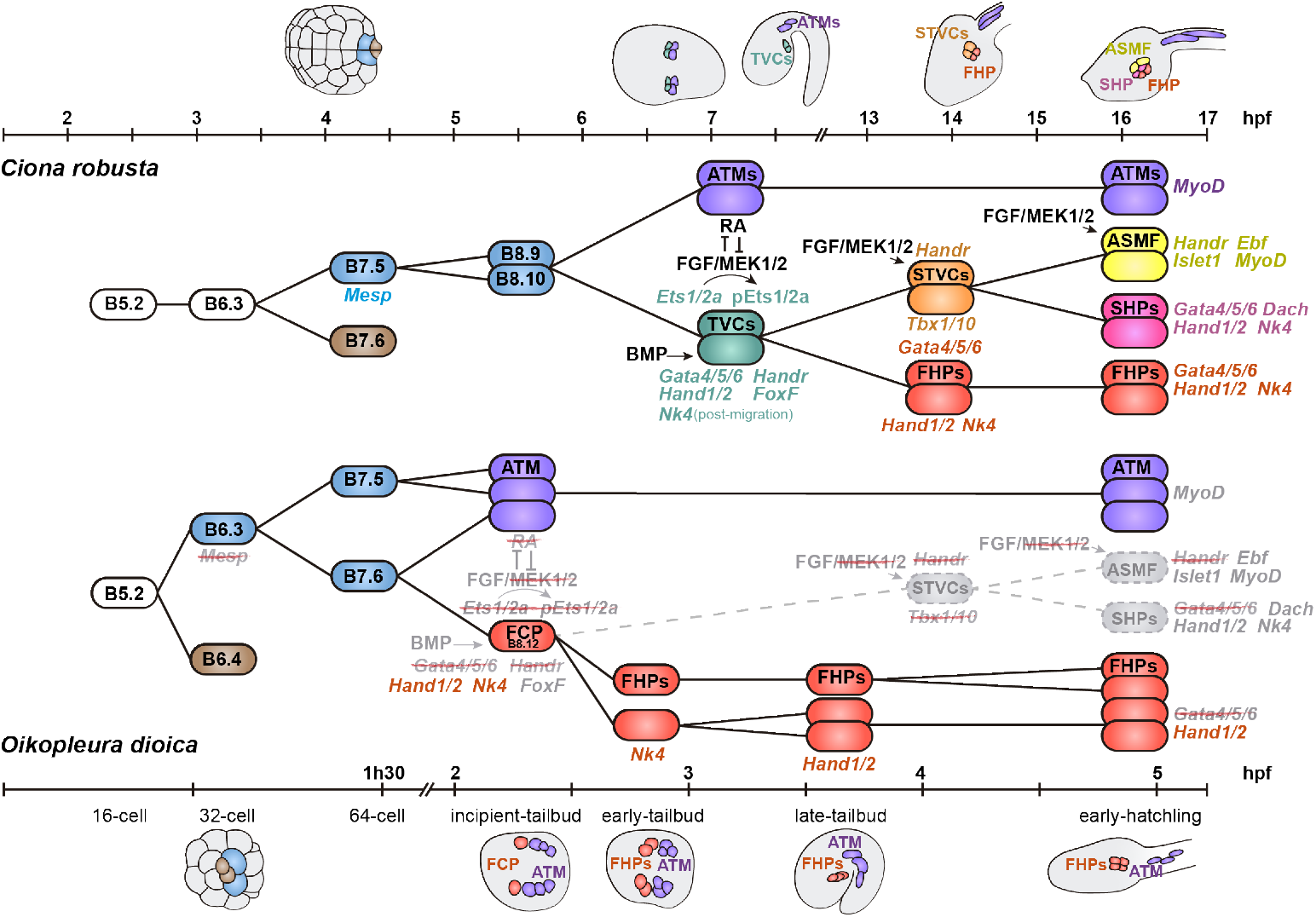
Cell lineage diagrams of *C. robusta* and *O. dioica* cardiopharyngeal progenitors. The hearts of *Oikopleura* and *Ciona* share the same cell fate map and the same ontogeny with minor differences. While ascidians comprise two TVCs and two ATMs per side, *Oikopleura* only comprises one FCP and one ATM per side. Moreover, in *Oikopleura* the germline (brown), the cardiac precursors (green), the ATM (purple), and the FHPs (red) appear one cleavage earlier than in ascidians. Genes and structures that do not play a role in cardiac development in *O. dioica* are represented in grey, absent genes are strikethrough.

Moreover, the origin of the first cardiac precursors in *O. dioica* and *C. robusta* (FCP and TVCs, respectively) share the same ontogenic origin, since in both cases likely occur through an asymmetric division at the interface between the trunk and tail, in which the anterior daughter cell will end up in the ventral part of the trunk giving rise to the heart, while the posterior daughter cell will remain in the tail giving rise to the ATMs. The common ontogenic origin of the cardiac lineage and the most anterior axial muscles can be therefore considered an ancestral feature of tunicates, homologous to the shared origin of the cardiac and branchiomeric muscles in vertebrates. The timing of the split between the cardiac and the axial muscle lineages between *O. dioica* and *C. robusta* occurs again one cleavage earlier in the appendicularian (after the division of B7.6 at the approximately 110-cell incipient-tailbud stage) than in the ascidian (after the divisions of B8.9 and B8.10 at the approximately 300-cell post-neurula stage). This timing difference is probably connected to the difference in the number of cardiac precursors between *O. dioica* and ascidians, with just one single FCP in each side in *O. dioica* in contrast to the two TVCs in each side in ascidians, as well as just one single row of muscle cells in each side of the tail of appendicularians in contrast to the two rows in the tail of ascidians^52,62^.

The “one-cleavage earlier” trend of fate decisions observed in this work is consistent with the observations already made by Delsman back in 1910^63^ when describing that cell ingression during gastrulation in *O. dioica* occurred one cleavage earlier than in ascidians. This idea that fate restriction occurs earlier in *O. dioica* than in ascidians has been corroborated by the characterization of the cell lineage fate map in *O. dioica*^15,64^, as well as by expression analyses showing for instance that the onset of *Otx*^65^ and *Brachyury*^66^ during early embryogenesis occurred “one-cleavage earlier” than in ascidians. This trend of “one-cleavage earlier” fate decision in *O. dioica* could be considered as a generalized feature of the evolution of the developmental program of appendicularians, which might have likely contributed to the acceleration of development, morphological simplification and cell number reduction of this group of tunicates.

### 3.2 Modular deconstruction of the cardiopharyngeal gene regulatory network

One of the most striking findings of our work is the numerous losses of genes and cardiopharyngeal subfunctions that have suffered the cardiopharyngeal GRN in appendicularians. In agreement to the modular model for the control of heart cell identity proposed by Wang et al., (2019)^26^, our comparative analysis between ascidians and appendicularians allows proposing an evolutionary scenario in which we can recognize how the modular deconstruction of the cardiopharyngeal GRN might have affected the evolution of the heart and siphon muscle in appendicularians (**Fig. 7**). In ascidians, the binary fate decision between the ATM and TVC lineages occurs in the trunk-tail interface, under the antagonistic posterior influence of retinoic acid signaling promoting the fate of ATMs in the tail, and the anterior influence of FGF signaling promoting the multipotent cardiopharyngeal fate of the TVCs in the trunk^32–34^. FGF action is mediated by the phosphorylation of Ets1/2a, which had been previously upregulated by *Mesp* in the TVCs^32–34^. The multipotent state of the TVCs is maintained until the *FoxF*-dependent ventral migration of the TVCs, and the action of *Gata4/5/6* and BMP-signaling from the ventral epidermis that triggers the cardiogenic kernel upon the activation of *Nk4*^26,31,36,37^. In *O. dioica*, however, our results show that the binary fate decision between the ATM and FCP lineages occurs in a very different molecular context due to the gene losses of *Mesp* and *Ets1/2a*, the loss of *MEK1/2* affecting the FGF/MAPK signaling pathway, and the loss of the retinoic acid signaling pathway (**Fig. 7**)^59^. Our results also show that, in contrast to TVCs, FCPs activate the expression of *Nk4* as soon as B8.12 blastomeres are born from the division of B7.6, suggesting that the FCP is not multipotent, but determined to the cardiac fate, as revealed by the activation of the cardiogenic kernel. This is the reason why we have termed these cells as FCPs instead of TVCs as in ascidians. Our results, moreover, show that the activation of *Nk4* is independent of *Gata4/5/6* (which has been lost in appendicularians), independent of *FoxF* (which is not expressed in the FCP), and independent of FGF- and BMP-signaling (as shown by our inhibitory treatments). In the light of these results, we propose an evolutionary scenario in which all these losses highlights the deconstruction of an “early-multipotent” (EM) module of the cardiopharyngeal GRN that in ascidians contributes to the maintenance of the multipotent state in the TVCs, which in the FCP of *O. dioica* has been lost.

In ascidians, the multipotent states of TVCs and second trunk ventral cells (STVCs) allow the differentiation of the second heart field and the atrial siphon and longitudinal muscles. In appendicularians, however, our results revealing the ancestral loss of *Tbx1/10*, the loss of FGF and BMP cardiac inductive functions, together with the losses of expression of *Dach* in the heart and *Islet1*, *Ebf* and *MyoD* in muscles of *O. dioica* highlights the deconstruction of a “late multipotent” (LM) module of the cardiopharyngeal GRN that in ascidians is responsible for the maintenance of the multipotency in the STVC and fate specification of the SHP and ASMF (**Fig. 7**). These results suggest, therefore, an evolutionary scenario in which the heart of *O. dioica* might be homologous to the FHP of ascidians, while the homologous tissues to the SHP, atrial siphon muscles, and longitudinal muscles have been lost.

### 3.3 Evolution of pelagic appendicularians from a sessile ascidian-like ancestor

The deconstruction of GRN modules that are key for ascidian cardiopharyngeal development suggests a parsimonious evolutionary scenario in which appendicularians might have suffered a process of evolutionary simplification from an ancestral ascidian-like condition. These findings are compatible with the adaptive evolution of three innovations that likely facilitated the transition from an ancestral sessile ascidian-like lifestyle to the pelagic free lifestyle of appendicularians upon the innovation of the house: *I*) an accelerated cardiac development, *II*) the formation of an open-wide laminar heart and *III*) the loss of siphon muscle.

The “one-cleavage earlier” trend and the loss of the EM module likely facilitated the earlier activation of the cardiac developmental kernel in appendicularians than in ascidians, causing an accelerated cardiac development that resulted in the activation of the beating of the heart by 8.5 hpf in *O. dioica*, in contrast to few days after metamorphosis in ascidians. Accelerated cardiac development, therefore, could have been plausibly selected as an adaptation to the high energetic demand of appendicularians associated with their intense and constant beating of the tail to power water circulation through the house right after the inflation of the first house as soon as 10 hpf, which clearly differs from the sporadic movements of the tail at larval stages used for dispersion both in appendicularians and ascidians^46,67^.

In addition to accelerated cardiac development, the low number of cells in the cardiac lineage together with the apparent loss of the SHP in appendicularians is compatible with the transformation of an ascidian-like tubular heart into an open-wide laminar structure that beats against the stomach like in *O. dioica*. Considering that hemolymph circulation in appendicularians is not only powered by the heart, but also by the movements of the tail^47^, the adaptive innovation of a laminar cardiac structure plausibly offered a more efficient system to pump hemolymph waves propelled by the tail movements through an open-wide structure than through the less accessible space of a tubular ascidian-like heart.

Finally, the loss of muscles to propel fluid inside the organism could be considered the result of regressive evolution during the transition from a sessile filter-feeding ascidian-like strategy, in which contraction of siphon muscles is responsible for water current, to a pelagic filter-feeding strategy based on the innovation of the appendicularian house, in which water circulation is mostly propelled by tail beating^68^. Considering that in addition of tail beating, water circulation is also propelled by fined tuned movement of the cilia in the gills that can efficiently reverse the direction of the water flow, inwards during feeding or outwards during the inflation of the house, it is also plausible to think that instead of being a process of regressive evolution, the loss of siphon muscle could have been adaptive by reducing the interference of graceless muscle contractions with the orchestrated movement of the cilia that precisely regulate the direction of the water flow^69,70^.

In conclusion, our results support an evolutionary adaptive scenario in which the deconstruction of cardiopharyngeal GRN could have contributed to the loss of features that characterize ascidian-like sessile lifestyle, and the acquisition of novel features that enabled its transition to a pelagic free-living active style connected to the innovation of the house **(Fig. 8)**. The conclusion that the last common ancestor of tunicates was sessile is compatible with the commonly accepted assumption that appendicularian branching is basal among tunicates, but it is also compatible with the possibility that appendicularians are phylogenetically related to ascidians of the Aplousobranchia order^14^. Our conclusion that the last common tunicate ancestor was sessile provides a novel framework for future comparative studies of the cardiopharyngeal GRN responsible for the development of diverse cardiac structures among different ascidian and appendicularian species and for future efforts to clarify the phylogenetic relationship between appendicularians and the rest of tunicates.

**Fig 8.**
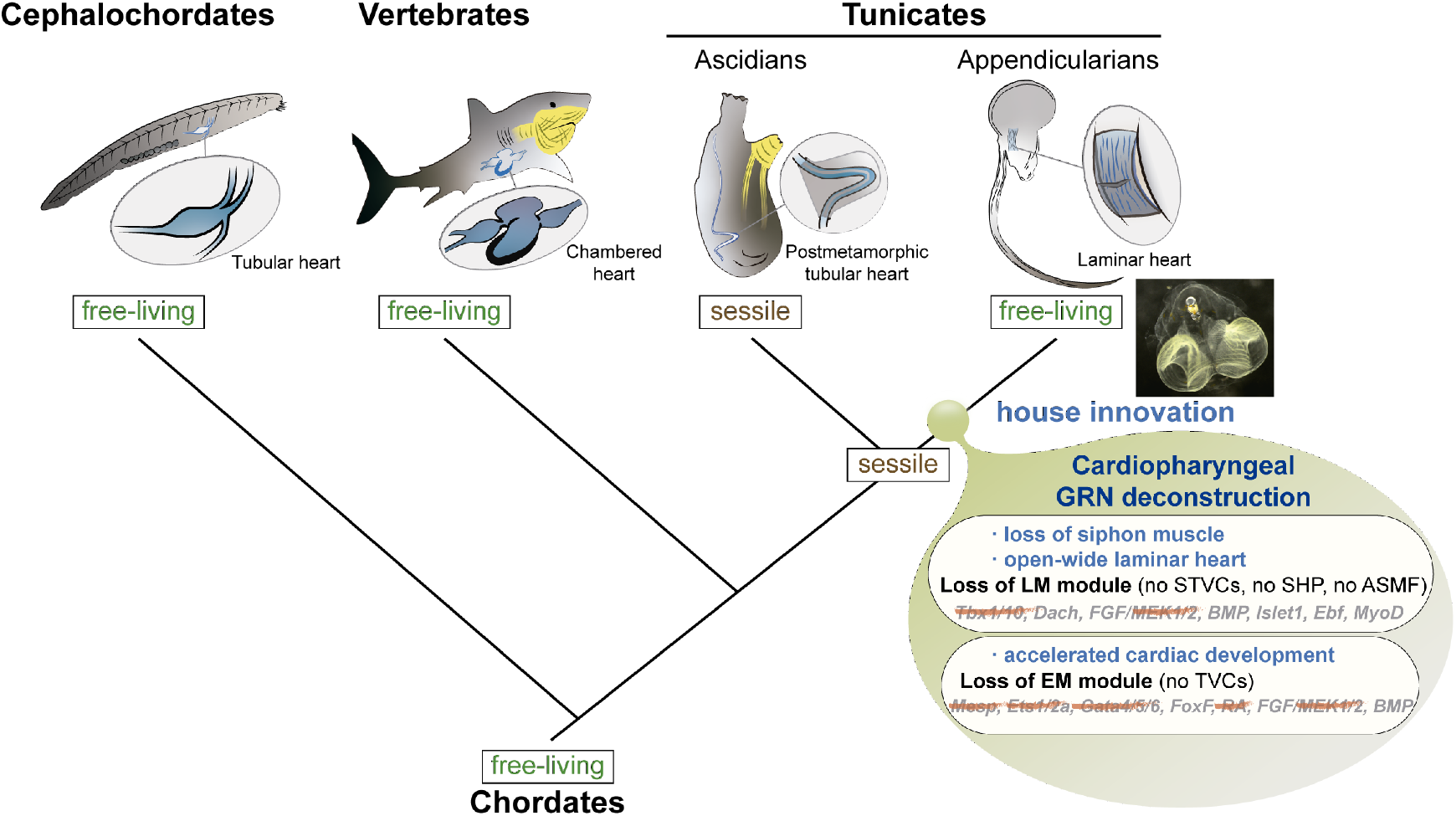
The deconstruction of the cardiopharyngeal GRN in appendicularians favored their transition from an ascidian-like ancestral condition to a free-living house-based life style. The loss of the siphon muscle, the development of an open-wide laminar heart, and the acquisition of an accelerated development can be considered adaptive consequences of the deconstruction of the cardiopharyngeal GRN that facilitated the innovation of the free-living life style of appendicularians based on the house filtrating device. GRN, gene regulatory network; STVCs, second trunk ventral cells; SHP, second heart progenitors; ASMF, atrial siphon muscle field; TVCs, trunk ventral cells; LM, late multipotent; EM, early multipotent.

## 4 Materials and methods

### 4.1 Biological material

*O. dioica* specimens were obtained from the Mediterranean coast of Barcelona (Catalonia, Spain). Culturing of *O. dioica* and embryo collections have been performed as previously described in^71^.

### 4.2 Genome databases searches and phylogenetic analysis

Protein sequences from the tunicate *C. robusta* and the vertebrate *Homo sapiens* were used as queries in BLASTp and tBLASTn searches in genome databases of selected species: https://blast.ncbi.nlm.nih.gov/Blast.cgi for *Branchiostoma floridae*, *Branchiostoma belcheri*, *Branchiostoma lanceolatum*, *Gallus gallus*, *Lepisosteus oculatus*, and *Latimeria chalumnae*; http://www.aniseed.cnrs.fr/ for ascidian species (*Ciona savignyi*, *Phallusia fumigata*, *Phallusia mammillata*, *Halocynthia roretzi*, *Halocynthia aurantium*, *Botryllus schlosseri*, *Botryllus leachii*, *Molgula occulta*, *Molgula oculata*, and *Molgula occidentalis*); http://oikoarrays.biology.uiowa.edu/Oiko/ for *O. dioica*, and a local blast for six other appendicularian species (*Oikopleura albicans*, *Oikopleura vanhoeffeni*, *Oikopleura longicauda*, *Mesochordaeus erythrocephalus*, *Bathochordaeus stygius*, *Fritillaria borealis*) with public genomes^53^. The orthology between cardiac genes was initially assessed by blast reciprocal best hit (BRBH)^72^ and subsequently by phylogenetic analysis based on ML inferences calculated with PhyML v3.0 and an automatic substitution model^73^ using protein alignment generated by the MUSCLE^74^ program and reviewed manually. Accession numbers are provided in **Sup. Table 1**.

### 4.3 Cloning and expression analysis

*O. dioica* genes were PCR amplified from cDNA obtained as described in^59^. Then, they were cloned using the Topo TA Cloning^®^ Kit (K4530-20, Invitrogen) to synthesize antisense digoxigenin (DIG) and fluorescein (FITC) riboprobes for whole-mount in situ hybridization (WMISH) and double fluorescent whole-mount in situ hybridization (FWMISH) (**Sup. Table 2**). The WMISH were performed as previously described^59,65,66^.

For FWMISH, fixed embryos were rehydrated in PBT (PBS/0.2%Tween-20), treated with 50 mM DTT in PBT (for 10 min at room temperature (RT)), washed in 0.1 M triethanolamine in PBT (2 x 5 min at RT), treated with two successive dilutions of acetic anhydride (0.25% and 0.5%) in 0.1 M triethanolamine (10 min at RT), and washed in PBT (2x for 5 min at room temperature). Prehybridization was carried out in a mixture of 50% formamide, 5x SSC, 0.1 mg/ml heparin, 0.15% Tween 20, 5 mM EDTA, 0.5 mg/ml yeast RNA and 1x Denhardt’s reagent for 2 hours at 63°C. Then, hybridization was carried out in the same solution but adding the two probes at 0.5-1 ng/μL each, overnight at 63°C. Next day, embryos were washed in successive dilution of SSC (2x for 10 min in 2xSSC/0.2%Tween-20 at 65°C; 2x for 10 min in 0.2xSSC/0.2%Tween-20 at 65°C; 1x for 5 min in 0.1xSSC/0.2%Tween-20 at RT) and 2x for 5 min in MABT (0.1 M maleic acid, 0.15 M NaCl, 0.1% Tween 20, pH 7.5). Blocking was performed by washing the embryos in a mixture of MABT, 2.5 mg/ml BSA and 5% sheep serum. Finally, anti-fluorescein antibody conjugated with POD (1:1000 in blocking solution) (11426346910, Roche) was added to the samples for overnight incubation at 4°C. The day after, samples were washed in MABT (8x for 15 min at RT) and then in TNT (0.1 M Tris-HCl pH7, 0.15M NaCl, 0.3% TritonX-100) for 10 minutes. For the staining, embryos were incubated in TSA-tetramethylrhodamine (NEL742001KT, Perkin Elmer) for 10 min. Then, they were washed in TNT for 10 min, in PBT for 10 min, in 2%H2O2/PBT for 45 min, in PBT (2x for 5 min), and in MABT (2x for 5 min). Then, a second blocking step was performed followed by the addition of an anti-Digoxigenin-POD (11207733910, Roche) antibody that was incubated overnight at 4°C. The morning after, as the previous day, the samples were washed and the coloration reaction was added, that in this case included TSA-Fluorescein system-green (NEL741E001KT, Perkin Elmer) for 1 hour 30 min. After coloration, embryos were washed in TNT (2 x 5 min at RT) and PBT (2 x 5 min at RT). Mounting was made in 80%glycerol/PBS with Hoechst-33342 1μM (Invitrogen-62249). A confocal microscopy LSM880 (Zeiss) was used for imaging of samples and FIJI^75^ was used to compose the confocal series and adjust the brightness and contrast.

### 4.4 Pharmacological treatments

For FGF inhibition, animals were treated with 50 μM and 100 μM of SU5402 (SML0443, Merk) from 2-cell stage (30 minutes post fertilization, mpf) and from 32-cell stage (70 mpf), respectively, to hatchling stage (4 hours post fertilization, hpf) in darkness. For BMP inhibition, animals were treated with 10μM of LDN (SML1119, Merk) from 2-cell stage and from 32-cell stage until hatchling stage. To perform these treatments, eggs were pooled in 4mL of SSW and fertilized with 200 μL of sperm dilution (the sperm of 3 males in 5 mL of SSW). At the desired time, embryos were transferred to a 3 mm Petri dish plate with 4 mL of treatment solution at 19°C. Control embryos were incubated in DMSO 0,2% or 0,3% (v/v) depending on the treatment. The effects of the treatments were scored by in situ hybridization. For tailbud embryos, we used cross-hybridizing *ActnM1*, *Nk4* and *Brachyury* probes^52,76^, while for hatchling embryos we used the specific *ActnM1* probe^52^ (**Sup. Table 2**).

## 6 Acknowledgements

The authors thank to A. Moncusí, M. Plana-Carmona and P. Bujosa for technical assistance with *Myosin*, *TnnT* and *Fgf* initial analyses, all team members of the C.C. and R.A. laboratories for fruitful discussions, and to S. Artime for assistance with the animal facility. C.C. was supported by BFU2016-80601-P and PID2019-110562GB-I00, R.A. by BIO2015-67358-C2-1-P and J.G-F. by BFU2017-861152-P grants from Ministerio de Ciencia y Innovatión (Spain). C.C. R.A. were also supported by grant 2017-SGR-1665 from Generalitat de Catalunya. A.F-R. was supported by a FPU14/02654, G.S-S. by FPU18/02414 fellowship from Ministerio de Educatión Cultura y Deporte, M.F-T. by PREDOC2020/58 fellowship from the University of Barcelona.

## 7 Author contributions

A.F-R carried cardiac developmental atlas, genome surveys, phylogenetic analyses, WMISH experiments, cell lineage mapping and BMP inhibitory treatments. M. F-T. contributed to *Gata* genome survey and FWMISH. G. S-S. contributed to FGF genome survey and FGF inhibitory treatments. E. D-B. contributed to *Tbx* genome survey and WMISH. M.J-L. contributed to *Ets* and *Tbx* phylogenies. A.F-R interpreted the data and made the figures. C.C. conceptualized the project, J.G-F. and R.A. provided resources, R.A. and C.C. supervised the experiments. C.C. and A.F-R. wrote the MS. All authors commented on the manuscript and agreed to its final version.

## 9 Ethics approval

This project does not involve any ethical issues related to informed consent, data protection issues, or Humans. The experimentation of aquatic invertebrate animals such as the planktonic *Oikopleura dioica* is not subjected to the regulation of animal experimentation, because this only applies to vertebrates organisms (Real Decreto 223 de 14 Marzo de 1998, en Cataluña Ley 5/1995,DOGC2073,5172). In any case, experimental procedures will follow the EU animal care guidelines, and have been approved by the Ethical Animal Experimentation Committee (CEEA-2009) of the University of Barcelona.

