## Supplementary material for "Cardiopharyngeal deconstruction and ancestral tunicate sessility"

**Supplementary Table 1**. Accession numbers of the sequences used in phylogenetic analysis.

#### Mesp/Math/Neurogenin

| Gene name | Gene/Scaffold ID | | Database |
|---|---|---|---|
| BbeMath6 | XP_019619932.1 | | NCBI |
| BbeMesp | XP_019645972.1 | | NCBI |
| BbeNeurogenin | XP_019635448.1 | | NCBI |
| BflMath6 | XP_002596084.1 | | NCBI |
| BflMesp | ABD57444.1 | | NCBI |
| BflNeurogenin | AAF81766.1 | | NCBI |
| BlaMath6 | BL15626_cuf0 | manual annotation | NCBI: Bl71nemr |
| BlaMesp | BL14804_evm1 | manual annotation | NCBI: Bl71nemr |
| BlaNeurogenin | ACE79717.1 | | NCBI |
| BleMath | Boleac.CG.SB_v3.S636.g12639.01.p | | Aniseed |
| BleMesp | S12 | manual annotation | Aniseed |
| BleNeurogenin | Boleac.CG.SB_v3.S623.g12503.01.p | | Aniseed |
| BleNeurogenin | Boleac.CG.SB_v3.S887.g15003.01.p | | Aniseed |
| BscMath | Boschl.CG.Botznik2013.chr5.g36966.01.t | | Aniseed |
| BscMesp | ChrUn | manual annotation | Aniseed |
| BscNeurogenin | Boschl.CG.Botznik2013.chr10.g56010.01.p | | Aniseed |
| BspMath | SCLE01239960 | manual annotation | NCBI: ASM436795v1 |
| BspNeurogenin | SCLE01432724 | manual annotation | NCBI: ASM436795v1 |
| CroMath6 | KH.C9.872_XP_002122506.1 | | Aniseed |
| CroMesp | KH.C3.100_NP_001071761.1 | | Aniseed |
| CroNeurogenin | KH.C6.129 | | Aniseed |
| CsaMath8 | Cisavi.CG.ENS81.R132.671158-674092.07637.p | | Aniseed |
| CsaMesp | BAD18072.1 | manual annotation | Aniseed |
| CsaNeurogenin | Cisavi.CG.ENS81.R532.88248-90192.05863.p | | Aniseed |
| FboMath | SDII01115774 | manual annotation | NCBI: ASM436807v1 |
| FboNeurogenin | SDII01071635 | manual annotation | NCBI: ASM436807v1 |
| FboNeurogenin | SDII01061863 | manual annotation | NCBI: ASM436807v1 |
| GgaMath8 | NP_001171210.1 | | NCBI |
| GgaMesogenin | NP_990015.2 | | NCBI |
| GgaMesp | CAD12882.1 | | NCBI |
| GgaMesp1 | XP_003641880.1 | | NCBI |
| GgaNeurogenin | NP_990214.1 | | NCBI |
| GgaNeurogenin | XP_003641519.2 | | NCBI |
| GgaNeurogenin | NP_990127.1 | | NCBI |
| HauMath | S239 | manual annotation | Aniseed |
| HauMesp | S7142 | manual annotation | Aniseed |
| HauNeurogenin | Haaura.CG.MTP2014.S859.g06918.01.p | | Aniseed |
| HroMath6 | CG.MTP2014.S5.g02224.01.p | | Aniseed |
| HroMesp | Harore.CG.MTP2014.S97.g06901.01.p | | Aniseed |
| HroNeurogenin | CG.MTP2014.S5.g08306.01.p | | Aniseed |
| HsaMath8 | NP_116216.2 | | NCBI |
| HsaMesogenin | NP_001099039.1 | | NCBI |
| HsaMesp1 | NP_061140.1 | | NCBI |
| HsaMesp2 | NP_001035047.1 | | NCBI |
| HsaNeurogenin | NP_076924.1 | | NCBI |
| HsaNeurogenin | NP_066279.2 | | NCBI |
| HsaNeurogenin | NP_006152.2 | | NCBI |
| LchMath8 | XP_006003191.1 | | NCBI |
| LchMesogenin | XP_005997585.1 | | NCBI |
| LchNeurogenin | XP_006003536.1 | | NCBI |
| LchNeurogenin | XP_006004992.1 | | NCBI |
| LchNeurogenin | XP_006011419.1 | | NCBI |
| LocMath | XP_006629700.2 | | NCBI |
| LocMesogenin | XP_015213357.1 | | NCBI |
| LocMesogenin | XP_006629065.1 | | NCBI |
| LocMesogenin | XP_015198957.1 | | NCBI |
| LocMesp2 | XP_006629064.1 | | NCBI |
| LocNeurogenin | XP_015202930.1 | | NCBI |
| LocNeurogenin | XP_015205245.1 | | NCBI |
| MerMath | SCLF01217498 | manual annotation | NCBI: ASM436797v1 |
| MerNeurogenin | SCLF01145285 | manual annotation | NCBI: ASM436797v1 |
| MocciMath | Moocci.CG.ELv1_2.S264788.g06238.01.p | | Aniseed |
| MocciMesp | S145165 | manual annotation | Aniseed |
| MocciNeurogenin | Moocci.CG.ELv1_2.S647877.g29858.01.p | | Aniseed |
| MoccuMath | Mooccu.CG.ELv1_2.S602351.g36667.01.p | | Aniseed |
| MoccuNeurogenin | S438802 | manual annotation | Aniseed |
| MoculMath | Moocul.CG.ELv1_2.S30383.g01453.01.p | | Aniseed |
| MoculMesp | Moocul.CG.ELv1_2.S92333.g07428.01.p | | Aniseed |
| MoculNeurogenin | Moocul.CG.ELv1_2.S15977.g00531.01.p | | Aniseed |
| MooccuMesp | S460719 | manual annotation | Aniseed |
| OalMath | SCLG01000009 | manual annotation | NCBI: ASM436787v1 |
| OalNeurogenin | SCLG01000119.1 | manual annotation | NCBI: ASM436787v1 |
| OdiMath6 | GSOIDP00016418001 | | Oikobase |
| OdiNeurogenin | GSOIDP00008211001 | | Oikobase |
| OloMath | SCLD01059283 | manual annotation | NCBI: ASM436789v1 |
| OloNeurogenin | SCLD01144392 | manual annotation | NCBI: ASM436789v1 |
| OloNeurogenin | SCLD01090848 | manual annotation | NCBI: ASM436789v1 |
| OvaMath | SCLH01000781 | manual annotation | NCBI: ASM436785v1 |
| OvaNeurogenin | SCLH01000494.1 | manual annotation | NCBI: ASM436785v1 |
| PfuMath | Phfumi.CG.MTP2014.S940.g03013.01.p | | Aniseed |
| PfuMesp | S15 | manual annotation | Aniseed |

| | | | |
|---|---|---|---|
| PfuNeurogenin | Phfumi.CG.MTP2014.S12979.g09049.01.p | | Aniseed |
| PmaMath | phmamm.CG.MTP2014.S497.g09451.02.p | | Aniseed |
| PmaMesp | S234 | manual annotation | Aniseed |
| PmaNeurogenenin | Phmamm.CG.MTP2014.S229.g05747.01.p | | Aniseed |

### Ets/Erg

| Gene name | Gene/Scaffold ID | | Database |
|---|---|---|---|
| BfErg | XP_002613111 | | NCBI |
| BflEts | XP_002610126.1 | | NCBI |
| BlaErg | BL14695_evm0 | | NCBI: Bl71nemr |
| BlaEts | 14693 | | NCBI: Bl71nemr |
| BleErga | S23.g05401.01 | | Aniseed |
| BleErgb | S391.g08928.01 | | Aniseed |
| BleErgc | S74 | manual annotation | Aniseed |
| BleEts1/2a | S422.g09525.01.p | | Aniseed |
| BleEts1/2b | S498.g10696.01.t | | Aniseed |
| BscErga | chr7.g26841.01.p | | Aniseed |
| BscErgb | chrUn.g52263.01 | | Aniseed |
| BscErgc | chrUn.g09892.01.t | | Aniseed |
| BscEts1/2a | ChrUn_p381405427 | manual annotation | Aniseed |
| BscEts1/2b | chr5.g69093.01.p | | Aniseed |
| BspEts1/2b2 | SCLE01158590 | manual annotation | NCBI: ASM436795v1 |
| BspEts1/2b1 | SCLE01307272.1 | manual annotation | NCBI: ASM436795v1 |
| CroErga | KH.C4.539.v1.A.ND1-1 | | Aniseed |
| CroErgb1 | KH.C10.148.v1.A.SL6-1 | | Aniseed |
| CroErgb2 | KH.C10.420.v1.A.SL2-1 | | Aniseed |
| CroErgc | KH.C2.290.v1.A.SL2-1 | | Aniseed |
| CroEts1/2a | KH.C10.113.v4.A.SL1-1 | | Aniseed |
| CroEts1/2b | KH.C11.10.v1.R.ND1-1 | | Aniseed |
| CsaErga | R15.1436501-1439403.13220 | manual annotation | Aniseed |
| CsaErgb1 | R44.172747-182062.06822 | manual annotation | Aniseed |
| CsaErgb2 | R44.150990-155653.06815 | manual annotation | Aniseed |
| CsaErgc | R.85.343670-348289.01847 | manual annotation | Aniseed |
| CsaEts1/2a | R44.821279-826601.11000.p | manual annotation | Aniseed |
| CsaEts1/2b | R16.345322-348455.08453.p | manual annotation | Aniseed |
| FboEts1/2b | SDII01087108 | manual annotation | NCBI: ASM436807v1 |
| GgaErg | NP_989611.1 | | NCBI |
| GgaEts1 | XP_015153454.1 | | NCBI |
| GgaEts2 | NP_990643.1 | | NCBI |
| HauErga | S207.g03311.01.p | | Aniseed |
| HauErgb | S806.g06729.01 | | Aniseed |
| HauErgc | S3595.g10346.01.p | | Aniseed |
| HauEts1/2a | S287 | manual annotation | Aniseed |
| HauEts1/2b | S120.g02312.01.p | | Aniseed |
| HroErga | S37.g15279.01.p | | Aniseed |
| HroErgb | S156.g00533.01 | | Aniseed |
| HroErgc | S414.g02267.01.p | | Aniseed |
| HroEts1/2a | S361 | manual annotation | Aniseed |
| HroEts1/2b | S7.g06548.01.p | | Aniseed |
| HsaErg | NP_891548.1 | | NCBI |
| HsaEts1 | NP_001137292.1 | | NCBI |
| HsaEts2 | NP_005230.1 | | NCBI |
| LchErg | XP_006002586.1 | | NCBI |
| LchEts1 | XP_005989874.1 | | NCBI |
| LchEts2 | XP_006002591.1 | | NCBI |
| LocErg | XP_006627839.1 | | NCBI |
| LocEts1 | XP_015192990.1 | | NCBI |
| LocEts2 | XP_015197244.1 | | NCBI |
| Mer9Ets1/2b2 | SCLF01704939 | manual annotation | NCBI:ASM436797v1 |
| MerEts1/2b1 | SCLF01279254.1 | manual annotation | NCBI:ASM436797v1 |
| MocciErga | S511319.g19317.01 | | Aniseed |
| MocciErgb | S244705.g05536.01.t | | Aniseed |
| MocciErgc | S79112.g01052.01 | | Aniseed |
| MocciEts1/2a | S632673.g27652.01.p | | Aniseed |
| MoccuErga | S269753.g10395.01 | | Aniseed |
| MoccuErgb | S197076.g06857.01.p | | Aniseed |
| MoccuErgc | S413423.g19312.01 | | Aniseed |
| MoccuEts1/2a | S711677 | manual annotation | Aniseed |
| MocuErgc | S60328.g03798.01 | | Aniseed |
| MoculErga | S51598.g02982.01 | | Aniseed |
| MoculErgb | S97310.g08238.01.t | | Aniseed |
| MoculEts1/2a | S99115.g08619.01.p | | Aniseed |
| OalEts1/2b1 | SCLG01000003 | manual annotation | NCBI: ASM436787v1 |
| OalEts1/2b2 | SCLG01000710.1 | manual annotation | NCBI: ASM436787v1 |
| OdiErg | GSOIDP00014411001_S53 | manual annotation | Oikobase |
| OdiEts1/2b2 | GSOIDP00003683001 | | Oikobase |
| OdiEts1/2b1 | GSOIDP00006792001_S212 | manual annotation | Oikobase |
| OloEts1/2b1 | SCL0D01110143 | manual annotation | NCBI: ASM436789v1 |
| OloEts1/2b2 | SCLD01120505.1 | manual annotation | NCBI: ASM436789v1 |
| OvaEts1/2b1 | SCLH01001059 | manual annotation | NCBI: ASM436785v1 |
| OvaEts1/2b2 | SCLH01001181.1 | manual annotation | NCBI: ASM436785v1 |
| PfuErga | S2966_S20668 | manual annotation | Aniseed |
| PfuErgb | S3340.g05634.01 | | Aniseed |
| PfuErgc | S1489.g03780.02.p | | Aniseed |
| PfuEts1/2a | S9396 | manual annotation | Aniseed |
| PfuEts1/2b | S357.g01840.01.p | | Aniseed |
| PmaErga | S491.g09369.01.p | | Aniseed |
| PmaErgb | S18.g00779.01 | | Aniseed |
| PmaErgc | S64.g02300.02.p | | Aniseed |
| PmaEts1/2a | S1111.g13716.02.p | | Aniseed |
| PmaEts1/2b | S484.g09279.01.p | | Aniseed |

### Mek

| Gene name | Gene/Scaffold ID | | Database |
|---|---|---|---|
| BbeMEK12 | XP_019619563.1 | | NCBI |
| BbeMEK36 | XP_019644388.1 | | NCBI |
| BbeMEK4 | XP_019642021.1 | | NCBI |
| BbeMEK5 | XP_019624095.1 | | NCBI |
| BbeMEK7 | XP_019643631.1 | | NCBI |
| Bfl_STKIN25 | XP_002598737.1 | | NCBI |
| Bfl_STKIN3 | XP_002589064.1 | | NCBI |
| BflMEK12 | XP_002601633.1 | | NCBI |
| BflMEK36 | XP_002595698.1 | | NCBI |
| BflMEK4 | XP_002607255.1 | | NCBI |
| BflMEK5 | XP_035666864.1 | | NCBI |
| BflMEK7 | XP_002589630.1 | | NCBI |
| BlaMEK12 | BL22940_cuf4 | manual annotation | NCBI: Bl71nemr |
| BlaMEK36 | BL19281_evm0 | manual annotation | NCBI: Bl71nemr |
| BlaMEK4 | BL05717_evm0 | manual annotation | NCBI: Bl71nemr |
| BlaMEK5 | BL03615_evm11 | manual annotation | NCBI: Bl71nemr |
| BlaMEK7 | BL23231_evm0 | manual annotation | NCBI: Bl71nemr |
| BleMEK12 | Boleac.CG.SB_v3.S442.g09834.01.p | | Aniseed |
| BleMEK36 | Boleac.CG.SB_v3.S25.g05985.01.p | | Aniseed |
| BleMEK4 | Boleac.CG.SB_v3.S88.g14953.01.p | | Aniseed |
| BleMEK5 | Boleac.CG.SB_v3.S34.g08001.01.p | | Aniseed |
| BleMEK7 | Boleac.CG.SB_v3.S100.g00402.01.p | | Aniseed |
| BscMEK12 | Boschl.CG.Botznik2013.chrUn.g28277.01.p | | Aniseed |
| BscMEK36 | chr2+chrUN | manual annotation | Aniseed |
| BscMEK4 | Boschl.CG.Botznik2013.chrUn.g03398.01.p+Boschl.CG.Botznik2013.chr5.g39649.01.p | manual annotation | Aniseed |
| BscMEK5 | Boschl.CG.Botznik2013.chr12.g61303.01.p | | Aniseed |
| BscMEK7 | Boschl.CG.Botznik2013.chrUn.g27076.01.p | | Aniseed |
| BspMEK36 | SCLE01439045.1+SCLE01416150.1+SCLE01459439.1 | manual annotation | NCBI: ASM436795v1 |
| BspMEK7 | SCLE01016762.1+SCLE01426643.1+SCLE01329307.1+SCLE01046941.1 | manual annotation | NCBI: ASM436795v1 |
| Cro_STKIN25 | KH.C7.308.v1.A.nonSL2-1 | | Aniseed |
| Cro_STKIN3 | KH.C5.341.v1.A.nonSL1-1 | | Aniseed |
| CroMEK12 | KH.L147.22.v1.A.SL1-1 | | Aniseed |
| CroMEK36 | KH.L172.1.v1.A.ND1-1 | | Aniseed |
| CroMEK4 | KH.C8.343.v1.A.SL2-1 | | Aniseed |
| CroMEK5 | KH.C11.84.v1.A.SL3-1 | | Aniseed |
| CroMEK7 | KH.L22.44.v1.A.SL2-1 | | Aniseed |
| CsaMEK12 | Cisavi.CG.ENS81.R0.1216963-1218983.14576.p | | Aniseed |
| CsaMEK36 | Cisavi.CG.ENS81.R905.3972-11331.06316.p | | Aniseed |
| CsaMEK4 | Cisavi.CG.ENS81.R134.411618-418460.08740.p | | Aniseed |
| CsaMEK5 | Cisavi.CG.ENS81.R16.1649016-1652382.14365.p | | Aniseed |
| CsaMEK7 | Cisavi.CG.ENS81.R58.2891242-2899577.17580.p | | Aniseed |
| FboMEK36a | SDII01028860.1 | manual annotation | NCBI: ASM436807v1 |
| FboMEK36b | SDII01028860.1 | manual annotation | NCBI: ASM436807v1 |
| GgaMEK1 | XP_015147582.1 | | NCBI |
| GgaMEK2 | XP_015155404.1 | | NCBI |
| GgaMEK3 | XP_025010848.1 | | NCBI |
| GgaMEK4 | XP_015150723.1 | | NCBI |
| GgaMEK5 | XP_015147584.1 | | NCBI |
| GgaMEK6 | XP_003642396.1 | | NCBI |
| GgaMEK7 | XP_025001730.1 | | NCBI |
| HauMEK12 | Haaura.CG.MTP2014.S88.g01839.02.p | | Aniseed |
| HauMEK36 | Haaura.CG.MTP2014.S1027.g07428.01.p | | Aniseed |
| HauMEK4 | Haaura.CG.MTP2014.S11.g00363.01.p+Haaura.CG.MTP2014.S11.g00364.01.p | manual annotation | Aniseed |
| HauMEK5 | Haaura.CG.MTP2014.S2757.g09855.01.p | | Aniseed |
| HauMEK7 | Haaura.CG.MTP2014.S228.g03495.01.p | | Aniseed |
| HroMEK12 | Harore.CG.MTP2014.S158.g12521.02.p | | Aniseed |
| HroMEK36 | Harore.CG.MTP2014.S42.g10988.01.p | | Aniseed |
| HroMEK4 | Harore.CG.MTP2014.S12.g12037.01.p+Harore.CG.MTP2014.S12.g10989.01.p | manual annotation | Aniseed |
| HroMEK5 | Harore.CG.MTP2014.S150.g06112.01.p | | Aniseed |
| HroMEK7 | Harore.CG.MTP2014.S21.g00956.01.p | | Aniseed |
| Hsa_STKIN25 | NP_001258906.1 | | NCBI |
| Hsa_STKIN3 | NP_006272.2 | | NCBI |
| HsaMEK1 | NP_002746.1 | | NCBI |
| HsaMEK2 | NP_109587.1 | | NCBI |
| HsaMEK3 | NP_659731.1 | | NCBI |
| HsaMEK4 | NP_003001.1 | | NCBI |
| HsaMEK5 | NP_002748.1 | | NCBI |
| HsaMEK6 | NP_002749.2 | | NCBI |
| HsaMEK7 | NP_001284485.1 | | NCBI |
| LchMEK1 | XP_005990160.1 | | NCBI |
| LchMEK2 | XP_005999133.1 | | NCBI |
| LchMEK3 | XP_006008481.1 | | NCBI |
| LchMEK4 | XP_006011477.1 | | NCBI |
| LchMEK5 | XP_006001847.1 | | NCBI |
| LchMEK6 | XP_005992212.2 | | NCBI |
| LchMEK7 | XP_014347656.1 | | NCBI |
| LocMEK1 | XP_006628796.1 | | NCBI |
| LocMEK2 | XP_006639937.1 | | NCBI |
| LocMEK3 | XP_015215390.1 | | NCBI |
| LocMEK4 | XP_006635184.2 | | NCBI |
| LocMEK5 | XP_006628877.1 | | NCBI |
| LocMEK6 | XP_006635234.1 | | NCBI |
| LocMEK7 | XP_015204544.1 | | NCBI |
| MerMEK36 | SCLF01578774.1+SCLF01664085.1+SCLF01097487.1+SCLF01627178.1 | manual annotation | NCBI:ASM436797v1 |
| MerMEK7 | SCLF01730825.1+SCLF01141466.1+SCLF01164915.1 | manual annotation | NCBI:ASM436797v1 |
| MoocciMEK12 | Moocci.CG.ELv1_2.S600390.g24657.01.p | | Aniseed |
| MoocciMEK36 | Moocci.CG.ELv1_2.S257470.g05970.01.p | | Aniseed |
| MoocciMEK4 | Moocci.CG.ELv1_2.S263366.g06185.01.pS301794 | | Aniseed |
| MoocciMEK5 | Moocci.CG.ELv1_2.S171171.g03276.01.p | | Aniseed |
| MoocciMEK7 | Moocci.CG.ELv1_2.S646071.g29530.01.p | | Aniseed |
| MooccuMEK12 | Mooccu.CG.ELv1_2.S441521.g21440.01.p+Mooccu.CG.ELv1_2.S558906.g34076.01.t+Mooccu.CG.ELv1_2.S550223.g32758.01.t | manual annotation | Aniseed |
| MooccuMEK36 | Mooccu.CG.ELv1_2.S540638.g31404.01.p | | Aniseed |
| MooccuMEK4 | Mooccu.CG.ELv1_2.S722400.g47409.01.p | | Aniseed |
| MooccuMEK5 | Mooccu.CG.ELv1_2.S721609.g47275.01.p | | Aniseed |

| | | | |
|---|---|---|---|
| MooccuMEK7 | Mooccu.CG.ELv1_2.S703145.g44956.01.p+S703145+S665598+S703145+Mooccu.CG.ELv1_2.S703145.g44957.01.p | manual annotation | Aniseed |
| MooculMEK12 | Moocul.CG.ELv1_2.S91994.g07329.01.p | | Aniseed |
| MooculMEK36 | Moocul.CG.ELv1_2.S20893.g00805.01.p | | Aniseed |
| MooculMEK4 | Moocul.CG.ELv1_2.S31823.g01546.01.p | | Aniseed |
| MooculMEK5 | Moocul.CG.ELv1_2.S110643.g11702.01.p | | Aniseed |
| MooculMEK7 | Moocul.CG.ELv1_2.S80168.g05886.01.p | | Aniseed |
| OalMEK36 | SCLG01000032.1 | manual annotation | NCBI: ASM436787v1 |
| OalMEK7a | SCLG01000544.1 | manual annotation | NCBI: ASM436787v1 |
| Odi_STKIN25 | GSOIDP00001006001 | | Oikobase |
| Odi_STKIN4 | GSOIDP00013245001 | | Oikobase |
| OdiMEK36 | GSOIDP00015053001 | | Oikobase |
| OdiMEK7 | GSOIDP00010943001+Sc37_231336-231910 | manual annotation | Oikobase |
| OloMEK36a | SCLD01163008.1 | manual annotation | NCBI: ASM436789v1 |
| OloMEK36b | SCLD01098264.1 | manual annotation | NCBI: ASM436789v1 |
| OloMEK36c | SCLD01109975.1 | manual annotation | NCBI: ASM436789v1 |
| OloMEK7a | SCLD01093505.1 | manual annotation | NCBI: ASM436789v1 |
| OloMEK7b | SCLD01178439.1 | manual annotation | NCBI: ASM436789v1 |
| OvaMEK36 | SCLH01000769.1 | manual annotation | NCBI: ASM436785v1 |
| OvaMEK7 | SCLH01022748.1+SCLH01001128.1+SCLH01045676.1 | manual annotation | NCBI: ASM436785v1 |
| PfuMEK12 | Phfumi.CG.MTP2014.S42.g00590.01.p | manual annotation | Aniseed |
| PfuMEK36 | Phfumi.CG.MTP2014.S223.g01451.01.p | | Aniseed |
| PfuMEK4 | Phfumi.CG.MTP2014.S1108.g03276.01.p | | Aniseed |
| PfuMEK5 | Phfumi.CG.MTP2014.S835.g02829.01.p | | Aniseed |
| PfuMEK7 | Phfumi.CG.MTP2014.S2409.g04821.01.p | | Aniseed |
| PmaMEK12 | Phmamm.CG.MTP2014.S3266.g17598.01.p | | Aniseed |
| PmaMEK36 | Phmamm.CG.MTP2014.S195.g05134.01.p | | Aniseed |
| PmaMEK4 | Phmamm.CG.MTP2014.S12.g00494.01.p | | Aniseed |
| PmaMEK5 | Phmamm.CG.MTP2014.S469.g09085.01.p | | Aniseed |
| PmaMEK7 | Phmamm.CG.MTP2014.S179.g04836.02.p | | Aniseed |

### Gata

| Gene name | Gene/Scaffold ID | | Database |
|---|---|---|---|
| BbeGata123 | XP_019627141.1 | | NCBI |
| BbeGata456 | XP_019618792.1 | | NCBI |
| BflGTF123 | ACR66214.1 | | NCBI |
| BflGTF456a | ACR66216.1 | | NCBI |
| BlaGata123 | AFJ79490.1 | | NCBI |
| BlaGata456 | AFJ79491.1 | | NCBI |
| BleGata123 | S314 | manual annotation | Aniseed |
| BleGata456 | Boleac.CG.SB_v3.S147.g02878.01.p | | Aniseed |
| BscGata123 | ChrUn_Chr10 | manual annotation | Aniseed |
| BscGata123 | Chr10 | manual annotation | Aniseed |
| BscGata456 | ChrUn | manual annotation | Aniseed |
| BspGata | SCLE01420081 | manual annotation | NCBI: ASM436795v1 |
| BspGata | SCLE01408957 | manual annotation | NCBI: ASM436795v1 |
| BspGata | SCLE01414813 | manual annotation | NCBI: ASM436795v1 |
| BspGata | SCLE01211222 | manual annotation | NCBI: ASM436795v1 |
| CroGata123 | KH.S696.1.v1.R.ND1_1 | | Aniseed |
| CroGata456 | KH.L20.1.v1.A.SL2-1 | | Aniseed |
| CsaGata123 | Cisavi.CG.ENS81.R41.2121992-2129284.14153.p | | Aniseed |
| CsaGata456 | Cisavi.CG.ENS81.R77.2996926-3008818.14416.p | | Aniseed |
| FboGata | SDII01097846 | manual annotation | NCBI: ASM436807v1 |
| FboGata | SDII01078871 | manual annotation | NCBI: ASM436807v1 |
| GgaGata1 | NP_990795.1 | | NCBI |
| GgaGata2 | NP_001003797.1 | | NCBI |
| GgaGata3 | NP_001008444.1 | | NCBI |
| GgaGata4 | NP_001280035.1 | | NCBI |
| GgaGata5 | NP_990752.2 | | NCBI |
| GgaGata6 | NP_990751.1 | | NCBI |
| HauGata123 | Haaura.CG.MTP2014.S24.g00692.01.p_S24 | | Aniseed |
| HauGata456 | Haaura.CG.MTP2014.S112.g02206.01.p | | Aniseed |
| HroGata123 | Harore.CG.MTP2014.S12.g11731.01.t_S12 | | Aniseed |
| HroGata456 | Harore.CG.MTP2014.S45.g04355.01.p | | Aniseed |
| HsaGata1 | NP_002040.1 | | NCBI |
| HsaGata2 | NP_001139133.1 | | NCBI |
| HsaGata3 | NP_001002295.1 | | NCBI |
| HsaGata4 | NP_001295022.1 | | NCBI |
| HsaGata5 | NP_536721.1 | | NCBI |
| HsaGata6 | NP_005248.2 | | NCBI |
| LchGata1 | XP_006008598.1 | | NCBI |
| LchGata2 | XP_014341892.1 | | NCBI |
| LchGata3 | XP_005993937.1 | | NCBI |
| LchGata4 | XP_014351854.1 | | NCBI |
| LchGata5 | XP_014344220.1 | | NCBI |
| LchGata6 | XP_005995770.1 | | NCBI |
| LocGata1 | XP_006625356.2 | | NCBI |
| LocGata2 | XP_006631129.1 | | NCBI |
| LocGata3 | XP_006633355.1 | | NCBI |
| LocGata4 | XP_006642996.1 | | NCBI |
| LocGata5 | XP_015220493.1 | | NCBI |
| LocGata6 | XP_006633923.1 | | NCBI |
| MerGata | SCLF01270895 | manual annotation | NCBI:ASM436797v1 |
| MerGata | SCLF01415120 | manual annotation | NCBI:ASM436797v1 |
| MerGata | SCLF01211403_1 | manual annotation | NCBI:ASM436797v1 |
| MerGata | SCLF01530214 | manual annotation | NCBI:ASM436797v1 |
| MerGata | SCLF01239871 | manual annotation | NCBI:ASM436797v1 |
| MocciGata123 | S487156 | manual annotation | Aniseed |
| MocciGata456 | S458966 | manual annotation | Aniseed |
| MoccuGata123 | S723572 | manual annotation | Aniseed |
| MoccuGata456 | S557110_S209818 | manual annotation | Aniseed |
| MoculGata123 | S97234 | manual annotation | Aniseed |
| MoculGata456 | Moocul.CG.ELv1_2.S44512.g02355.01.p | | Aniseed |
| OalGata | SCLG01000167 | manual annotation | NCBI: ASM436787v1 |
| OalGata | SCLG010000323 | manual annotation | NCBI: ASM436787v1 |

| | | | |
|---|---|---|---|
| OdGata1/2/3c | GSOIDP00008593001 | | Oikobase |
| OdGata1/2/3d | GSOIDP00000004001 | | Oikobase |
| OdGata1/2/3b | GSOIDP00011092001 | | Oikobase |
| OdGata1/2/3a | GSOIDP00007921001 | | Oikobase |
| OloGata | SCLD01106159 | manual annotation | NCBI: ASM436789v1 |
| OloGata | SCLD01172774 | manual annotation | NCBI: ASM436789v1 |
| OloGata | SCLD01119148 | manual annotation | NCBI: ASM436789v1 |
| OloGata | SCLD01126560 | manual annotation | NCBI: ASM436789v1 |
| OloGata | SCLD01018184 | manual annotation | NCBI: ASM436789v1 |
| OvaGata | SCLH01000841 | manual annotation | NCBI: ASM436785v1 |
| OvaGata | SCLH01000110 | manual annotation | NCBI: ASM436785v1 |
| OvaGata | SCLH01000261 | manual annotation | NCBI: ASM436785v1 |
| OvaGata | SCLH01002033_F | manual annotation | NCBI: ASM436785v1 |
| OvaGata | SCLH01002033_R | manual annotation | NCBI: ASM436785v1 |
| PfuGata123 | S244_S11342 | manual annotation | Aniseed |
| PfuGata456 | S6271_0_S2222 | manual annotation | Aniseed |
| PmaGata123 | S535 | manual annotation | Aniseed |
| PmaGata456 | Phmamm.CG.MTP2014.S835.g12260.01.p | | Aniseed |

### FoxF

| Gene name | Gene/Scaffold ID | | Database |
|---|---|---|---|
| BbeFoxF | XP_019628964.1 | | NCBI |
| BbeFoxQ | XP_019628968.1 | | NCBI |
| BflFoxF | CAH69695.1 | | NCBI |
| BflFoxQ | XP_035680367.1 | | NCBI |
| BlaFoxF | BL02311_evm2 | manual annotation | NCBI: Bl71nemr |
| BlaFoxQ | BL00713_evm0 | manual annotation | NCBI: Bl71nemr |
| BleFoxF | S21 | manual annotation | Aniseed |
| BscFoxF | Chr9_ChrUn | manual annotation | Aniseed |
| BspFoxF | SCLE01409615 | | NCBI: ASM436795v1 |
| CrFoxQ | KH.L44.19.v1.A.nonSL4-1 | | Aniseed |
| CroFoxF | KH.C3.170.v1.A.SL1-1 | | Aniseed |
| CsaFoxF | Cisavi.CG.ENS81.R11.1287035-1294060.15471.p | | Aniseed |
| CsaFoxQ | Cisavi.CG.ENS81.R1.1226140-1229479.11615.p | | Aniseed |
| FboFoxF | SDII01127328 | manual annotation | NCBI: ASM436807v1 |
| GgaFoxF1 | XP_414186.5 | | NCBI |
| GgaFoxF2 | XP_015137672.2 | | NCBI |
| GgaFoxQ | XP_015137671.2 | | NCBI |
| HauFoxF | Haaura.CG.MTP2014.S882.g06990.01.p | | Aniseed |
| HroFoxF | Harore.CG.MTP2014.S48.g08804.01.p | | Aniseed |
| HsaFoxF1 | NP_001442.2 | | NCBI |
| HsaFoxF2 | NP_001443.1 | | NCBI |
| HsFoxQ1 | NP_150285.3 | | NCBI |
| LchFoxF1 | XP_005995947.1 | | NCBI |
| LchFoxF2 | XP_006001190.1 | | NCBI |
| LchFoxQ | XP_006013316.1 | | NCBI |
| LocFoxF1 | XP_006641203.1 | | NCBI |
| LocFoxF2 | XP_006634525.1 | | NCBI |
| LocFoxQ | XP_006634621.1 | | NCBI |
| MerFoxF | SCLF01086355 | manual annotation | NCBI:ASM436797v1 |
| MocciFox | S456097 | manual annotation | Aniseed |
| MoccuFoxF | S685721 | manual annotation | Aniseed |
| MoculFoxF | Moocul.CG.ELv1_2.S114506.g13471.01.p | | Aniseed |
| OalFoxF | SCLG01000319 | manual annotation | NCBI: ASM436787v1 |
| OdFoxQ | GSOIDP00009755001 | | Oikobase |
| OdiFoxF | GSOIDP00007763001 | | Oikobase |
| OloFox | SCLD01163618 | manual annotation | NCBI: ASM436789v1 |
| OvaFoxF | SCLH01001792 | manual annotation | NCBI: ASM436785v1 |
| PfuFoxF | Phfumi.CG.MTP2014.S1012.g03122.01.p | | Aniseed |
| PfuFoxQ | Phfumi.CG.MTP2014.S3505.g05754.01.p | | Aniseed |
| PmaFoxF | S371 | manual annotation | Aniseed |
| PmaFoxQ | Phmamm.CG.MTP2014.S654.g10956.01.p | | Aniseed |

## Nk

| Gene name | Gene/Scaffold ID | | Database |
|---|---|---|---|
| BbeNkx2.1 | XP_019648010.1 | | NCBI |
| BbeNkx2.2 | XP_019647989.1 | | NCBI |
| BbeNkx4 | XP_019646148.1 | | NCBI |
| BflNkx2.1 | XP_002589194.1 | | NCBI |
| BflNkx2.2 | XP_002589199.1 | | NCBI |
| BflNkx4 | XP_002610435.1 | | NCBI |
| BlaNkx4 | AWV91585.1 | | NCBI |
| BleNk4 | S7 | | aniseed |
| BleNk2 | S959 | | aniseed |
| BscNk4 | chr9 | | aniseed |
| BscNk2 | ChrUn | | aniseed |
| BspNk4_5'long | SCLE01112255.1 | manual annotation | NCBI: ASM436795v1 |
| CroNK2a | KH.C10.338.v1.A.SL4 | | aniseed |
| CroNk4 | KH.C8.482.v1.A.SL2-1 | | aniseed |
| CsaNK2 | ENS81.R37.1525232 | | aniseed |
| CsaNk4 | Cisavi.CG.ENS81.R24.410826-416079.10470.p | | aniseed |
| FboNk2 | SDII1122225 | manual annotation | NCBI: ASM436807v1 |
| FboNk2 | SDII01045043 | manual annotation | NCBI: ASM436807v1 |
| GgaNkx2.1 | NP_989947.1 | | NCBI |
| GgaNkx2.2 | NP_001264647.1 | | NCBI |
| GgaNkx2.3 | NP_990362.2 | | NCBI |
| GgaNkx2.4 | XP_015138916.1 | | NCBI |
| GgaNkx2.5 | NP_990495.1 | | NCBI |
| GgaNkx2.6 | NP_990468.1 | | NCBI |
| GgaNkx2.8 | XP_003641408.3 | | NCBI |

| | | | |
|---|---|---|---|
| HauNk4 | S249RC_S5459 | manual annotation | aniseed |
| HauNk2 | Haaura.CG.MTP2014.S265.g03837.01.p | | aniseed |
| HroNk4 | S41_333208-368464 | manual annotation | aniseed |
| HroNk2 | Harore.CG.MTP2014.S132.g01719.01.p | | aniseed |
| HsaNk2.4 | NP_149416.1 | | NCBI |
| HsaNk2.6 | NP_001129743.2 | | NCBI |
| HsaNKx2.2 | NP_002500.1 | | NCBI |
| HsaNkx2.3 | NP_660328.2 | | NCBI |
| HsaNkx2.5 | NP_004378.1 | | NCBI |
| LchNkx2.1 | XP_005989231.1 | | NCBI |
| LchNkx2.2 | XP_006013284.1 | | NCBI |
| LchNkx2.3 | XP_006001881.1 | | NCBI |
| LchNkx2.4 | XP_006008054.1 | | NCBI |
| LchNkx2.5 | XP_005992571.1 | | NCBI |
| LchNkx2.6 | XP_005991959.1 | | NCBI |
| LchNkx2.8 | XP_005989262.1 | | NCBI |
| LocNkx2.1 | XP_006632731.1 | | NCBI |
| LocNkx2.2 | XP_006626197.1 | | NCBI |
| LocNkx2.3 | XP_006630496.1 | | NCBI |
| LocNkx2.4 | XP_015208466.1 | | NCBI |
| LocNkx2.5 | XP_006631743.1 | | NCBI |
| LocNkx2.6 | XP_006625550.1 | | NCBI |
| LocNkx2.8 | XP_006632762.2 | | NCBI |
| MerNk4_5'long | SCLF01095022.1 | manual annotation | NCBI: ASM436797v1 |
| MocciNk2 | S77334 | manual annotation | aniseed |
| MocciNk4 | S332293 | manual annotation | aniseed |
| MoccuNk2 | S690034 | manual annotation | aniseed |
| MoccuNk4 | Mooccu.CG.ELv1_2.S657543.g40314.01.p | | aniseed |
| MoculNk2 | S113495 | manual annotation | aniseed |
| MoculNk4 | Moocul.CG.ELv1_2.S112088.g12247.01.p_modify | | aniseed |
| OalNk4_5'long | SCLG01000524.1 | manual annotation | NCBI: ASM436787v1 |
| OalNk2 | SCLG01000088 | manual annotation | NCBI: ASM436787v1 |
| OdiNk4_BCN_5long | | manual annotation | |
| OdiNK2a | GSOIDP00010368001 | | oikobase |
| OdiNK2b | GSOIDP00011992001 | | oikobase |
| OloNk4_5'long | SCLD01104089.1 | manual annotation | NCBI: ASM436789v1 |
| OvaNk4_5'long | SCLH01002003.1 | manual annotation | NCBI: ASM436785v1 |
| OvaNk2b | SCLH01000727 | manual annotation | NCBI: ASM436785v1 |
| PfuNk2 | Phfumi.CG.MTP2014.S953.g03028.02.p | | aniseed |
| PfuNk4 | S1495 | manual annotation | aniseed |
| PmaNk2 | Phmamm.CG.MTP2014.S597.g10419.03.p | | aniseed |
| PmaNk4 | S185 | manual annotation | aniseed |

### Hand

| Gene name | Gene/Scaffold ID | | Database |
|---|---|---|---|
| BbeHand | XP_019625359.1 | | NCBI |
| BflHand | AEL13770.1 | | NCBI |
| BlaHand | AFJ79493.1 | | NCBI |
| BleHandr | S96 | manual annotation | Aniseed |
| BleHand | Boleac.CG.SB_v3.S154.g03139.01.p | | Aniseed |
| BscHand | ChrUn | manual annotation | Aniseed |
| BscHandr | Boschl.CG.Botznik2013.chr12.g10820.01.p | | Aniseed |
| BspHand | SCLE01437072 | manual annotation | NCBI: ASM436795v1 |
| CroHand | KH.C14.604.v1.A.SL2_1 | | Aniseed |
| CroHandr | KH.C1.1116.v1.A.SL1_1 | | Aniseed |
| CsaHandr | Cisavi.CG.ENS81.R14.220962-222552.16137.p | | Aniseed |
| CsaHand | Cisavi.CG.ENS81.R49.2261498-2262994.16124.p | | Aniseed |
| GgaHand1 | NP_990296.1 | | NCBI |
| GgaHand2 | NP_990297.2 | | NCBI |
| HauHand | Haaura.CG.MTP2014.S167.g02904.01.p | | Aniseed |
| HauHandr | Haaura.CG.MTP2014.S72.g01613.01.p | | Aniseed |
| HroHand | Harore.CG.MTP2014.S40.g09177.01.p | | Aniseed |
| HroHandr | Harore.CG.MTP2014.S74.g11434.01.p | | Aniseed |
| HsaHand1 | NP_004812.1 | | NCBI |
| HsaHand2 | NP_068808.1 | | NCBI |
| LchHand1 | XP_005988374.1 | | NCBI |
| LchHand2 | XP_005993037.1 | | NCBI |
| LocHand1 | XP_006632012.1 | | NCBI |
| LocHand2 | XP_006630021.1 | | NCBI |
| MerHand | SCLF01004291_SCLF01727318_SCLF01160949 | manual annotation | NCBI:ASM436797v1 |
| MocciHandr | Moocci.CG.ELv1_2.S624793.g26726.01.p | | Aniseed |
| MoccuHandr | Mooccu.CG.ELv1_2.S674203.g41775.01.p | | Aniseed |
| MoculHandr | Moocul.CG.ELv1_2.S82437.g06150.01.p | | Aniseed |
| MocciHand | S492667 | manual annotation | Aniseed |
| MooculHand | Mooccu.CG.ELv1_2.S546820.g32354.01.p | | Aniseed |
| MooculHand | S112757 | manual annotation | Aniseed |
| OalHand | SCLG01000563 | manual annotation | NCBI: ASM436787v1 |
| OdiHand | GSOIDP00005552001 | | Oikobase |
| OloHand | SCLD01012489_SCLD01051243 | manual annotation | NCBI: ASM436789v1 |
| OloHand | SCLD01091277 | manual annotation | NCBI: ASM436789v1 |
| OvaHand | SCLH01000568 | manual annotation | NCBI: ASM436785v1 |
| PfuHand | Phfumi.CG.MTP2014.S2952.g05296.01.p | | Aniseed |
| PfuHandr | S5260 | manual annotation | Aniseed |
| PmaHandr | Phmamm.CG.MTP2014.S4.g00171.02.p | | Aniseed |
| PmaHand | Phmamm.CG.MTP2014.S2382.g16605.01.p | | Aniseed |

### Tbx

| Gene name | Gene/Scaffold ID | | Database |
|---|---|---|---|
| BflBrachyury1 | XP_002590391.1 | | NCBI |
| BflBrachyury2 | XP_002590390.1 | | NCBI |

| | | | |
|---|---|---|---|
| BflTbr1 | AAG34893.2 | | NCBI |
| BflTbx1/10 | AAG34887.2 | | NCBI |
| BflTbx15 | XP_002599057.1 | | NCBI |
| BflTbx2_3 | XP_002598922.1 | | NCBI |
| BflTbx20 | XP_002612350.1 | | NCBI |
| BflTbx4_5 | ABU50779.1 | | NCBI |
| BflTbx6_16 | XP_002612013.1 | | NCBI |
| BlaBrachyury | BL23557_evm0 | manual annotation | NCBI: Bl71nemr |
| BlaTbr1 | BL08925 | manual annotation | NCBI: Bl71nemr |
| BlaTbx1/10 | BL01988_evm1 | manual annotation | NCBI: Bl71nemr |
| BlaTbx15 | BL95766_evm0 | manual annotation | NCBI: Bl71nemr |
| BlaTbx2_3 | BL12502_BL02331_BL1275 | manual annotation | NCBI: Bl71nemr |
| BlaTbx20 | BL16028_evm0 | manual annotation | NCBI: Bl71nemr |
| BlaTbx4_5 | BL06379_evm0 | manual annotation | NCBI: Bl71nemr |
| BlaTbx6_16 | BL04677_cuf1 | manual annotation | NCBI: Bl71nemr |
| BleTbx1/10 | S60.g12196.01 | | Aniseed |
| BleTbx15/18/22 | S749.g13814.01 | | Aniseed |
| BleTbx19/Brachyury | S38.g08688.01 | | Aniseed |
| BleTbx2/3 | S553.g11558.01 | | Aniseed |
| BleTbx20 | S562.g11653.01 | | Aniseed |
| BleTbx6 | S256.g06109.01 | | Aniseed |
| BscTbx1/10 | chrUn_g48325.01 | | Aniseed |
| BscTbx19/Brachyury | chr7.g63408.01 | | Aniseed |
| BscTbx2/3 | chr7.g53212.01 | | Aniseed |
| BscTbx6 | chr1.g48703.01 | | Aniseed |
| BspBrachyury/Tbx19 | SCLE01228003.1 | manual annotation | NCBI: ASM436795v1 |
| BspTbx15/18/22 | SCLE01017494.1 | manual annotation | NCBI: ASM436795v1 |
| BspTbx2/3 | SCLE01446362.1 | manual annotation | NCBI: ASM436795v1 |
| BspTbx2/3 | SCLE01429136.1 | manual annotation | NCBI: ASM436795v1 |
| BspTbx6/16/24/VegT | SCLE01434700.1 | manual annotation | NCBI: ASM436795v1 |
| CroEomes/Tbr1 | KH.C3.773 | | Aniseed |
| CroTbx1/10 | KH.C7.628 | | Aniseed |
| CroTbx15/18/22 | KH.S589.4 | | Aniseed |
| CroTbx19/Brachyury | KH.S1404.1 | | Aniseed |
| CroTbx2/3 | KH.L96.87 | | Aniseed |
| CroTbx20 | KH.C1.224 | | Aniseed |
| CroTbx6.1 | KH.S654.1 | | Aniseed |
| CroTbx6.2 | KH.S654.2 | | Aniseed |
| CroTbx6.3 | KH.S654.3 | | Aniseed |
| CsaEomes/Tbr1 | R107.324117-331918.03246 | | Aniseed |
| CsaTbx1/10 | R125.93093-108455.01412 | | Aniseed |
| CsaTbx15/18/22 | R118.128673-1340382 | | Aniseed |
| CsaTbx19/Brachyury | R388.258080-264338.03855 | | Aniseed |
| CsaTbx2/3 | R90.318198-331613.16578 | | Aniseed |
| CsaTbx20 | R125.93093-108455.01412 | | Aniseed |
| CsaTbx6.1 | R16.5012720-5014970.20043 | | Aniseed |
| CsaTbx6.1 | R16.5041500-5055627.20057 | | Aniseed |
| FboBrachyury/Tbx19 | SDII01062704.1 | manual annotation | NCBI: ASM436807v1 |
| FboTbx15/18/22 | SDII01116622.1 | manual annotation | NCBI: ASM436807v1 |
| FboTbx2/3 | SDII01124641.1 | manual annotation | NCBI: ASM436807v1 |
| FboTbx20 | SCII01131462.1 | manual annotation | NCBI: ASM436807v1 |
| GgaBrachyury | XP_015139531.2 | | NCBI |
| GgaEomes | NP_001308484.1 | | NCBI |
| GgaTbr1 | XP_015145201.1 | | NCBI |
| GgaTbx1 | XP_025011481.1 | | NCBI |
| GgaTbx15 | XP_416537.3 | | NCBI |
| GgaTbx18 | NP_989784.2 | | NCBI |
| GgaTbx19 | NP_990281.1 | | NCBI |
| GgaTbx2 | XP_001235321.4 | | NCBI |
| GgaTbx20 | NP_989475.1 | | NCBI |
| GgaTbx21 | XP_015154966.2 | | NCBI |
| GgaTbx22 | NP_989437.1 | | NCBI |
| GgaTbx3 | NP_001257807.1 | | NCBI |
| GgaTbx4 | NP_001025708.1 | | NCBI |
| GgaTbx5 | XP_015150317.1 | | NCBI |
| GgaTbx6 | XP_025011193.1 | | NCBI |
| HauEomes/Tbr1 | S1059.g07498.01 | | aniseed |
| HauTbx15/18/22 | S4943.g10821.01 | | aniseed |
| HauTbx19/Brachyury | S324.g04307.01 | | aniseed |
| HauTbx2/3 | S9.g00298.01 | | aniseed |
| HauTbx6.1 | S829.g06809.03 | | aniseed |
| HauTbx6.2 | S1817.g08930.01 | | aniseed |
| HroEomes/Tbr1 | S78.g14749.01 | | aniseed |
| HroTbx15/18/22 | S61.g03659.01 | | aniseed |
| HroTbx19/Brachyury | S42.g07916.02 | | aniseed |
| HroTbx2/3 | S120.g12743.01 | | aniseed |
| HroTbx6.1 | S933.g10799.03 | | aniseed |
| HroTbx6.2 | S461.g13082.01 | | aniseed |
| HsaBrachyury | NP_001257413.1 | | NCBI |
| HsaEomes | NP_001265111.1 | | NCBI |
| HsaTbr1 | NP_006584.1 | | NCBI |
| HsaTbx1 | NP_005983.1 | | NCBI |
| HsaTbx15 | NP_689593.2 | | NCBI |
| HsaTbx18 | NP_001073977.1 | | NCBI |
| HsaTbx19 | NP_005140.1 | | NCBI |
| HsaTbx2 | NP_005985.3 | | NCBI |
| HsaTbx20 | NP_001159692.1 | | NCBI |
| HsaTbx21 | NP_037483.1 | | NCBI |
| HsaTbx22 | NP_058650.1 | | NCBI |
| HsaTbx6 | NP_004599.2 | | NCBI |
| HsTbx10 | NP_005986.2 | | NCBI |
| HsTbx3 | NP_005987.3 | | NCBI |
| HsTbx4 | NP_001308049.1 | | NCBI |
| HsTbx5 | NP_000183.2 | | NCBI |
| LchBrachyury | XP_005993410.1 | | NCBI |
| LchEomes | XP_006011925.1 | | NCBI |
| LchTbr1 | XP_006010469.1 | | NCBI |
| LchTbx1 | XP_014340634.1 | | NCBI |
| LchTbx15 | XP_014344889.1 | | NCBI |

| | | | |
|---|---|---|---|
| LchTbx18 | XP_005995114.1 | | NCBI |
| LchTbx19 | XP_014346004.1 | | NCBI |
| LchTbx2 | XP_005986878.1 | | NCBI |
| LchTbx20 | XP_006005638.1 | | NCBI |
| LchTbx21 | XP_005990126.1 | | NCBI |
| LchTbx22 | XP_014347604.1 | | NCBI |
| LchTbx3 | XP_006007893.1 | | NCBI |
| LchTbx4 | XP_005986877.1 | | NCBI |
| LchTbx5 | XP_006007890.1 | | NCBI |
| LchTbx6 | XP_014344551.1 | | NCBI |
| LocBrachyury | XP_015218241.1 | | NCBI |
| LocEomes | XP_006635610.1 | | NCBI |
| LocTbr1 | XP_006636611.1 | | NCBI |
| LocTbx1 | XP_006640182.1 | | NCBI |
| LocTbx15 | XP_006639276.1 | | NCBI |
| LocTbx18 | XP_006625898.1 | | NCBI |
| LocTbx19 | XP_015216992.1 | | NCBI |
| LocTbx2 | XP_006640960.1 | | NCBI |
| LocTbx20 | XP_006635615.1 | | NCBI |
| LocTbx21 | XP_015217756.1 | | NCBI |
| LocTbx22 | XP_006633032.1 | | NCBI |
| LocTbx3 | XP_006640338.1 | | NCBI |
| LocTbx4 | XP_015222868.1 | | NCBI |
| LocTbx5 | XP_006640339.1 | | NCBI |
| LocTbx6 | XP_006641063.1 | | NCBI |
| MerBrachyury/Tbx19 | SCLF01152421.1 | manual annotation | NCBI:ASM436797v1 |
| MerMod_Tbx2/3 | SCLF01056641.1 | manual annotation | NCBI:ASM436797v1 |
| MerTbx15/18/22 | SCLF01251203.1 | manual annotation | NCBI:ASM436797v1 |
| MerTbx2/3 | SCLF01026372.1 | manual annotation | NCBI:ASM436797v1 |
| MerTbx2/3 | SCLF01048308.1 | manual annotation | NCBI:ASM436797v1 |
| MerTbx6/16/24/VegT | SCLF01036633.1 | manual annotation | NCBI:ASM436797v1 |
| MocciEomes/Tbr1 | S477916.g16708.01 | | aniseed |
| MocciTbx15/18/22 | S530819.g20232.01 | | aniseed |
| MocciTbx19/Brachyury | S634979.g27904.01 | | aniseed |
| MocciTbx2/3 | S479065.g16818.01 | | aniseed |
| MocciTbx6 | S497697.g18857.01 | | aniseed |
| MoccuEomes/Tbr1 | S716823.g46618.01 | | aniseed |
| MoccuTbx15/18/22 | S640811.g39040.01 | | aniseed |
| MoccuTbx19/Brachyury | S705330.g45232.01 | | aniseed |
| MoculEomes/Tbr1 | 33300.g01638.01 | | aniseed |
| MoculTbx15/18/22 | S109292.g11175.01 | | aniseed |
| MoculTbx19/Brachyury | S90886.g07224.01 | | aniseed |
| MoculTbx2/3 | S99115.g08615.01 | | aniseed |
| MoculTbx6 | S69504.g04594.01 | | aniseed |
| OalBrachyury/Tbx19 | SCLG01000117.1 | manual annotation | NCBI: ASM436787v1 |
| OalTbx15/18/22 | SCLG01000201.1 | manual annotation | NCBI: ASM436787v1 |
| OalTbx2/3 | SCLG01022965.1 | manual annotation | NCBI: ASM436787v1 |
| OalTbx2/3 | SCLG01000157.1 | manual annotation | NCBI: ASM436787v1 |
| OalTbx2/3 | SCLG01000314.1 | manual annotation | NCBI: ASM436787v1 |
| OalTbx2/3 | SCLG01000339.1 | manual annotation | NCBI: ASM436787v1 |
| OalTbx6/16/24/VegT | SCLG01000014.1 | manual annotation | NCBI: ASM436787v1 |
| OdiBrachyury/Tbx19 | GSOIDP00000279001 | | Oikobase |
| OdiTbx15/18/22 | GSOIDP00014261001 | | Oikobase |
| OdiTbx15/18/22 | GSOIDP00014264001 | | Oikobase |
| OdiTbx2/3 | GSOIDP00004289001 | | Oikobase |
| OdiTbx2/3 | GSOIDP00011582001 | | Oikobase |
| OdiTbx2/3 | GSOIDP00004999001 | | Oikobase |
| OdiTbx2/3 | GSOIDP00007455001 | | Oikobase |
| OdiTbx6/16/24/VegT | GSOIDP00006159001 | | Oikobase |
| OloBrachyury/Tbx19 | SCLD01093191.1 | manual annotation | NCBI: ASM436789v1 |
| OloTbx15/18/22 | SCLD01119088.1 | manual annotation | NCBI: ASM436789v1 |
| OloTbx15/18/22 | SCLD01116716.1 | manual annotation | NCBI: ASM436789v1 |
| OloTbx2/3 | SCLD01098799.1 | manual annotation | NCBI: ASM436789v1 |
| OloTbx2/3 | SCLD01094440.1 | manual annotation | NCBI: ASM436789v1 |
| OloTbx2/3 | SCLD01116750.1 | manual annotation | NCBI: ASM436789v1 |
| OloTbx2/3 | SCLD01177323.1 | manual annotation | NCBI: ASM436789v1 |
| OloTbx6/16/24/VegT | SCLD01038142.1 | manual annotation | NCBI: ASM436789v1 |
| OvaBrachyury/Tbx19 | SCLH01001620.1 | manual annotation | NCBI: ASM436785v1 |
| OvaTbx15/18/22 | SCLH01000284.1 | manual annotation | NCBI: ASM436785v1 |
| OvaTbx2/3 | SCLH01000456.1 | manual annotation | NCBI: ASM436785v1 |
| OvaTbx2/3 | SCLH01000169.1 | manual annotation | NCBI: ASM436785v1 |
| OvaTbx2/3 | SCLH01000635.1 | manual annotation | NCBI: ASM436785v1 |
| OvaTbx2/3 | SCLH01000589.1 | manual annotation | NCBI: ASM436785v1 |
| OvaTbx6/16/24/VegT | SCLH01000086.1 | manual annotation | NCBI: ASM436785v1 |
| PfuEomes/Tbr1 | S70.g00816.01 | | aniseed |
| PfuTbx1/10 | S5861.g07133.01 | | aniseed |
| PfuTbx19/Brachyury | S1283.g03521.01 | | aniseed |
| PfuTbx20 | S21906.g09793.01 | | aniseed |
| PfuTbx6.1 | S4584.g06449.01 | | aniseed |
| PfuTbx6.2 | S4556.g06436.01 | | aniseed |
| PmaEomes/Tbr1 | S631.g10737.02 | | aniseed |
| PmaTbx1/10 | S356.g07736.01 | | aniseed |
| PmaTbx15/18/22 | S497.g09447.01 | | aniseed |
| PmaTbx19/Brachyury | S305.g07005.01 | | aniseed |
| PmaTbx2/3 | S17.g00705.01 | | aniseed |
| PmaTbx20 | S175.g04755.01 | | aniseed |
| PmaTbx6.1 | S1645.g15326.01 | | aniseed |
| PmaTbx6.2 | S855.g12396.01 | | aniseed |

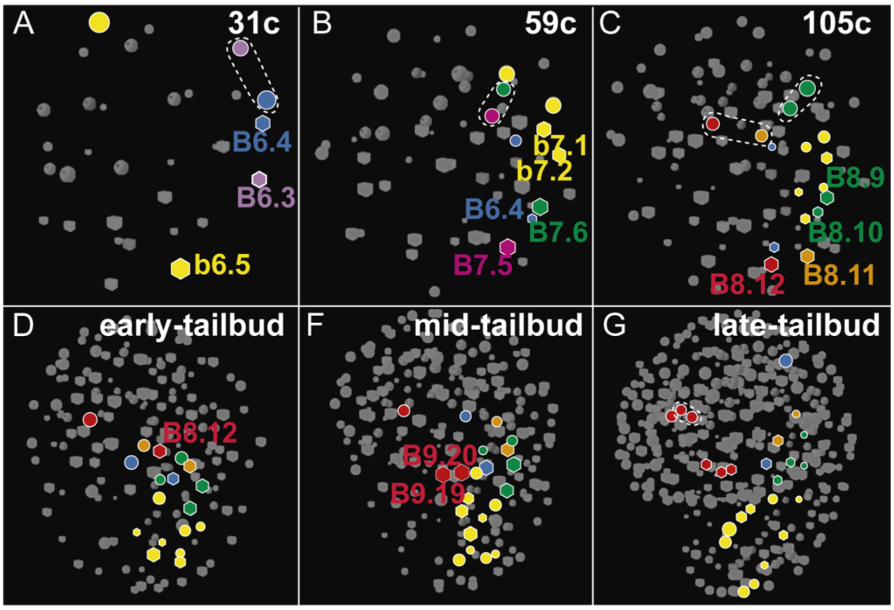


**Sup. Fig. 1**. **4D-reconstruction of a virtual cell tracing experiment based on nuclear position from the 30 cell stage to tailbud stages of *O. dioica* embryos** (Modified from Stach 2008). The fate of the blastomeres are indicated in different colors: muscle and heart (purple, pink), posterior tail muscle cells (yellow), anterior tail muscle cells (orange, green), heart (red), germ-line (blue). Circles and hexagons represent blastomeres derived from right and left sides of the embryo, respectively. Dashed lines indicate twin blastomeres.

Sup. Fig. 2A

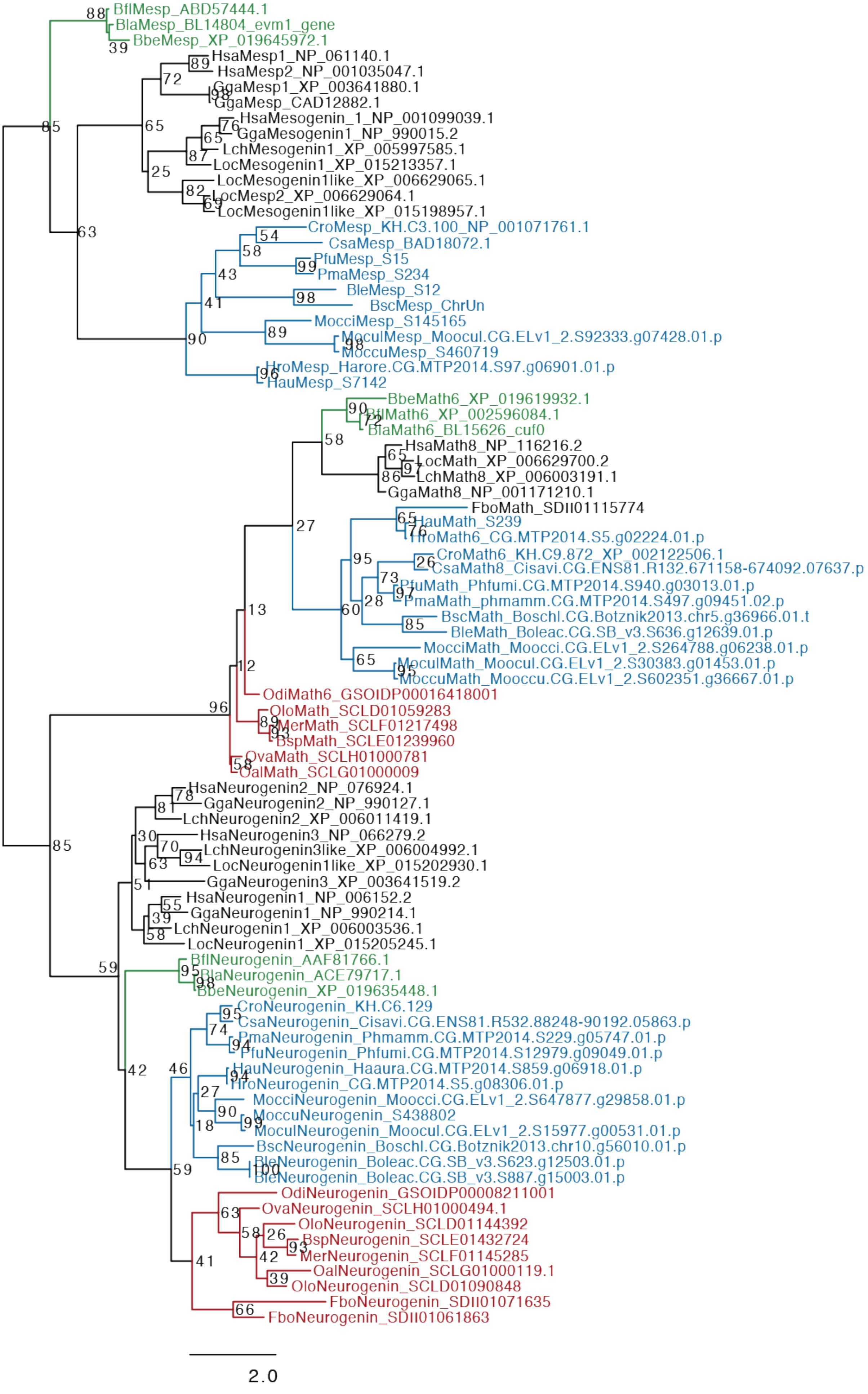
BflMesp_ABD57444.1
BlaMesp_BL14804_evm1_gene
BbeMesp_XP_019645972.1
HsaMesp1_NP_061140.1
HsaMesp2_NP_001035047.1
GgaMesp1_XP_003641880.1
GgaMesp_CAD12882.1
HsaMesogenin_1_NP_001099039.1
GgaMesogenin1_NP_990015.2
LchMesogenin1_XP_005997585.1
LocMesogenin1_XP_015213357.1
LocMesogenin1like_XP_006629065.1
LocMesp2_XP_006629064.1
LocMesogenin1like_XP_015198957.1
CroMesp_KH.C3.100_NP_001071761.1
CsaMesp_BAD18072.1
PfuMesp_S15
PmaMesp_S234
BleMesp_S12
BscMesp_ChrUn
MocciMesp_S145165
MoculMesp_Moocul.CG.ELv1_2.S92333.g07428.01.p
MoccuMesp_S460719
HroMesp_Harore.CG.MTP2014.S97.g06901.01.p
HauMesp_S7142
BbeMath6_XP_019619932.1
BflMath6_XP_002596084.1
BlaMath6_BL15626_cuf0
HsaMath8_NP_116216.2
LocMath_XP_006629700.2
LchMath8_XP_006003191.1
GgaMath8_NP_001171210.1
FboMath_SDII01115774
HauMath_S239
HroMath6_CG.MTP2014.S5.g02224.01.p
CroMath6_KH.C9.872_XP_002122506.1
CsaMath8_Cisavi.CG.ENS81.R132.671158-674092.07637.p
PfuMath_Phfumi.CG.MTP2014.S940.g03013.01.p
PmaMath_phmamm.CG.MTP2014.S497.g09451.02.p
BscMath_Boschl.CG.Botznik2013.chr5.g36966.01.t
BleMath_Boleac.CG.SB_v3.S636.g12639.01.p
MocciMath_Moocci.CG.ELv1_2.S264788.g06238.01.p
MoculMath_Moocul.CG.ELv1_2.S30383.g01453.01.p
MoccuMath_Mooccu.CG.ELv1_2.S602351.g36667.01.p
OdiMath6_GSOIDP00016418001
OloMath_SCLD01059283
MerMath_SCLF01217498
BspMath_SCLE01239960
OvaMath_SCLH01000781
OalMath_SCLG01000009
HsaNeurogenin2_NP_076924.1
GgaNeurogenin2_NP_990127.1
LchNeurogenin2_XP_006011419.1
HsaNeurogenin3_NP_066279.2
LchNeurogenin3like_XP_006004992.1
LocNeurogenin1like_XP_015202930.1
GgaNeurogenin3_XP_003641519.2
HsaNeurogenin1_NP_006152.2
GgaNeurogenin1_NP_990214.1
LchNeurogenin1_XP_006003536.1
LocNeurogenin1_XP_015205245.1
BflNeurogenin_AAF81766.1
BlaNeurogenin_ACE79717.1
BbeNeurogenin_XP_019635448.1
CroNeurogenin_KH.C6.129
CsaNeurogenin_Cisavi.CG.ENS81.R532.88248-90192.05863.p
PmaNeurogenin_Phmamm.CG.MTP2014.S229.g05747.01.p
PfuNeurogenin_Phfumi.CG.MTP2014.S12979.g09049.01.p
HauNeurogenin_Haaura.CG.MTP2014.S859.g06918.01.p
HroNeurogenin_CG.MTP2014.S5.g08306.01.p
MocciNeurogenin_Moocci.CG.ELv1_2.S647877.g29858.01.p
MoccuNeurogenin_S438802
MoculNeurogenin_Moocul.CG.ELv1_2.S15977.g00531.01.p
BscNeurogenin_Boschl.CG.Botznik2013.chr10.g56010.01.p
BleNeurogenin_Boleac.CG.SB_v3.S623.g12503.01.p
BleNeurogenin_Boleac.CG.SB_v3.S887.g15003.01.p
OdiNeurogenin_GSOIDP00008211001
OvaNeurogenin_SCLH01000494.1
OloNeurogenin_SCLD01144392
BspNeurogenin_SCLE01432724
MerNeurogenin_SCLF01145285
OalNeurogenin_SCLG01000119.1
OloNeurogenin_SCLD01090848
FboNeurogenin_SDII01071635
FboNeurogenin_SDII01061863
2.0

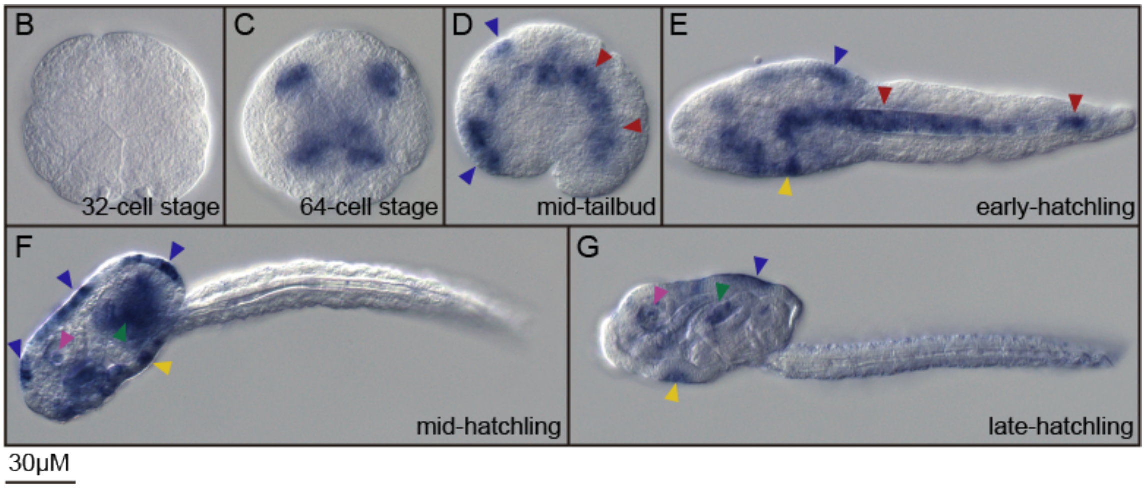


**Sup. Fig. 2**. (**A**) Unrooted phylogenetic tree, represented in a rectangular layout for the sake clarity, showing the presence of bHLH homologs of *Neurogenin* and *Math* in appendicularians, but the absence of *Mesp*, suggesting an ancestral loss of this gene family in this clade. Bootstrap values are shown in the nodes. Vertebrates: *Gallus gallus* (Gga), *Homo sapiens* (Hsa), *Latimeria chalumnae* (Lch), *Lepisosteus oculatus* (Loc); Tunicates: *Bathochordaeus sp*. (Bsp). *Botrylloides leachii* (Ble), *Botrylloides schlosseri* (Bsc), *Ciona robusta* (Cro), *Ciona savignyi* (Csa), *Fritillaria borealis* (Fbo), *Halocynthia aurantium* (Hau), *Halocynthia roretzi* (Hro), *Mesochordaeus erythrocephalus* (Mer), *Molgula occidentalis* (Mocci), *Molgula occulta* (Moccu), *Molgula oculata* (Mocul), *Oikopleura albicans* (Oal), *Oikopleura dioica* (Odi), *Oikopleura longicauda* (Olo), *Oikopleura vanhoeffeni* (Ova), *Phallusia fumigata* (Pfu), *Phallusia mammillata* (Pma); Cephalochordates: *Branchiostoma belcheri* (Bbe), *Branchiostoma floridae* (Bfl), *Branchiostoma lanceolatum* (Bla). (**B-G**) Developmental expression pattern of *O. dioica Math6* homolog. Whole mount in situ hybridization in different stages of *O. dioica* development showing expression in the notochord in tailbud and early-hatchling embryos (red arrowheads) (**C, D**), in epidermis (blue arrowheads) (**C-F**), in the annal domain in hatchling stages (yellow arrowheads) (**D-F**), in later stages of neural system development (pink arrowheads) (**E-F**), and in later stages of digestive system development (green arrowheads) (**E-F**). Images from tailbud in advance correspond to left lateral views orientated anterior towards the left and dorsal towards the top.

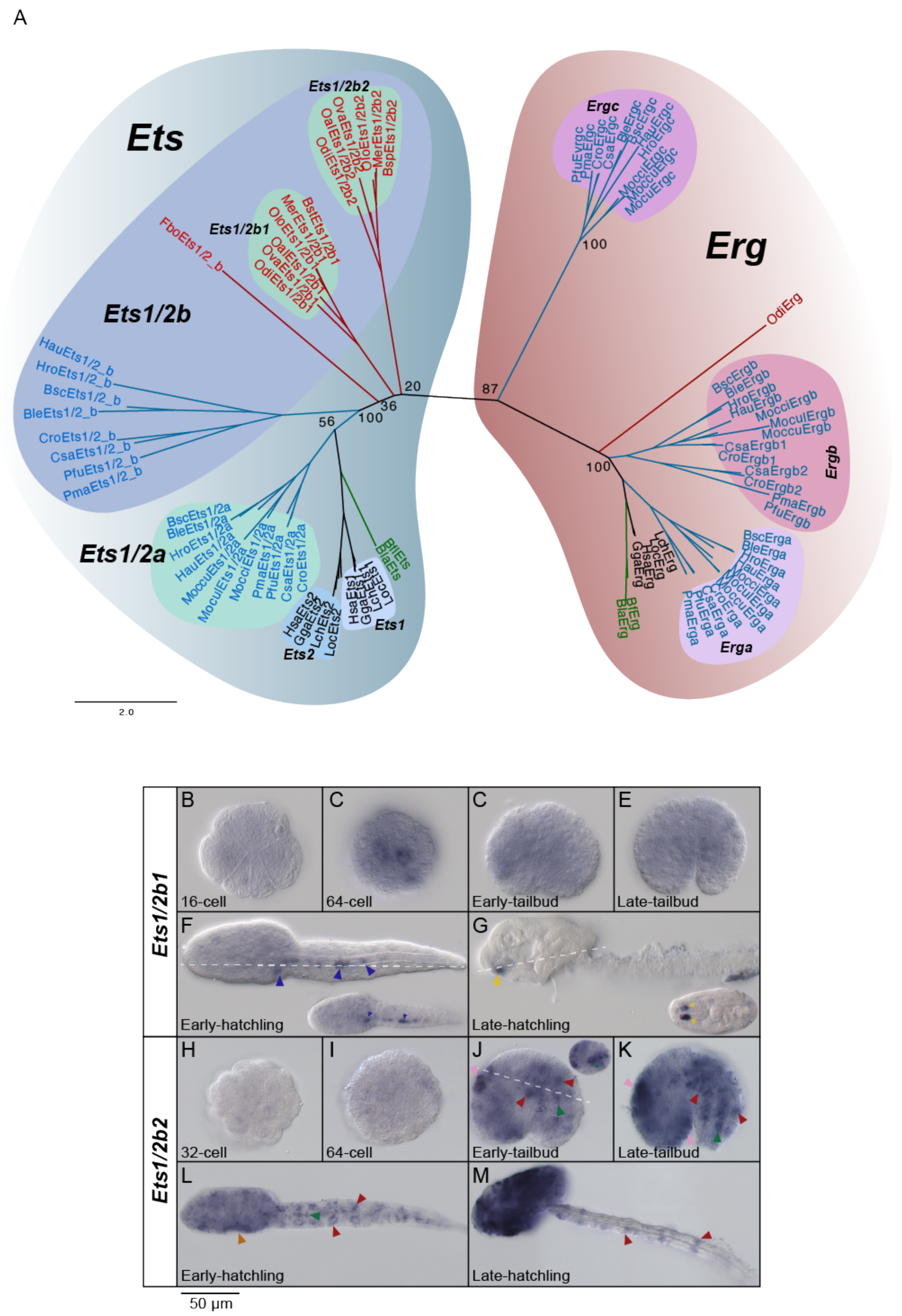
A
Ets
Erg
Ets1/2b
Ets1/2a
Ets1/2b1
Ets1/2b2
FboEts1/2_b
HauEts1/2_b
HroEts1/2_b
BscEts1/2_b
BleEts1/2_b
CroEts1/2_b
CsaEts1/2_b
PfuEts1/2_b
PmaEts1/2_b
Ets1
Ets2
Ergc
Ergb
Erga
OdiErg
20
36
56
100
87
100
100
2.0
B
C
C
E
16-cell
64-cell
Early-tailbud
Late-tailbud
F
G
Early-hatchling
Late-hatchling
H
I
J
K
32-cell
64-cell
Early-tailbud
Late-tailbud
L
M
Early-hatchling
Late-hatchling
Ets1/2b1
Ets1/2b2
50 µm

**Sup. Fig. 3. Phylogenetic analysis of the *Ets* family and developmental expression patterns of *O. dioica Ets1/2b* paralogs.** (**A**) Phylogenetic analysis of the *Ets* and *Erg* protein families showed a high bootstrap value separating both protein families what corroborated the existence of two Ets1/2 proteins in appendicularians. Vertebrates: *Gallus gallus* (Gga), *Homo sapiens* (Hsa), *Latimeria chalumnae* (Lch), *Lepisosteus oculatus* (Loc); Tunicates: *Bathochordaeus sp*. (Bsp). *Botrylloides leachii* (Ble), *Botrylloides schlosseri* (Bsc), *Ciona robusta* (Cro), *Ciona savignyi* (Csa), *Fritillaria borealis* (Fbo), *Halocynthia aurantium* (Hau), *Halocynthia roretzi* (Hro), *Mesochordaeus erythrocephalus* (Mer), *Molgula occidentalis* (Mocci), *Molgula occulta* (Moccu), *Molgula oculata* (Mocul), *Oikopleura albicans* (Oal), *Oikopleura dioica* (Odi), *Oikopleura longicauda* (Olo), *Oikopleura vanhoeffeni* (Ova), *Phallusia fumigata* (Pfu), *Phallusia mammillata* (Pma); Cephalochordates: *Branchiostoma belcheri* (Bbe), *Branchiostoma floridae* (Bfl), *Branchiostoma lanceolatum* (Bla). Whole mount in situ hybridization of *O. dioica Ets1/2b1* did not show any clear expression before hatchling stages (**B-E**). In early-hatchling stage *Ets1/2b1* revealed expression in the migratory endodermal strand cells (blue arrowheads) (**F**). In late-hatchling the expression signal was restricted to the buccal gland (yellow arrowheads) (**G**). *Ets1/2b2* did not show expression until tailbud stage (**H, I**). In tailbud embryos, expression signal was detected in tail muscle cells (red arrowheads), the notochord (green arrowheads) and the epidermis of the trunk (pink arrowheads) (**J, K**). In early-hatchling expression signal continued in the tail muscle and the notochord and increased in the anal domain (orange arrowheads) (**L**). In late hatchling stage, the *Ets1/2b2* expression covered the entire oikoplastic epithelium, and continued in the muscle cells of the tail (**M**). Large images from tailbud in advance correspond to left lateral views oriented anterior towards the left and dorsal towards the top. Inset images are dorsal views of optical cross sections at the levels of dashed lines.

Sup. Fig. 4A

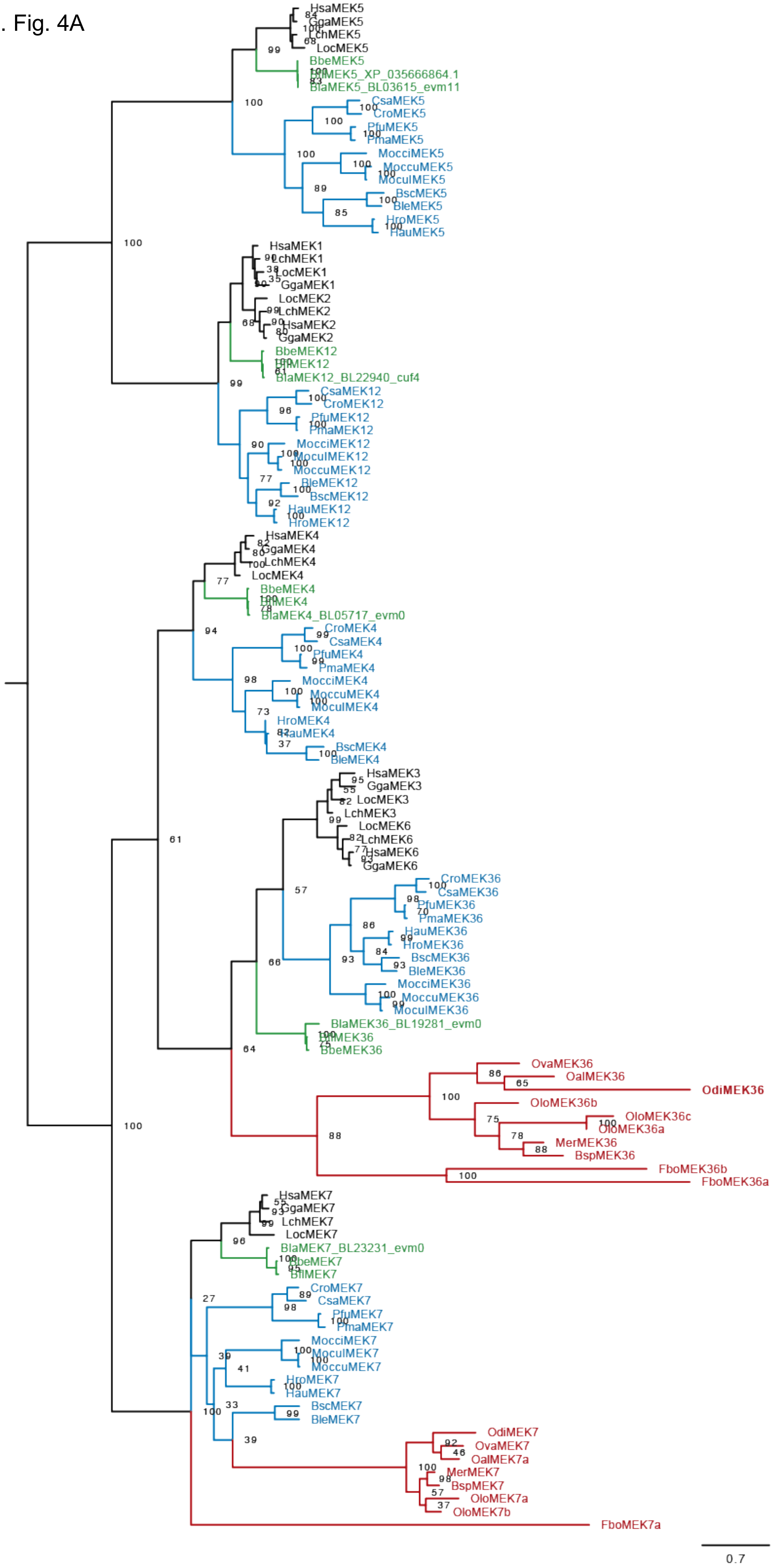
HsaMEK5
GgaMEK5
LchMEK5
LocMEK5
BbeMEK5
BfiMEK5_XP_035666864.1
BlaMEK5_BL03615_evm11
CsaMEK5
CroMEK5
PfuMEK5
PmaMEK5
MocciMEK5
MoccuMEK5
MoculMEK5
BscMEK5
BleMEK5
HroMEK5
HauMEK5
HsaMEK1
LchMEK1
LocMEK1
GgaMEK1
LocMEK2
LchMEK2
HsaMEK2
GgaMEK2
BbeMEK12
BfiMEK12
BlaMEK12_BL22940_cuf4
CsaMEK12
CroMEK12
PfuMEK12
PmaMEK12
MocciMEK12
MoculMEK12
MoccuMEK12
BleMEK12
BscMEK12
HauMEK12
HroMEK12
HsaMEK4
GgaMEK4
LchMEK4
LocMEK4
BbeMEK4
BfiMEK4
BlaMEK4_BL05717_evm0
CroMEK4
CsaMEK4
PfuMEK4
PmaMEK4
MocciMEK4
MoccuMEK4
MoculMEK4
HroMEK4
HauMEK4
BscMEK4
BleMEK4
HsaMEK3
GgaMEK3
LocMEK3
LchMEK3
LocMEK6
LchMEK6
HsaMEK6
GgaMEK6
CroMEK36
CsaMEK36
PfuMEK36
PmaMEK36
HauMEK36
HroMEK36
BscMEK36
BleMEK36
MocciMEK36
MoccuMEK36
MoculMEK36
BlaMEK36_BL19281_evm0
BfiMEK36
BbeMEK36
OvaMEK36
OalMEK36
OdiMEK36
OloMEK36b
OloMEK36c
OloMEK36a
MerMEK36
BspMEK36
FboMEK36b
FboMEK36a
HsaMEK7
GgaMEK7
LchMEK7
LocMEK7
BlaMEK7_BL23231_evm0
BbeMEK7
BfiMEK7
CroMEK7
CsaMEK7
PfuMEK7
PmaMEK7
MocciMEK7
MoculMEK7
MoccuMEK7
HroMEK7
HauMEK7
BscMEK7
BleMEK7
OdiMEK7
OvaMEK7
OalMEK7a
MerMEK7
BspMEK7
OloMEK7a
OloMEK7b
FboMEK7a
0.7

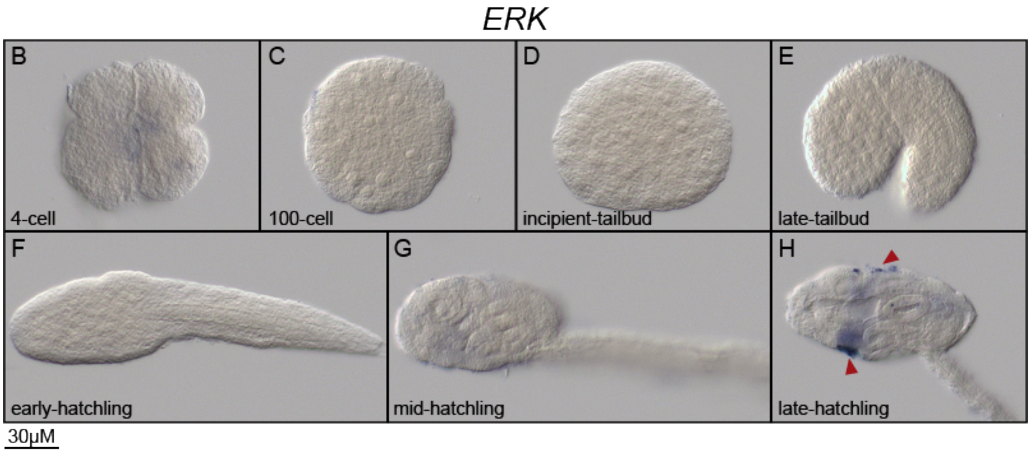


**Sup. Fig. 4.** (**A**) ML phylogenetic tree of the *MEK* subfamilies in chordates revealing the loss of the *MEK4*, *MEK5* and *MEK1/2* subfamilies in appendicularians, but the surviving of *MEK7* and *MEK3/6* subfamilies. Scale bar indicates amino acid substitutions. Bootstrap values are shown in the nodes. Vertebrates: *Gallus gallus* (Gga), *Homo sapiens* (Hsa), *Latimeria chalumnae* (Lch), *Lepisosteus oculatus* (Loc); Tunicates: *Bathochordaeus sp*. (Bsp). *Botrylloides leachii* (Ble), *Botrylloides schlosseri* (Bsc), *Ciona robusta* (Cro), *Ciona savignyi* (Csa), *Fritillaria borealis* (Fbo), *Halocynthia aurantium* (Hau), *Halocynthia roretzi* (Hro), *Mesochordaeus erythrocephalus* (Mer), *Molgula occidentalis* (Mocci), *Molgula occulta* (Moccu), *Molgula oculata* (Mocul), *Oikopleura albicans* (Oal), *Oikopleura dioica* (Odi), *Oikopleura longicauda* (Olo), *Oikopleura vanhoeffeni* (Ova), *Phallusia fumigata* (Pfu), *Phallusia mammillata* (Pma); Cephalochordates: *Branchiostoma belcheri* (Bbe), *Branchiostoma floridae* (Bfl), *Branchiostoma lanceolatum* (Bla). (**B-H**) Developmental expression pattern of O. dioica *ERK* homolog. Whole mount in situ hybridization in different stages of *O. dioica* did not detect expression of *ERK* in any studied stage (**B-G**) until late-hatchling stage when expression is detected in the oikoplastic epithelium (**H**). Images from tailbud in advanced correspond to left lateral views orientated anterior towards the left and dorsal towards the top. Red arrowheads indicate the expression in the oikoplastic epithelium.

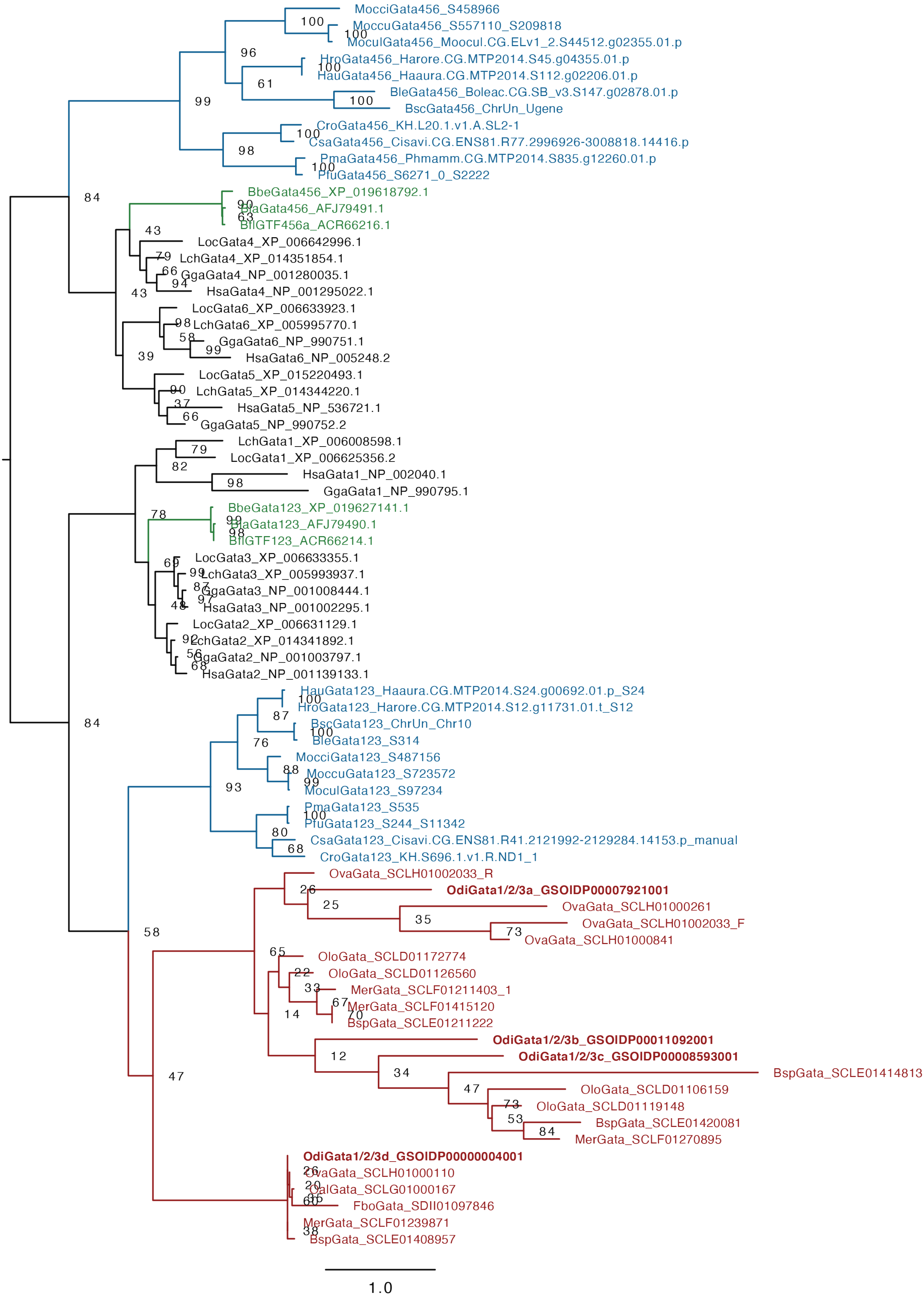
MocciGata456_S458966
MoccuGata456_S557110_S209818
MoculGata456_Moocul.CG.ELv1_2.S44512.g02355.01.p
HroGata456_Harore.CG.MTP2014.S45.g04355.01.p
HauGata456_Haaura.CG.MTP2014.S112.g02206.01.p
BleGata456_Boleac.CG.SB_v3.S147.g02878.01.p
BscGata456_ChrUn_Ugene
CroGata456_KH.L20.1.v1.A.SL2-1
CsaGata456_Cisavi.CG.ENS81.R77.2996926-3008818.14416.p
PmaGata456_Phmamm.CG.MTP2014.S835.g12260.01.p
PfuGata456_S6271_0_S2222
BbeGata456_XP_019618792.1
BlaGata456_AFJ79491.1
BflGTF456a_ACR66216.1
LocGata4_XP_006642996.1
LchGata4_XP_014351854.1
GgaGata4_NP_001280035.1
HsaGata4_NP_001295022.1
LocGata6_XP_006633923.1
LchGata6_XP_005995770.1
GgaGata6_NP_990751.1
HsaGata6_NP_005248.2
LocGata5_XP_015220493.1
LchGata5_XP_014344220.1
HsaGata5_NP_536721.1
GgaGata5_NP_990752.2
LchGata1_XP_006008598.1
LocGata1_XP_006625356.2
HsaGata1_NP_002040.1
GgaGata1_NP_990795.1
BbeGata123_XP_019627141.1
BlaGata123_AFJ79490.1
BflGTF123_ACR66214.1
LocGata3_XP_006633355.1
LchGata3_XP_005993937.1
GgaGata3_NP_001008444.1
HsaGata3_NP_001002295.1
LocGata2_XP_006631129.1
LchGata2_XP_014341892.1
GgaGata2_NP_001003797.1
HsaGata2_NP_001139133.1
HauGata123_Haaura.CG.MTP2014.S24.g00692.01.p_S24
HroGata123_Harore.CG.MTP2014.S12.g11731.01.t_S12
BscGata123_ChrUn_Chr10
BleGata123_S314
MocciGata123_S487156
MoccuGata123_S723572
MoculGata123_S97234
PmaGata123_S535
PfuGata123_S244_S11342
CsaGata123_Cisavi.CG.ENS81.R41.2121992-2129284.14153.p_manual
CroGata123_KH.S696.1.v1.R.ND1_1
OvaGata_SCLH01002033_R
OdiGata1/2/3a_GSOIDP00007921001
OvaGata_SCLH01000261
OvaGata_SCLH01002033_F
OvaGata_SCLH01000841
OloGata_SCLD01172774
OloGata_SCLD01126560
MerGata_SCLF01211403_1
MerGata_SCLF01415120
BspGata_SCLE01211222
OdiGata1/2/3b_GSOIDP00011092001
OdiGata1/2/3c_GSOIDP00008593001
BspGata_SCLE01414813
OloGata_SCLD01106159
OloGata_SCLD01119148
BspGata_SCLE01420081
MerGata_SCLF01270895
OdiGata1/2/3d_GSOIDP00000004001
OvaGata_SCLH01000110
OalGata_SCLG01000167
FboGata_SDII01097846
MerGata_SCLF01239871
BspGata_SCLE01408957
1.0

**Sup. Fig. 5.** ML phylogenetic tree of the Gata subfamilies in chordates revealing the loss of the *Gata4/5/6* in appendicularians, but the surviving and lineage specific duplications of *Gata1/2/3* in appendicularians. Scale bar indicates amino acid substitutions. Bootstrap values are shown in the nodes. Vertebrates: *Gallus gallus* (Gga), *Homo sapiens* (Hsa), *Latimeria chalumnae* (Lch), *Lepisosteus oculatus* (Loc); Tunicates: *Bathochordaeus sp*. (Bsp). *Botrylloides leachii* (Ble), *Botrylloides schlosseri* (Bsc), *Ciona robusta* (Cro), *Ciona savignyi* (Csa), *Fritillaria borealis* (Fbo), *Halocynthia aurantium* (Hau), *Halocynthia roretzi* (Hro), *Mesochordaeus erythrocephalus* (Mer), *Molgula occidentalis* (Mocci), *Molgula occulta* (Moccu), *Molgula oculata* (Mocul), *Oikopleura albicans* (Oal), *Oikopleura dioica* (Odi), *Oikopleura longicauda* (Olo), *Oikopleura vanhoeffeni* (Ova), *Phallusia fumigata* (Pfu), *Phallusia mammillata* (Pma); Cephalochordates: *Branchiostoma belcheri* (Bbe), *Branchiostoma floridae* (Bfl), *Branchiostoma lanceolatum* (Bla).

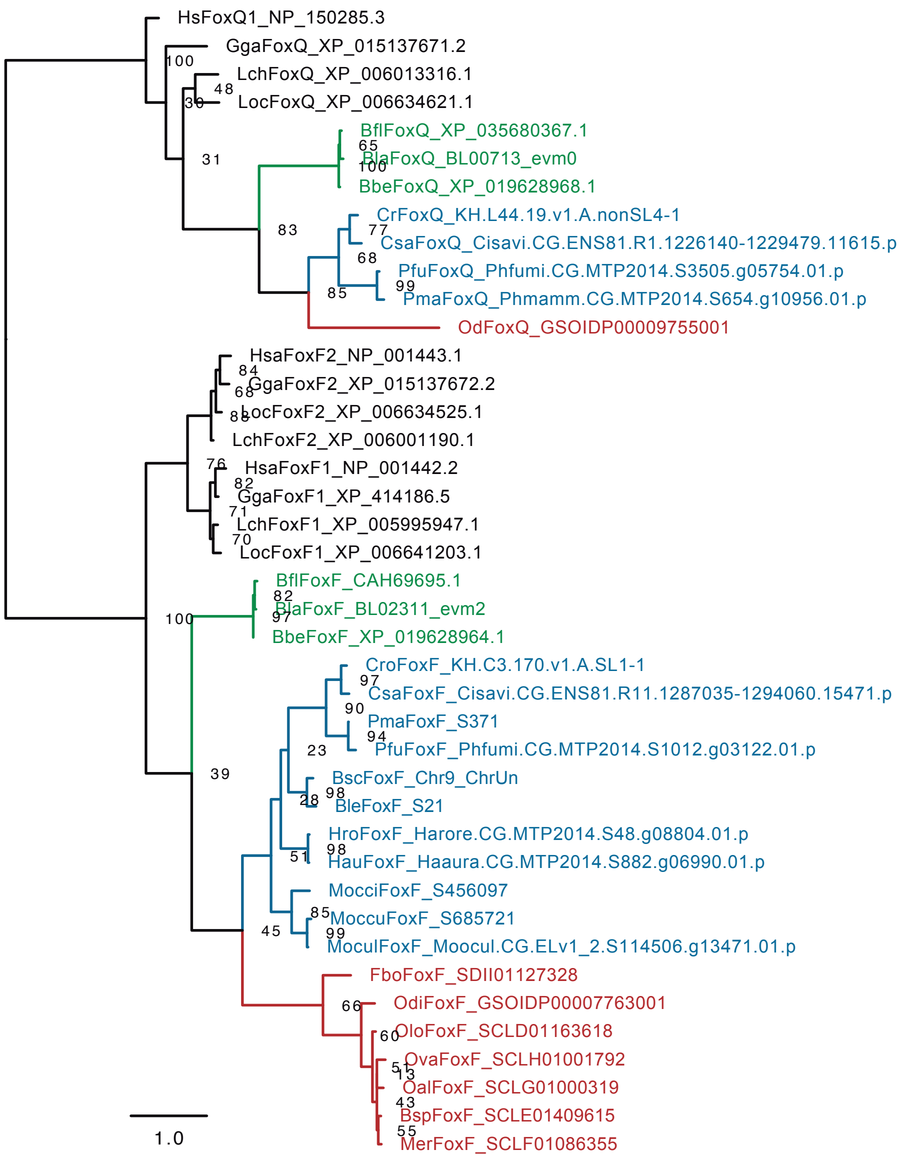


**Sup. Fig. 6.** ML phylogenetic tree of the FoxF subfamily in chordates reveals the presence of an ortholog of *FoxF* in appendicularians. The sister *FoxQ* subfamily was used as outgroup to root the tree. Scale bar indicates amino acid substitutions. Bootstrap values are shown in the nodes. Vertebrates: *Gallus gallus* (Gga), *Homo sapiens* (Hsa), *Latimeria chalumnae* (Lch), *Lepisosteus oculatus* (Loc); Tunicates: *Bathochordaeus sp*. (Bsp). *Botrylloides leachii* (Ble), *Botrylloides schlosseri* (Bsc), *Ciona robusta* (Cro), *Ciona savignyi* (Csa), *Fritillaria borealis* (Fbo), *Halocynthia aurantium* (Hau), *Halocynthia roretzi* (Hro), *Mesochordaeus erythrocephalus* (Mer), *Molgula occidentalis* (Mocci), *Molgula occulta* (Moccu), *Molgula oculata* (Mocul), *Oikopleura albicans* (Oal), *Oikopleura dioica* (Odi), *Oikopleura longicauda* (Olo), *Oikopleura vanhoeffeni* (Ova), *Phallusia fumigata* (Pfu), *Phallusia mammillata* (Pma); Cephalochordates: *Branchiostoma belcheri* (Bbe), *Branchiostoma floridae* (Bfl), *Branchiostoma lanceolatum* (Bla).

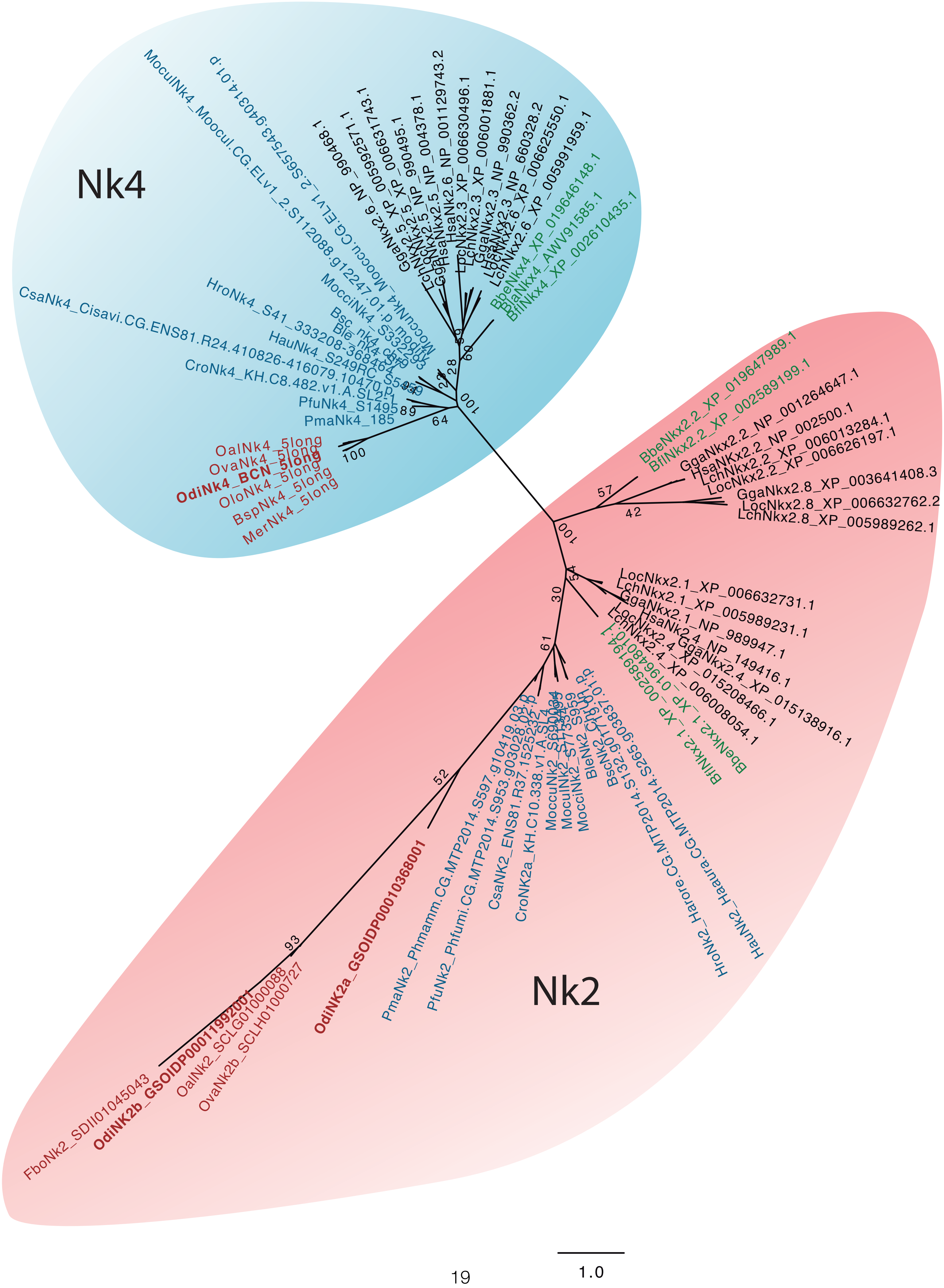

Nk4
Nk2
OdiNk4_BCN_5long
OalNk4_5long
OvaNk4_5long
OloNk4_5long
BspNk4_5long
MerNk4_5long
PmaNk4_185
PfuNk4_S1495
CsaNk4_Cisavi.CG.ENS81.R24.410826-416079.10470.p1
HroNk4_S41_333208-368464
HauNk4_S249RC_S5459
BbeNkx4_XP_019646148.1
BlaNkx4_AWV91585.1
BflNkx4_XP_002610435.1
BbeNkx2.2_XP_019647989.1
BflNkx2.2_XP_002589199.1
GgaNkx2.2_NP_001264647.1
HsaNkx2.2_NP_002500.1
LchNkx2.2_XP_006013284.1
LocNkx2.2_XP_006626197.1
GgaNkx2.8_XP_003641408.3
LocNkx2.8_XP_006632762.2
LchNkx2.8_XP_005989262.1
LocNkx2.1_XP_006632731.1
LchNkx2.1_XP_005989231.1
GgaNkx2.1_NP_989947.1
HsaNkx2.4_NP_149416.1
GgaNkx2.4_XP_015138916.1
LocNkx2.4_XP_015208466.1
LchNkx2.4_XP_006008054.1
BbeNkx2.1_XP_019648010.1
BflNkx2.1_XP_002589194.1
OdiNK2a_GSOIDP00010368001
OdiNK2b_GSOIDP00011992001
FboNk2_SDII01045043
OalNk2_SCLG01000088
OvaNk2b_SCLH01000727
PmaNk2_Phmamm.CG.MTP2014.S597.g10419.03.p
PfuNk2_Phfumi.CG.MTP2014.S953.g03028.03.p
CsaNK2_ENS81.R37.1525235-1
CroNK2a_KH.C10.338.v1.A.SL1
HroNk2_Harore.CG.MTP2014.S265.g03837.01.p
HauNk2_Haaura.CG.MTP2014
100
64
89
60
50
28
57
42
30
61
52
93
1.0

**Sup. Fig. 7.** ML phylogenetic tree of the Nk subfamilies in chordates reveals the presence of an ortholog of *Nk4* in appendicularians and two orthologs of the Nk2 subfamily. Scale bar indicates amino acid substitutions. Bootstrap values are shown in the nodes. Vertebrates: *Gallus gallus* (Gga), *Homo sapiens* (Hsa), *Latimeria chalumnae* (Lch), *Lepisosteus oculatus* (Loc); Tunicates: *Bathochordaeus sp*. (Bsp). *Botrylloides leachii* (Ble), *Botrylloides schlosseri* (Bsc), *Ciona robusta* (Cro), *Ciona savignyi* (Csa), *Fritillaria borealis* (Fbo), *Halocynthia aurantium* (Hau), *Halocynthia roretzi* (Hro), *Mesochordaeus erythrocephalus* (Mer), *Molgula occidentalis* (Mocci), *Molgula occulta* (Moccu), *Molgula oculata* (Mocul), *Oikopleura albicans* (Oal), *Oikopleura dioica* (Odi), *Oikopleura longicauda* (Olo), *Oikopleura vanhoeffeni* (Ova), *Phallusia fumigata* (Pfu), *Phallusia mammillata* (Pma); Cephalochordates: *Branchiostoma belcheri* (Bbe), *Branchiostoma floridae* (Bfl), *Branchiostoma lanceolatum* (Bla).

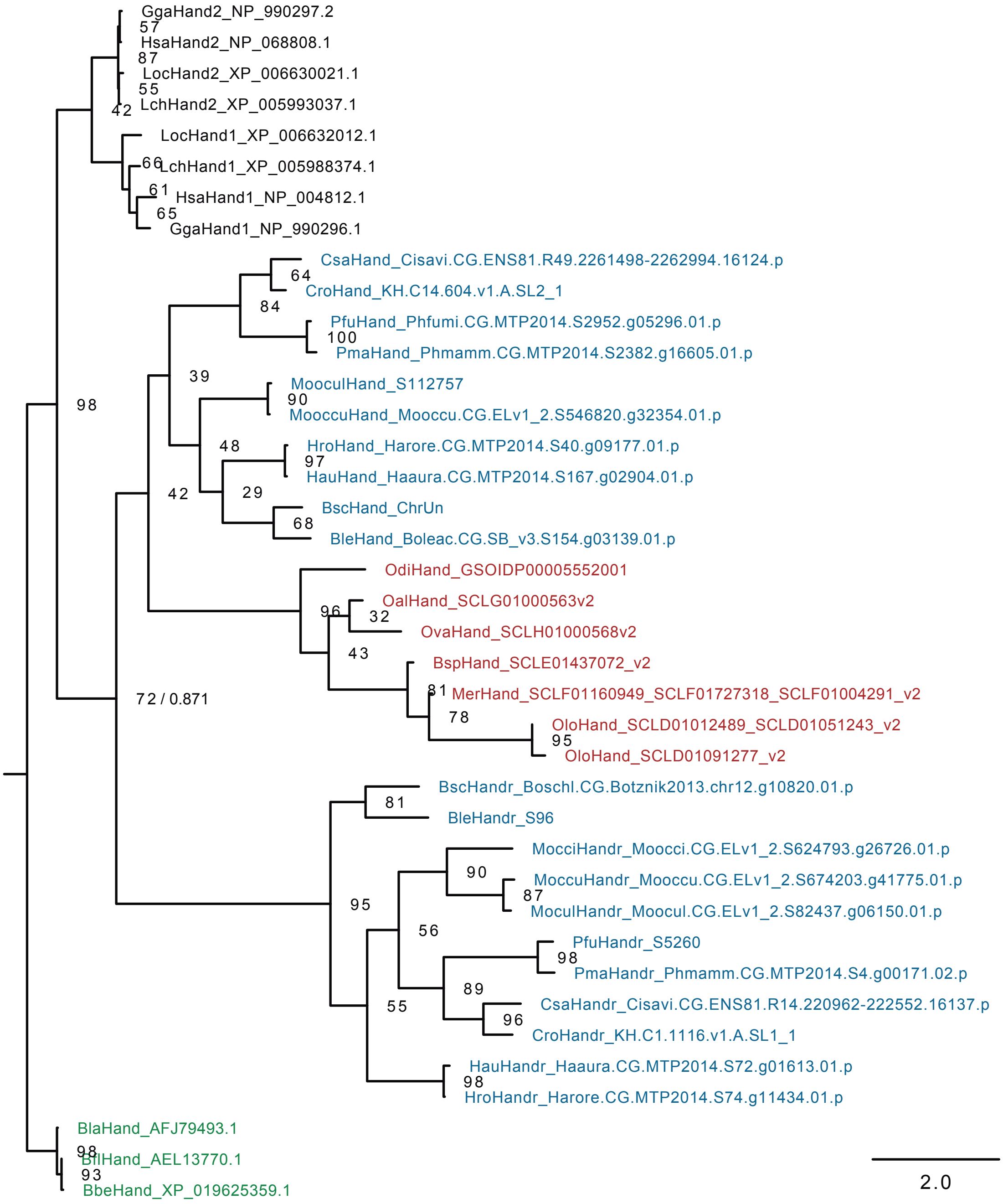


**Sup. Fig. 8.** ML phylogenetic tree of the Hand subfamilies suggest that member of this family in *O. dioica* is homologous to ascidian *Hand1/2*. Despite the tree suggests that the second paralog of ascidian (*Hand-r*) arose by a duplication at the base of the tunicate clade, and therefore subsequently lost in appendicularians. The low bootstrap support and the presence of shared long amino acid domain rich in K between the *Hand1/2* and *Hand-r* in ascidians, but absent in appendicularians, do not allow us to discard the possibility that *Hand-r* was originated by a duplication within the ascidian lineage, and its basal braching in the tunicate clade is due to a long branch attraction phenomenon. Scale bar indicates amino acid substitutions. Bootstrap values are shown in the nodes. Vertebrates: *Gallus gallus* (Gga), *Homo sapiens* (Hsa), *Latimeria chalumnae* (Lch), *Lepisosteus oculatus* (Loc); Tunicates: *Bathochordaeus sp*. (Bsp). *Botrylloides leachii* (Ble), *Botrylloides schlosseri* (Bsc), *Ciona robusta* (Cro), *Ciona savignyi* (Csa), *Fritillaria borealis* (Fbo), *Halocynthia aurantium* (Hau), *Halocynthia roretzi* (Hro), *Mesochordaeus erythrocephalus* (Mer), *Molgula occidentalis* (Mocci), *Molgula occulta* (Moccu), *Molgula oculata* (Mocul), *Oikopleura albicans* (Oal), *Oikopleura dioica* (Odi), *Oikopleura longicauda* (Olo), *Oikopleura vanhoeffeni* (Ova), *Phallusia fumigata* (Pfu), *Phallusia mammillata* (Pma); Cephalochordates: *Branchiostoma belcheri* (Bbe), *Branchiostoma floridae* (Bfl), *Branchiostoma lanceolatum* (Bla).

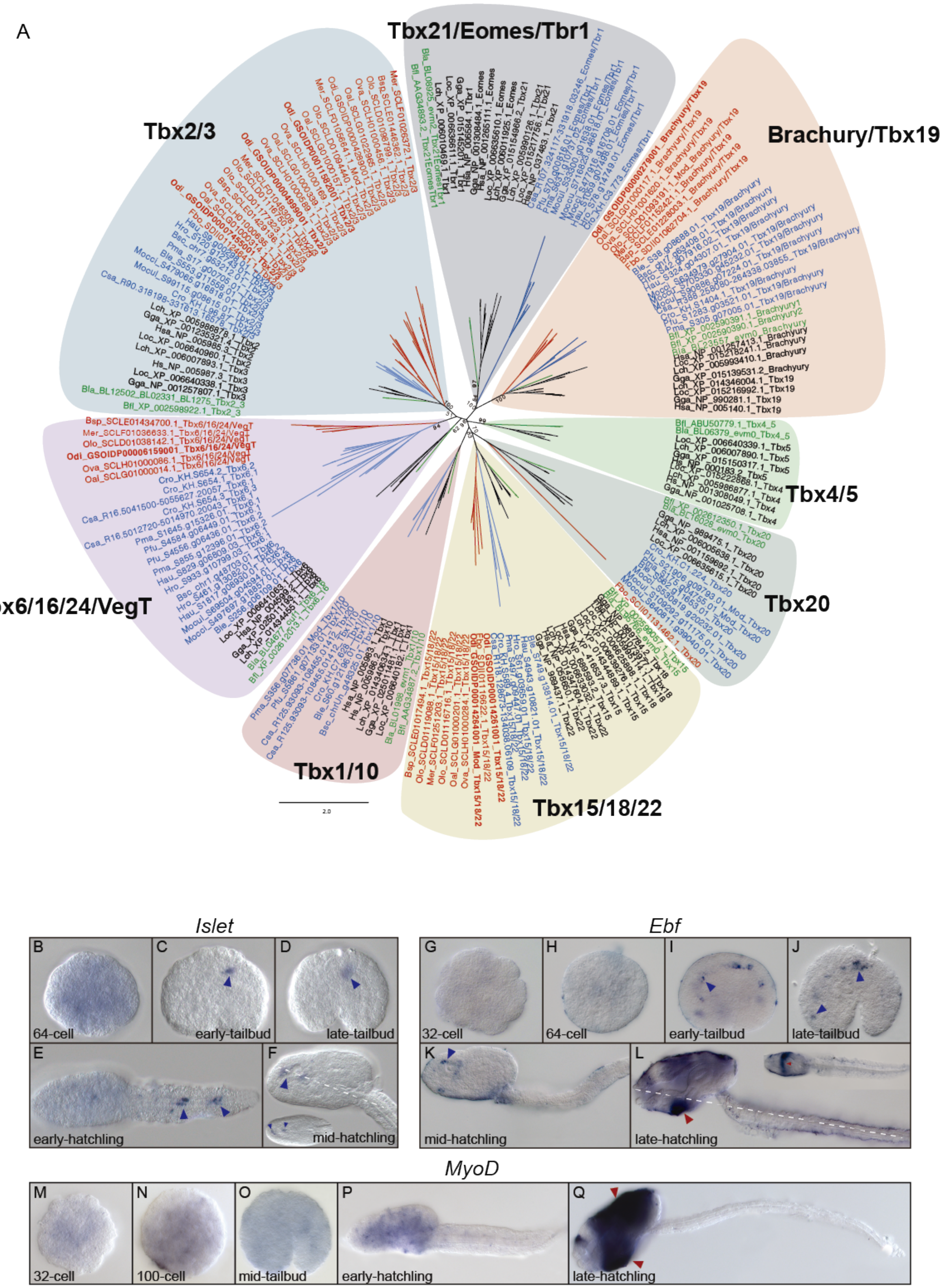
A
Tbx21/Eomes/Tbr1
Tbx2/3
Brachury/Tbx19
Tbx6/16/24/VegT
Tbx4/5
Tbx20
Tbx1/10
Tbx15/18/22
2.0
Islet
B
64-cell
C
early-tailbud
D
late-tailbud
E
early-hatchling
F
mid-hatchling
Ebf
G
32-cell
H
64-cell
I
early-tailbud
J
late-tailbud
K
mid-hatchling
L
late-hatchling
MyoD
M
32-cell
N
100-cell
O
mid-tailbud
P
early-hatchling
Q
late-hatchling

**Sup. Fig. 9. Phylogenetic analysis of the *Tbx* family and developmental expression patterns of *O. dioica Islet*, *Ebf,* and MyoD homologs**. (**A**) ML phylogenetic tree of the Tbx subfamilies in chordates reveals the loss of Tbx1/10 and Tbx21/Eomes/Tbr1 subfamilies in appendicularians and the ancestral loss of Tbx4/5 subfamily in tunicates. Scale bar indicates amino acid substitutions. Bootstrap values are shown in the nodes. Vertebrates: *Gallus gallus* (Gga), *Homo sapiens* (Hsa), *Latimeria chalumnae* (Lch), *Lepisosteus oculatus* (Loc); Tunicates: *Bathochordaeus sp.* (Bsp). *Botrylloides leachii* (Ble), *Botrylloides schlosseri* (Bsc), *Ciona robusta* (Cro), *Ciona savignyi* (Csa), *Fritillaria borealis* (Fbo), *Halocynthia aurantium* (Hau), *Halocynthia roretzi* (Hro), *Mesochordaeus erythrocephalus* (Mer), *Molgula occidentalis* (Mocci), *Molgula occulta* (Moccu), *Molgula oculata* (Mocul), *Oikopleura albicans* (Oal), *Oikopleura dioica* (Odi), *Oikopleura longicauda* (Olo), *Oikopleura vanhoeffeni* (Ova), *Phallusia fumigata* (Pfu), *Phallusia mammillata* (Pma); Cephalochordates: *Branchiostoma floridae* (Bfl), *Branchiostoma lanceolatum* (Bla). (**B-Q**) Whole mount in situ hybridization of *O. dioica Islet, Ebf* and *MyoD* homologs. 64-cell embryos did not showed expression of *Islet* (**B**) which was only detected in the developing nervous system from tailbud to hatchling embryos (**C-F**). *Ebf* (*COE*) did not showed expression in early stages (**G-H**) but we detected expression in the nervous system from tailbud to mid-hatchling stage (**I-K**) and in the epidermis of late-hatchling embryos (**L**). We did not detect expression of *MyoD* from 32-cell to hatchling embryos (**M-P**). In late-hatchling embryos *MyoD* was expressed in the oikoplastic epithelium (**Q**). Large images from tailbud in advance correspond to left lateral views oriented anterior towards the left and dorsal towards the top. Inset images are dorsal views of optical cross sections at the levels of dashed lines. Blue arrows indicate the developing nervous system. Red arrows indicate the oikoplastic epithelium.

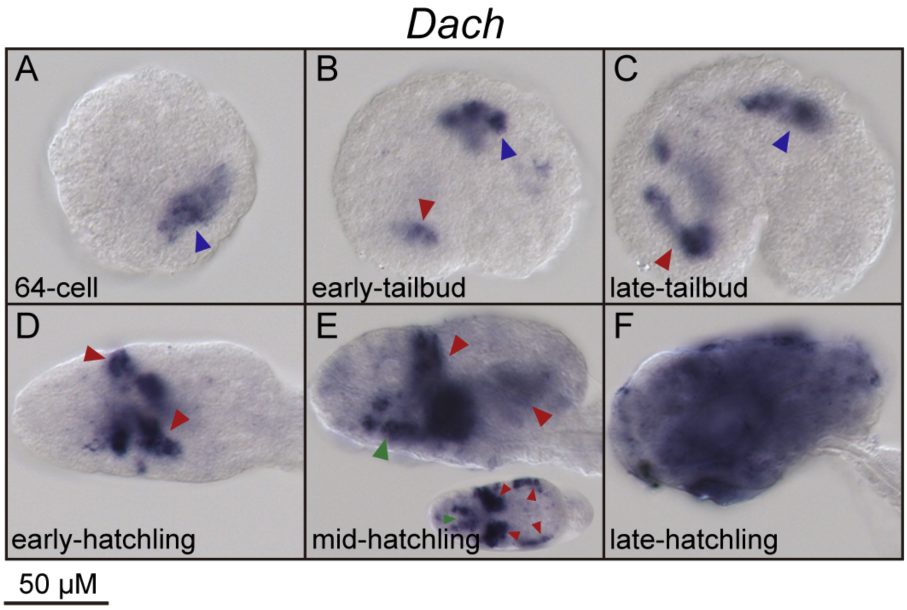


**Sup. Fig. 10. Developmental expression pattern of *O. dioica Dach* homolog**. Whole mount in situ hybridization of *O. dioica Dach* homolog show expression in the developing nervous system in 64-cell and tailbud stages (**A-C**). In tailbud stages, *Dach* start expressing in the trunk epidermis which is maintain until late-hatchling stages when it is expressed in the whole oikoplastic epithelium (**B-F**). In mid-hatchling stage, beside the epidermis, Dach expression is also detected in the endostyle (**E**). Images from tailbud in advance correspond to left lateral views oriented anterior towards the left and dorsal towards the top. Blue arrowheads indicate the nervous system. Red arrowheads indicate the epidermis. Green arrowheads indicate the endostyle.

**Supplementary Table 2**. Information about the cloning fragments and riboprobes and primers used to amplify them.

| Gene | Forward | Reverse | source; length; vector; RNA-pol/digestion enzyme |
|---|---|---|---|
| ***OdActnM1 cross-hybridizing*** | 5'..GTCCCCGCCATGTACGTCTG..3' | 5'..GCATCGGAATCGCTCGTTACCA..3' | gDNA exon 2 partial; 389bp; PCR4-TOPO; T3/NotI |
| ***OdActnM1 specific*** | 5'..GATCGTCCACCGAAAGTGC..3' | 5'..GTCAGCAACTGTTTGAATATATTG..3' | cDNA 3'UTR; 351bp; PCR4-TOPO; T7/PstI |
| ***OdBrachyury*** | 5'..GGTTCGCACTGGATGAAACAGCC..3' | 5'..TATCCGTTCTGACACCAGTCGTTC..3' | gDNA exon 3; 630bp; PCR4-TOPO; T3/NotI |
| ***OdDach*** | 5'..GAGATGGATCCTGCGCAGC..3' | 5'..GTAAGTTTAAGAATATCCGAAAATCC..3' | cDNA full lenght; 770bp; PCR4-TOPO; T7/SpeI |
| ***OdEbf (COE)*** | 5'..GAGATCATGTGTTCCCGATGTTG..3' | 5'..GTTGAGTGAAAGAAAACCTTGCT..3' | cDNA exon 5 to exon 8; 578bp; PCR4-TOPO; T3/NotI |
| ***OdERK*** | 5'..GAAGGAGCCTACGGCATAG..3' | 5'..GCTAGAATACATCCGACAGAC..3' | cDNA exon 1 to exon 5; 563bp; PCR4-TOPO; T3/NotI |
| ***OdEts1/2b1*** | 5'..GACGGCATTGATGGATTACAGCTAT..3' | 5'..GTTACTCTTCAGTCTCTGGCTC..3' | cDNA exon 3-4 to exon 7; 480bp; PCR4-TOPO; T3/NotI |
| ***OdEts1/2b2*** | 5'..GACTCTCTTTGCCATCAATCC..3' | 5'..GCCTTTTTTCGTCCAGCTAATG..3' | cDNA exon 2 to exon 5; 728bp; PCR4-TOPO;T3/NotI |
| ***OdFilamin C*** | 5'..GCTACGAGCCACAAGAGAAG..3' | 5'..GCGTATTCACCAGCCTCTGTT..3' | gDNA exon 7; 604bp; PCR4-TOPO; T7/PstI |
| ***OdFoxF*** | 5'..GGCTGGAAGAATTCCGTCCG..3' | 5'..GAGCTGATTCGCATGGGCAGG..3' | cDNA exon 3 to exon 7; 639bp; PCR4-TOPO; T7/SpeI |
| ***OdGata1/2/3b*** | 5'..GCCTCTTCTGATTCGCCATTC..3' | 5'..GGAATGACTGTTTGGTGTTGG..3' | cDNA exon 1 to exon 3; 791bp; PCR4-TOPO; T3/XbaI |
| ***OdGata1/2/3d*** | 5'..GGGCAGAATATGAAAATGTATTTT..3' | 5'..GCTGACCGTCCGCTAGTC..3' | cDNA full lenght; 1086bp; PCR4-TOPO; T7/BamHI |
| ***OdHand1/2*** | 5'..GATGGAGTTGAATTTGTATTCCGATC..3' | 5'..GATTCTTTTCTAATCAGATGGGCA..3' | cDNA full lenght; 462bp; PCR4-TOPO; T7/SpeI |
| ***OdIslet*** | 5'..GGTCTCCGGTGATGAATTCCT..3' | 5'..GCTTTGTCGTAGGTTTAGGCCA..3' | cDNA exon 4 to exon 8; 766bp; PCR4-TOPO; T7/PstI |
| ***OdMath6*** | 5'..GCCGAATTCCACACAACGAAG..3' | 5'..CAACGTTAGCAGGTATAGAATAG..3' | cDNA exon 1-2 to 3'UTR; 602bp; PCR4-TOPO; T7/SpeI |
| ***OdMyosin*** | 5'..GACTCTTCAAGTGGCTCGTCG..3' | 5'..GGCCAAATCCAATCCGAAATCG..3' | gDNA exon 2; 254bp; PCR4-TOPO; T3/NotI |
| ***OdNk4*** | 5'..GACCGAAAAATTACAACTATGAGC..3' | 5'..GCTGTAGCGCCGAGCTCAC..3' | cDNA exon 1 to exon 3; 649bp; PCR4-TOPO; T3/NotI |
| ***OdRapostlin*** | 5'..GAAGATATGAGCCACTTGCCTC..3' | 5'..GTTGTGATTGAAGAAGTTAGTCCAG..3' | cDNA exon 6 to exon 8; 628bp; PCR4-TOPO; T3/NotI |
| ***OdTnnT1*** | 5'..GACGAAGAAGATCTTGATAGC..3' | 5'..GGATTCGAAGTTGCGTACTG..3' | gDNA exon 2; 218bp; PCR4-TOPO; T7/PstI |
| ***OdTnnT4*** | 5'..GAAATCGGATCTAAAAAGG..3' | 5'..GTCATTTTCACGCACGCGCA..3' | cDNA exon 1 to exon 5; 569bp; PCR4-TOPO; T3/NotI |
| ***OdTnnT7*** | 5'..GCACGTCATCCCTAGAATA..3' | 5'..GAAGCTTGCCAACTCGTGT..3' | cDNA 5'UTR to exon 5; 670bp;PCR4-TOPO; T7/PstI |
| ***OdMyoD*** | 5'..GTTACAAAATGACTATGACGGAAAC..3' | 5'..GGCTTCCAAGTTTCTTGACCAG..3' | cDNA full lenght; 813bp; PCR4-TOPO; T7/PstI |

**Supplementary Table 4.** Developmental effects of LDN and SU5402 at different concentrations and periods of incubations.

| Starting time incubation | Concentration | Hatch success | Correct early-hatchling |
|---|---|---|---|
| From 2-cell stage | DMSO 0,2% | 83% (n=231) | 60% (n=168) |
| | LDN 10 μM | 31% (n=72) | 1% (n=2) |
| | SU5402 50μM | 4% (n=12) | 0% (n=0) |
| From 32-cell stage | DMSO 0,2% | 86% (n=391) | 68% (n=312) |
| | LDN 10 μM | 55% (n=216) | 10% (n=34) |
| | SU5402 50 μM | 91% (n=240) | 60% (n=158) |
| | DMSO 0,3% | 94% (n=262) | 85% (n=236) |
| | SU5402 100 μM | 66% (n=116) | 25% (n=45) |
